# The nascent transcriptome delineates the regulatory landscape in human health and disease

**DOI:** 10.1101/2025.09.24.676871

**Authors:** Sagar R. Shah, You Chen, Alden K. Leung, Priscila V. Castilho Navarro, Mauricio I. Paramo, Juhi Gupta, Aishwarya Gurumurthy, Rebuma F. Fite, Annika K. Weimer, Kelly Cochran, Austin Wang, Qian Du, Ahmed M. Mohyeldin, Dieter Egli, Anshul Kundaje, Remi J. Creusot, Russell J.H. Ryan, Michael P. Snyder, Andrew G. Clark, John T. Lis, Haiyuan Yu

## Abstract

Transcriptional regulatory elements (TREs) orchestrate gene expression programs fundamental to cellular identity and transitions across physiological and pathological states. Here, we present a high-resolution atlas of RNA Polymerase II-engaged TREs (enhancers and promoters) across all major human organ systems and a broad spectrum of developmental and disease states. This atlas is generated using PRO-cap, a highly sensitive method that detects nascent RNA at transcription initiation sites, a critical feature of active TREs. The base-pair resolution of PRO-cap enables systematic dissection of transcription initiation architecture, revealing associations among tissue specificity, evolutionary constraint, transcription factor usage, and regulatory connectivity. Integration with deep learning models such as ProCapNet further provides a framework for prioritizing noncoding variants from GWAS and eQTL studies. Moreover, this tissue-resolved atlas identifies lineage-specific regulatory programs and their alterations in diseases such as metastatic cancer, where TRE landscapes capture regulatory signatures reflecting both tissue of origin and adaptive responses to distant niches. Together, these findings establish transcription initiation at regulatory elements as a defining and mechanistically informative layer of gene regulation across development, physiology, and disease.

## Introduction

Transcriptional regulatory elements (TREs), including enhancers and promoters, shape gene expression programs that govern development, maintain cellular homeostasis, and drive disease progression. Traditionally, TREs have been defined using epigenomic signatures, such as chromatin accessibility and histone modifications, as implemented in resources like the ENCODE candidate cis-regulatory element (cCRE) annotations^1,2^. In addition to these epigenomic features, genome-wide studies have revealed pervasive RNA polymerase II (Pol II) transcription initiation at TREs, with promoters producing stable transcripts and enhancers generating short-lived enhancer RNAs (eRNAs)^3–12^. These transcription events have emerged as a hallmark of active TREs, providing a direct indicator of regulatory activity that is not captured by chromatin features alone^13–15^. Therefore, systematically mapping transcription across diverse physiological and pathological states is essential for building a comprehensive catalog of active TREs and advancing our understanding of gene regulation in human health and disease.

Large-scale efforts such as the FANTOM consortium established transcription initiation as a powerful principle for regulatory element discovery, using Cap Analysis of Gene Expression sequencing (CAGE-seq) to map promoters and enhancers across diverse human tissue and cell types^9^. However, because CAGE-seq primarily captures steady-state RNAs and is influenced by RNA stability and processing, it has limited sensitivity for highly unstable eRNAs. Run-on–based approaches overcome this limitation by selectively enriching for Pol II-engaged nascent RNAs. In particular, Precision nuclear Run-On sequencing with 5′-capped RNA enrichment (PRO-cap)^16^ enables precise genome-wide mapping of transcription start sites (TSSs) and identification of active TREs, achieving optimal sensitivity and specificity when benchmarked across 13 RNA-sequencing assays^17^.

To enable large-scale mapping of active TREs, we developed a streamlined and optimized PRO-cap protocol compatible with diverse human biospecimens, ranging from cadaveric tissues to clinical biopsies from both solid and liquid organs. Using this approach, we constructed a comprehensive transcription-based atlas of regulatory element activity across human tissues and cell types spanning diverse physiological and pathological states. In this study, we define active TREs operationally based on PRO-cap–detected transcription initiation, thereby restricting our analysis to transcriptionally engaged elements and excluding inactive and poised regulatory regions that can be transcriptionally active at another time and/or space^18^. By resolving transcription initiation and pausing with base-pair precision, this atlas uncovers the architecture, evolutionary constraints, and regulatory syntax of TREs, illuminating how transcription encodes regulatory logic underlying human traits and diseases.

## Results

### A transcription-based atlas of regulatory elements across physiological and pathological contexts

To establish a high-confidence, transcription-based map of human regulatory elements, we first sought to develop a scalable, robust platform for profiling nascent RNA across diverse biospecimens. We optimized the PRO-cap protocol to markedly increase throughput and versatility while preserving the sensitivity and specificity of the original assay^19^ (Fig. 1a). The streamlined workflow compresses library preparation time from 4 days to approximately 10 hours and reduces input requirements to 1-2 million cells, enabling broad profiling beyond cell culture across primary solid and liquid tissues, clinical specimens, and disease models derived from cadaveric and surgical biopsies. The protocol incorporates on-bead processing steps and optimized adaptor usage to minimize sample loss and library artifacts, improving efficiency and reproducibility. Libraries generated using this workflow exhibited size distributions, alignment rates, and peak profiles highly concordant with those obtained using the original PRO-cap protocol, validating fidelity while substantially enhancing usability.

**Figure 1.**
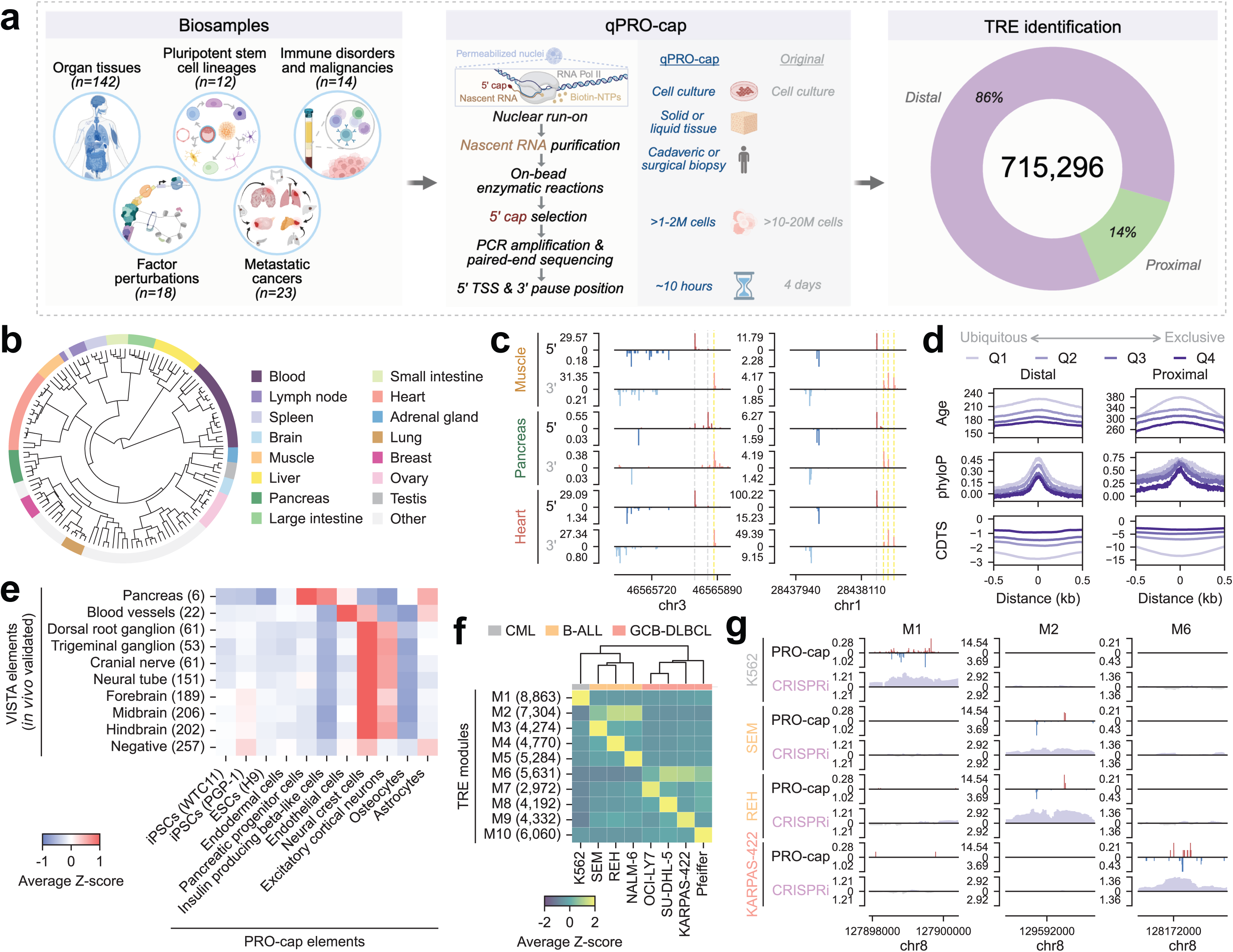
A transcription-based atlas of regulatory elements across human development, physiology, and disease. **(a)** Overview of the PRO-cap profiling strategy and resulting regulatory landscape. PRO-cap was applied across a broad collection of biosamples (n = 215) spanning all major human organ systems, pluripotent stem cells and their differentiated lineages, diverse disease states, and cells subjected to degron-mediated factor perturbation. The assay captures sites of transcription initiation and pausing via paired-end sequencing, enabling genome-wide identification of active TREs. Please see Supplementary Table 1 for all biosample and donor information for each PRO-cap library. **(b)** Dendrogram illustrating hierarchical clustering of tissue samples (n = 128) based on normalized PRO-cap expression levels of divergent distal TREs. Each leaf corresponds to a sample, and subtrees that primarily contain a single tissue type with at least three samples are highlighted. **(c)** Representative browser tracks of PRO-cap signals from three tissue types (muscle, pancreas, and heart) obtained from the same donor (ENCDO793LXB) across two distal genomic loci. Each track displays RPM-normalized PRO-cap reads mapped to the plus (red) and minus (blue) strands for each biosample. Darker shades indicate 5′ TSS positions, whereas lighter shades indicate 3′ pausing positions. Grey and yellow dashed lines highlight inter-tissue variations in 5′ and 3′ positions, respectively. **(d)** Metaplots of sequence age (million years ago), phyloP, and CDTS across distal (n=248,200) and proximal (n=61,184) divergent TREs grouped by specificity score quantiles, with Q1 indicating the most ubiquitously expressed and Q4 the most tissue/cell type-specific. Distances are shown as ±0.5 kb from the element center. **(e)** Heatmap showing the average Z-scores of PRO-cap signals at divergent TREs overlapping VISTA elements tested in transgenic mouse embryos. VISTA categories include enhancers validated in a given tissue (e.g., pancreas, regardless of activity in other tissues) and negative controls (elements not validated as enhancers in any tested tissue). Numbers in parentheses indicate the number of PRO-cap–detected elements overlapping VISTA elements within each category. **(f)** Hierarchically clustered heatmap of leukemia and lymphoma subtypes (CML, B-ALL, and GCB-DLBCL) based on divergent distal TRE expression patterns. The heatmap shows average Z-scores of elements within each TRE module (M1-M10) across samples. The number in parentheses indicates the number of elements in each TRE module. **(g)** Genome browser tracks of 5′ PRO-cap signal (red, plus strand; blue, minus strand) and CRISPRi scores (grey) from tiling screens for three representative loci from TRE modules M1, M2, and M6 across four leukemia and lymphoma cell lines (K562, SEM, REH, and KARPAS-422). Each locus corresponds to a 2.5 kb region at the MYC locus on chromosome 8, shown across the different cell lines.

Leveraging this optimized approach, we generated a compendium of TREs by mapping TSSs and pause sites spanning all major human organ systems, pluripotent stem cells and their differentiated lineages, transcription-related factor perturbation models, and a broad spectrum of disease contexts, including type 1 diabetes, leukemia, lymphoma, and a variety of metastatic cancers (Fig. 1a, Extended Data Fig. 1a, and Supplementary Table 1). Across the full dataset (n = 215 biosamples), PRO-cap libraries have sufficient sequencing depth (median deduplicated reads = 29.6 million; Supplementary Fig. 1a) and high reproducibility, with most biosamples having at least two technical replicates (median correlation of genome-wide reads = 0.902; Supplementary Fig. 1b,c). We also examined gene body ratios, defined as the fraction of length-normalized PRO-cap signal in the gene body relative to the sum of length-normalized signal at the TSS region and in the gene body, for highly expressed genes (see Methods for details). Higher gene body ratios indicate potential contamination from genic transcription derived from non-degraded 5′ monophosphates, likely due to incomplete enzymatic digestion during library preparation rather than true TRE-associated capped RNAs. Our libraries showed low gene body ratios, reflecting enrichment of signals at TSSs relative to gene bodies (Supplementary Fig. 1d). In addition, the median RNA length was 31bp, consistent with 5′-capped nascent RNA fragments from paused Pol II (Supplementary Fig. 1e). Together, these metrics demonstrate the high quality of the dataset, providing a reliable foundation for downstream analyses.

Across this dataset, we identified a union set of 715,296 TREs using the peak-calling tool Peak Identifier for Nascent Transcript Starts (PINTS)^17^. TREs were classified as proximal or distal based on their distance to annotated gene TSSs, corresponding to putative promoters and proximal enhancers or distal enhancers, respectively (Fig. 1a and Extended Data Fig. 1b). Distal TREs were further grouped by their inter- or intragenic location. Within each biosample, TREs were categorized by transcription initiation patterns as either divergent, defined by a pair of peaks on opposite DNA strands, or unidirectional, with peaks detected on a single strand (see Methods). Within the distal intragenic group, ∼15% of TREs localized to exonic regions (Extended Data Fig. 1c), a class of regulatory elements recently shown to exhibit enhancer activity within coding exons^20^. Genome-wide mapping revealed that both divergent and unidirectional TREs are broadly distributed and preferentially enriched in gene-dense regions (Supplementary Fig. 2), supporting a pervasive role for transcription initiation in organizing regulatory landscapes. The union set of TREs was generated by merging elements across all biosamples. Each element was classified as divergent if it was identified as divergent in any sample, and as unidirectional only if it was consistently unidirectional in every sample in which it appeared. Given that sequencing depth and technical variation can lead to misclassification of divergent TREs as unidirectional^21^, we restricted downstream analyses to divergent TREs to ensure stringency, while noting that the comprehensive catalog of unidirectional TREs provided here represents a complementary resource for future studies of regulatory diversity and transcriptional directionality.

### PRO-cap recapitulates tissue-specific regulatory logic

To further demonstrate the quality of this atlas, we next asked whether divergent TREs identified by PRO-cap follow established principles of tissue-specific gene regulation. Hierarchical clustering of biosamples using normalized PRO-cap signal from divergent distal and proximal TREs revealed strong tissue-level organization, with functionally related tissues clustering together, independent of donor status (Fig. 1b, Supplementary Fig. 3). For example, cardiac and skeletal muscle form a unified striated muscle cluster, while lymphoid organs, such as spleen, lymph node, and appendix, group into a coherent immune cluster. To quantitatively assess tissue specificity underlying this organization, we computed a specificity score for each element (see Methods). Distal elements have higher specificity scores than proximal ones (Extended Data Fig. 2a), consistent with the well-established tissue specificity of enhancers. In addition, liver, brain, and testis exhibit higher proportions of tissue-restricted TREs (Extended Data Fig. 2b), in agreement with previous work^9^. Together, these results confirm that PRO-cap-identified TREs recapitulate known principles of tissue-specific regulatory architecture.

At the level of individual TREs, PRO-cap detected the variation in both transcription abundance and profiles. Some TREs are broadly active, whereas others are highly tissue-restricted, with activity confined to a single tissue or a small group of related tissues (Extended Data Fig. 2c). Zooming into anatomically adjacent regions, such as segments of the large intestine, revealed both shared and segment-specific TREs, highlighting the fine-grained resolution of our PRO-cap assay. Moreover, PRO-cap also revealed substantial variation in transcriptional profiles, with distinct 5′ initiation and 3′ pausing patterns for specific TREs when compared across different tissues from the same donor (Fig. 1c). These high-resolution patterns suggest that transcriptional profiles themselves encode a regulated and biologically informative dimension of TRE activity that is not captured by histone modifications and open chromatin measurements.

### Evolutionary forces shape tissue-specific regulatory programs

Previous studies have extensively characterized the tissue specificity of regulatory elements using epigenomic-based maps, but how evolutionary pressures shape transcriptionally active regulatory programs remains poorly understood. We therefore examined how evolutionary constraint relates to tissue-specific TRE usage. Stratifying divergent distal and proximal elements by specificity scores into four quantiles, we found that elements with the lowest tissue specificity (Q1) were older than the most tissue-specific elements (Q4), based on a syntenic block-based sequence age assignment^22^ (Fig. 1d). Q1 elements were also the most evolutionarily conserved across species, as measured by phyloP scores^23^, with the signal peaking at the element center, corresponding to upstream regions enriched for TF binding sites. In addition, Q1 elements were the least tolerant to variation in the human population, as assessed by context-dependent tolerance scores (CDTS)^24^. Collectively, these findings indicate that broadly active TREs are subject to stronger purifying selection, consistent with their involvement in core cellular programs.

Next, to examine selective constraints in tissue-specific programs, we identified the top 5% of divergent elements with the highest expression specificity in each tissue type (Extended Data Fig. 3a). These tissue-restricted TREs (both distal and proximal) exhibited distinct evolutionary patterns: brain-specific elements were under the strongest constraint, whereas liver- and testis-specific elements showed signatures of more rapid evolution (Extended Data Fig. 3b). These trends are concordant with findings in mouse^25^, suggesting conserved regulatory hierarchies and tissue-dependent turnover dynamics across mammals. Taken together, these analyses reveal that tissue specificity and evolutionary constraint are tightly intertwined at actively transcribed TREs.

### Functional validation of PRO-cap detected TREs across development and disease

To functionally validate our dataset, we examined whether PRO-cap-detected TREs exhibit cell- and tissue-specific enhancer function *in vivo*, across both developmental and disease contexts. To that end, we first examined their PRO-cap activity across human developmental lineages. Hierarchical clustering showed that iPSC- and ESC-derived cells closely resembled pluripotent stem cells rather than their corresponding adult tissues (Extended Data Fig. 4a), in line with prior findings^26^. Comparison with candidate enhancers tested in transgenic mouse embryos from the VISTA Enhancer Browser^27^ revealed strong concordance between PRO-cap transcription levels and *in vivo* enhancer activity. VISTA elements with positive enhancer activity in specific tissues exhibited significantly higher PRO-cap signals in matching human cell types. For example, VISTA-validated blood vessel enhancers had elevated PRO-cap signals in endothelial cells relative to other cell types, while those associated with nervous system structures (i.e., dorsal root and trigeminal ganglia, carnival nerve, neural tube, forebrain, midbrain, and hindbrain) were highly transcribed in neural crest and excitatory cortical neurons (Fig. 1e). Furthermore, to capture the dynamic remodeling of TRE usage during lineage progression, we profiled a time course of pancreatic differentiation, from ESCs to definitive endoderm, pancreatic progenitors, and beta-like cells. Divergent distal elements exhibited significant temporal variation and clustered into distinct transcriptional trajectories (Extended Data Fig. 4b). By linking these dynamically transcribed elements to their nearest expressed genes as putative targets and performing functional enrichment analysis, we identified stage-specific activation of biological pathways aligned with known differentiation dynamics. Specifically, PRO-cap signals at TREs associated with pluripotency declined over time, while those linked to endodermal and pancreatic lineage commitment emerged as differentiation progressed (e.g., receptor tyrosine kinase signaling in beta-like cells^28^).

Having shown that PRO-cap captures developmentally regulated TRE programs with strong concordance to *in vivo* enhancer activity, we next turned to pathological contexts, asking whether PRO-cap can resolve subtype-specific regulatory landscapes in hematologic malignancies. Therefore, we performed PRO-cap on a panel of leukemia and lymphoma models comprising one chronic myelogenous leukemia (CML) cell line (K562), three B-cell acute lymphoblastic leukemia (B-ALL) cell lines (SEM, REH, and NALM-6) and four germinal center B-cell-like diffuse large B-cell lymphoma (GCB-DLBCL) lines (OCI-LY7, SU-DHL-5, KARPAS-422, and Pfeiffer), each harboring distinct cytogenetic features, including oncogenic fusion proteins^29–34^ and aberrant activation of the *MYC* 3′ enhancer^35^. Hierarchical clustering separated these hematologic malignancies according to lineage and differentiation stage (Fig. 1f), corroborating their known developmental origins. To delineate coordinated regulatory programs, we applied k-means clustering and identified ten TRE modules with subtype-specific activity patterns (Fig. 1f). While M2 and M6 were shared across all B-ALL and GCB-DLBCL lines, respectively, the other modules showed higher activity in CML (K562) or in individual B-ALL or GCB-DLBCL lines. *De novo* motif enrichment analysis identified TF motifs specific to each module (Extended Data Fig. 5a and Supplementary Table 2), reflecting their distinct mechanisms of regulatory dysfunction. For instance, M4, which is specific to REH cells carrying the ETV6-RUNX1 fusion, was enriched for GGAA repeats. This is consistent with recent work showing that loss of ETV6-mediated repression enables GGAA microsatellite repeats to function as active enhancers driving aberrant gene expression in ETV6-deficient B-ALL^36^. On the other hand, modules M8 and M9, specific to SU-DHL-5 and KARPAS-422, respectively, were enriched for POU2F motifs. Both cell lines exhibit dependence on a recently identified *MYC* 3′ enhancer regulated by a transcriptional triad (POU2F2, MEF2B, and POU2AF1) and show elevated *POU2F2* expression relative to other GCB-DLBCL lines^35^. Finally, we assessed the functional significance of these modules in a subtype-specific context. We performed CRISPR interference (CRISPRi) tiling screens at the *MYC* locus across five representative cell lines (K562, REH, SEM, NALM-6, and KARPAS-422), enabling systematic and unbiased perturbation of both PRO-cap–defined TREs and regions lacking PRO-cap signal as internal controls for specificity. For SU-DHL-5, we leveraged a previously published targeted nucleosome-free region screen^35^. Silencing of module-specific TREs (e.g. M2) led to a pronounced reduction in *MYC* expression in the corresponding cell lines (e.g., SEM, REH, and NALM-6), with minimal effects in other subtypes where these regions lack PRO-cap signal (Fig. 1g and Extended Data Fig. 5b,c). These results confirm the selective regulatory function of these TRE modules in distinct malignant contexts. Collectively, these analyses establish that PRO-cap–detected TREs are functionally relevant across development and disease, further demonstrating the quality and biological fidelity of our dataset.

### PRO-cap-detected TREs benchmarked against existing annotations

We next sought to position our PRO-cap-defined atlas within the landscape of existing regulatory annotations. For each tissue type (e.g., the lower lobe of left lung), elements were merged across biological replicates within each assay prior to overlap analysis (Supplementary Table 3). Then, we performed systematic benchmarking of our TREs against tissue-matched epigenomic and transcriptomic datasets. Most PRO-cap-detected TREs, both distal and proximal, overlapped with peaks defined by chromatin accessibility (i.e., ATAC-seq and DNase-seq) or histone marks (i.e., H3K27ac ChIP-seq), as well as ENCODE cCRE annotations that integrate multiple epigenomic features (Fig. 2a top, Supplementary Table 3). Notably, while proximal TREs showed substantial agreement with CAGE-seq-based annotations from the FANTOM consortium, the vast majority of distal TREs captured by PRO-cap were missed by CAGE-seq, which primarily detects stable RNAs. This underscores the high sensitivity of PRO-cap in capturing short-lived, unstable eRNAs, providing a more complete view of active enhancers that are underrepresented in mature RNA datasets.

**Figure 2.**
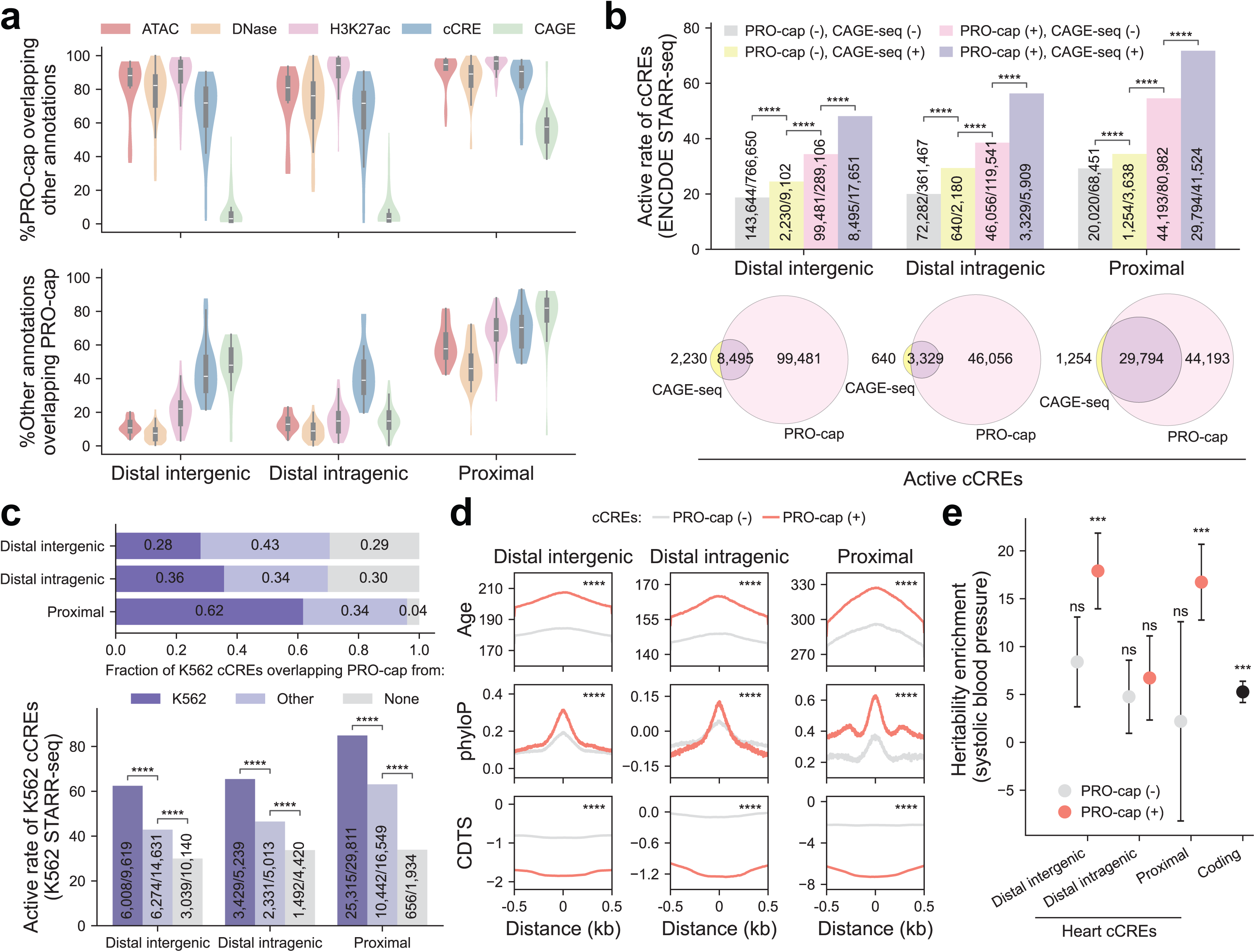
Transcription measured by PRO-cap is a critical mark for active TREs. **(a)** *Top:* Violin plots showing the percent of PRO-cap TREs overlapping ATAC-seq, DNase-seq, H3K27ac ChIP-seq, cCREs, and CAGE-seq peaks, stratified by TRE category. Each datapoint represents the percent overlap between two datasets for a given matched tissue type. *Bottom:* Violin plots showing the percent of elements annotated by ATAC-seq, DNase-seq, H3K27ac ChIP-seq, cCREs, or CAGE-seq that overlap with PRO-cap TREs. Each datapoint represents the percent overlap between two assays for a given matched tissue type (47, 49, 34, 29, and 25 tissue types compared between PRO-cap versus ATAC-seq, DNase-seq, H3K27ac ChIP-seq, cCREs, and CAGE-seq, respectively). **(b)** *Top:* Active rate (%) of distal intergenic, distal intragenic, and proximal cCREs, stratified by their overlaps with PRO-cap and/or CAGE-seq elements. *Bottom:* Venn diagram showing the number of STARR-seq–active cCREs overlapping PRO-cap and/or CAGE-seq elements. **(c)** *Top:* Fraction of distal intergenic, distal intragenic, and proximal K562 cCREs overlapping PRO-cap elements from K562, other cells/tissues (but not in K562), or absent from our entire dataset. *Bottom:* Active rate (%) of these cCRE categories (same as the top panel) measured by K562 STARR-seq. **(d)** Metaplots of sequence age (million years ago), phyloP, and CDTS across distal intergenic, distal intragenic, and proximal cCREs, stratified by PRO-cap TRE overlap. Distances are shown ±0.5 kb from the element center. The same set of elements shown in (b) was used in this analysis, whereby *PRO-cap (−)* denotes *PRO-cap (−), CAGE-seq (−)* and *PRO-cap (−), CAGE-seq (+)* elements, and *PRO-cap (+)* includes *PRO-cap (+), CAGE-seq (−)* and *PRO-cap (+), CAGE-seq (+)* elements. **(e)** Heritability enrichment for a representative cardiac trait (systolic blood pressure) across coding regions and heart-derived cCREs, stratified by their overlaps with heart-derived PRO-cap elements. BH-adjusted p-values (padj) are displayed as follows: ns, non-significant; ***, 0.0001 < padj ≤ 0.001; ****, padj ≤ 0.0001.

In reciprocal analyses, we anchored on other annotations and quantified their overlap with PRO-cap-detected TREs. Proximal elements showed consistent agreement across different methods (Fig. 2a bottom, Supplementary Table 3). In contrast, a considerable number of distal elements defined by individual epigenomic markers alone (i.e., ATAC-seq, DNase-seq, or H3K27ac ChIP-seq) did not overlap with PRO-cap TREs, whereas cCREs, which combine these marks, show substantially greater concordance. These results demonstrate that a single transcription-based assay can effectively capture regulatory information otherwise inferred from multiple epigenomic features, providing a robust approach for detecting active distal TREs. In addition, examination of CAGE-seq-detected elements relative to PRO-cap revealed a higher proportion of overlap in proximal and distal intergenic regions compared with distal intragenic regions, potentially reflecting the previously reported prevalence of spurious capping events from exonic sequences rather than true transcription initiation sites in CAGE-seq datasets^37,38^. Notably, we applied a minimal 1-bp overlap criterion to maximize detection of shared regions in both analyses, whether anchored on PRO-cap elements or other annotations. Even under this highly permissive definition, substantial discrepancies remain for certain assay pairs, indicating that these differences are biologically meaningful rather than artifacts of stringent overlap thresholds.

### Transcription is a critical mark of active TREs

Our benchmarking analyses indicate that, although chromatin accessibility and histone modifications demarcate large numbers of candidate regulatory elements, only a subset exhibit active transcriptional initiation. This raises the question of whether transcription provides additional functional resolution beyond epigenomic annotations.

To address this, we leveraged recently published functional activity calls for cCREs derived from genome-wide STARR-seq (i.e., self-transcribing active regulatory region sequencing) reporter assays across six cell lines^2^. We categorized cCREs based on their overlap with PRO-cap and/or CAGE-seq annotations. Across distal and proximal regions, cCREs overlapping either transcription-based annotation exhibited higher active rates than those without such annotations (Fig. 2b top and Supplementary Table. 3). These results indicate that transcription is a reliable marker of functional regulatory activity. Notably, some cCREs lacking PRO-cap or CAGE-seq annotations still showed activity, likely reflecting limitations of peak-calling thresholds, as these regions may harbor low-level transcription that falls below detection in final annotation sets. Supporting this, PRO-cap expression has been shown to scale with enhancer activity, supporting the functional relevance of even low-level and possibly sub-threshold initiation events^14^.

Notably, cCREs overlapping both PRO-cap and CAGE-seq showed the highest functional activity, followed by those overlapping PRO-cap alone, whereas cCREs detected only by CAGE-seq displayed more modest enrichment (Fig. 2b top). Among active cCREs, a greater number overlapped uniquely with PRO-cap TREs than with CAGE-seq annotations (Fig. 2b bottom and Supplementary Table. 3). Taken together, these results demonstrate that PRO-cap provides a more sensitive measure of transcription initiation and strongly predicts functional enhancer activity.

Some cCREs lack PRO-cap annotations yet a subset shows activity in STARR-seq assays (Fig. 2b top), raising the possibility that these elements represent poised enhancers. To investigate this, we categorized K562-derived cCREs based on their overlap with PRO-cap TREs in K562, in other cell or tissue types (but not in K562), or absent from our entire dataset (Fig. 2c top). As expected, cCREs overlapping PRO-cap TREs in the matched cell type (i.e., K562) showed the highest active rates, followed by those detected only in other cell or tissue types, whereas those not overlapping PRO-cap annotations across all biosamples exhibited the lowest rates (Fig. 2c bottom). These patterns are consistent with previous studies that show episomal assays likely capture intrinsic enhancer potential but do not fully reflect *in vivo* active cell-type–specific regulation^39,40^.

To further evaluate the functional relevance of transcribed cCREs, we examined their evolutionary conservation across the entire dataset. Strikingly, PRO-cap(+) cCREs across all genomic categories are significantly older and under stronger evolutionary constraints compared to PRO-cap(-) ones (Fig. 2d). Lastly, we asked whether PRO-cap(+) cCREs are preferentially enriched for trait/disease-associated variants. Using stratified linkage disequilibrium score regression (S-LDSC)^41,42^ to analyze genome-wide association study (GWAS) data of a representative cardiac phenotype, systolic blood pressure, we observed significantly greater heritability enrichment within PRO-cap(+) cCREs detected in heart samples (Fig. 2e). As expected, coding regions were enriched; however, PRO-cap(+) distal intergenic and proximal non-coding elements showed even greater enrichment.

Together, these results support transcription initiation as a robust marker of functionally active regulatory elements and position PRO-cap as a powerful approach for prioritizing biologically and clinically relevant regions. More broadly, they define transcription at TREs as a fundamental regulatory layer that complements and refines epigenomic annotations.

### Tissue-specific effects of disease- and trait-associated variants

Most variants identified by GWAS reside in noncoding regions, potentially altering TRE activity and perturbing their target gene expression in a context-dependent manner^6,43–46^. Building on our observation that PRO-cap(+) cCREs are enriched for disease-associated variants in a cardiac context (Fig. 2e), we extended this analysis to ask whether PRO-cap–defined elements capture the tissue- and cell-type contexts of genetic effects more broadly. Identifying the relevant biological setting in which these effects occur is essential for understanding disease mechanisms and informing precision medicine.

To evaluate the contribution of tissue-specific TRE annotation to trait heritability, we applied S-LDSC to 176 GWAS summary statistics spanning various human diseases and complex traits (Fig. 3a and Supplementary Fig. 4). Patterns of enrichment were highly tissue-specific, recapitulating known biological processes. For example, immune-related phenotypes exhibited significant enrichment in blood and lymphoid tissues, while neuropsychiatric diseases and many traits, including intelligence, educational attainment, body mass index, and smoking status, showed pronounced enrichment in brain tissues. Likewise, liver-specific enrichment was observed for biochemical traits (e.g., levels of bilirubin, low-density lipoprotein, and sex hormone-binding globulin), consistent with the liver’s roles in metabolism, transport, and detoxification.

**Figure 3.**
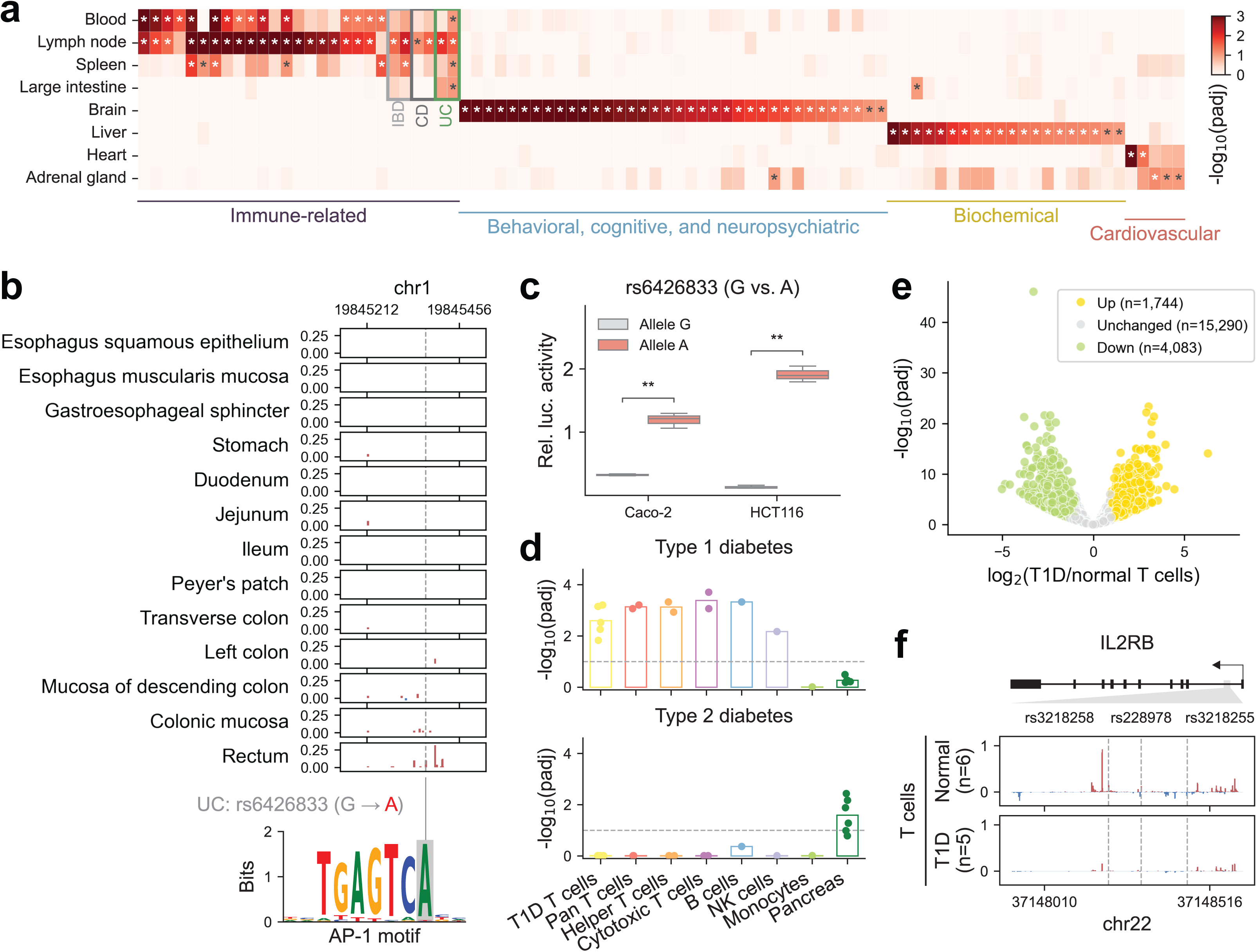
Transcriptional regulatory elements capture tissue-specific effects of trait- and disease-associated variants. **(a)** Heatmap showing the significance of enrichment for representative human diseases and complex traits across divergent distal tissue-specific TRE annotations. Asterisks (*) denote BH-adjusted p-values < 0.1 after Benjamini-Hochberg correction. The full panel is shown in Supplementary Fig. 4. IBD: Inflammatory bowel disease; CD: Crohn’s disease; UC: Ulcerative colitis. **(b)** Genome browser view of 5’ PRO-cap signal at a distal element across different gastrointestinal segments, harboring a fine-mapped GWAS variant (rs6426833) that alters an AP-1 motif. **(c)** Luciferase reporter assay showing enhancer activity for the risk allele (A) compared to the reference allele (G) in Caco-2 and HCT116 cells. Three biological replicates were performed for each allele in each cell line. **, 0.001 < padj ≤ 0.01. See full panel with positive and negative controls in Suppl. Fig 5. **(d)** The significance of enrichment for T1D (*upper panel*) and T2D (*lower panel*) across healthy immune cell types and pancreas, as well as T cells isolated from T1D patients. Each point represents an individual sample. The dashed line denotes the BH-adjusted p-value threshold of 0.1. **(e)** Volcano plot showing differential PRO-cap signal at TREs of T cells from T1D patients (n=5) versus non-diseased donors (n=6), with each point representing an element. TREs with significantly increased (yellow) or decreased (green) PRO-cap expression in T1D, defined aslog2FC| > 1 and BH-adjusted p-value < 0.05, are highlighted. **(f)** Browser shot of average PRO-cap signal tracks (RPM-normalized) at the *IL2RB* intronic enhancer locus in T cells from non-diseased donors and T1D patients. The *IL2RB* gene is depicted with an arrow indicating transcription orientation and black rectangles marking exons. The grey triangle highlights the region shown in the zoomed-in panels. Three fine-mapped T1D-associated variants (rs3218258, rs228978, and rs3218255) located within this locus are denoted by dashed lines.

Beyond recapitulating known biology at the organ level, these enrichment profiles enabled discrimination of disease subtypes with overlapping etiologies. For instance, inflammatory bowel disease (IBD) includes ulcerative colitis (UC) and Crohn’s disease (CD), each with distinct clinical features, including differences in the location of inflammation. We found that both UC- and CD showed enrichment in immune-related tissues; however, only UC displayed significant enrichment in the intestines, in line with its canonical disease manifestation^26,47,48^ (Fig. 3a). This tissue-specific signal was further supported at the variant level: a fine-mapped variant, rs6426833, associated with UC but not CD^49^, resides in a TRE with pronounced expression in the rectum (Fig. 3b), consistent with the clinical observation that UC inflammation typically originates in the rectum, with variable proximal extension^48^. Motif analysis predicted that the alternative allele (A) enhances binding of AP-1, a TF involved in the inflammatory response in UC^50,51^. To test this, we cloned the wild-type and mutant TREs into a luciferase reporter vector and measured enhancer activity in colorectal cancer cell lines Caco-2 and HCT116. The alternate allele consistently drove higher luciferase expression than the reference allele, confirming that this variant enhances regulatory activity in a large intestine-relevant cellular context (Fig. 3c and Supplementary Fig. 5a).

These tissue-level patterns underscore the importance of cellular context in mediating disease risk. In particular, autoimmune disorders are driven by dysregulation within specific immune cell populations. To further refine genetic resolution beyond whole tissues, we leveraged cell-type-resolved TRE annotations across a diverse panel of blood-derived immune cell types. This analysis revealed that trait-associated variants are consistently enriched in the most biologically relevant immune cell populations. Specifically, significant enrichment for white blood cell count phenotype is observed across all leukocyte cell types, whereas enrichment for specific count traits (i.e., monocyte, lymphocyte) is largely restricted to their corresponding cell types (Extended Data Fig. 6a). In addition, analysis of IBD and its subtypes, UC and CD, revealed broad and pronounced enrichment across all major immune cell populations (Extended Data Fig. 6b). This pervasive enrichment pattern reinforces the complex and multifaceted immune dysregulation that characterizes IBD pathogenesis^52^.

Furthermore, our analyses of type 1 and type 2 diabetes (T1D and T2D, respectively) highlighted their distinct pathogenic mechanisms. T1D, driven by autoimmune dysfunction, exhibited strong enrichment in T cell-associated TREs, as well as in B and natural killer (NK) cells, but minimal enrichment in monocytes (Fig. 3d and Extended Data Fig. 6c). In contrast, T2D, a metabolic disease, showed its strongest enrichment in pancreatic tissue, but not in any of the immune cell types. To directly interrogate disease-relevant regulatory activity, we generated PRO-cap libraries from T cells isolated from patients with T1D, enabling direct interrogation of disease-relevant regulatory activity. We observed significant enrichment of T1D, but not T2D, among TREs active in patient-derived T cells (Fig. 3d). We next identified a set of differentially expressed TREs (|log2FC| > 1 and adjusted p-value < 0.05) between T cells from T1D patients and non-diseased donors using DESeq2^53^ (Fig. 3e). Motif enrichment analysis of these TREs revealed significant enrichment for binding sites of TFs central to T cell function and autoimmune pathogenesis, including members of RFX, FOX, SMAD, and AP-1 families (Extended Data Fig. 7a and Supplementary Table 2). Differentially expressed TREs were linked to the nearest expressed genes, and pathway enrichment analysis demonstrated that these putative target genes were associated with altered cell surface receptor signaling (Extended Data Fig. 7b). Together, these results recapitulate established patterns of T cell transcriptional dysregulation in T1D, reinforcing that disease-associated TRE activity reflects canonical immune regulatory circuitry.

We next examined whether these differentially expressed TREs contain variants associated with T1D. Notably, an intronic enhancer locus within the *IL2RB* (interleukin-2 receptor beta) gene, harboring several potential causal variants^54^, exhibited robust transcriptional activity in healthy donor T cells but showed significantly reduced activity in T1D patient-derived T cells (log2FC=-1.43; adjusted P-value=4.02e-4; Fig. 3f and Extended Data Fig. 7c). In addition, we observed reduced PRO-cap signal at the *IL2RB* gene promoter in T1D samples (log2FC=-0.52; adjusted P-value=1.76e-2; Supplementary Fig. 5b), consistent with the decreased activity of the putative intragenic enhancers shown in Fig. 3f. Given the central role of IL2RB in T cell homeostasis, reduced promoter and intragenic TRE activity is expected to attenuate IL-2 receptor signaling. This links enhancer dysregulation to impaired cytokine signaling in T1D, with potential implications for disease progression and response to IL-2–based therapies.

Finally, to link these context-specific TREs to molecular regulatory effects, we evaluated their role in mediating genetic effects on gene expression across tissue types. To this end, we analyzed cis-expression quantitative trait loci (cis-eQTLs) from GTEx^55^ using TORUS^56^ and observed significant enrichment in distal elements from the corresponding tissue types (Extended Data Fig. 8a). In contrast, proximal elements showed broader enrichment across tissues (Extended Data Fig. 8b). As an orthogonal approach, we assessed whether fine-mapped eQTL variants, stratified by different posterior inclusion probability (PIP) thresholds, preferentially overlap tissue-specific distal TREs. Bayesian fine-mapping prioritizes likely causal variants within associated loci by assigning each variant a PIP, which reflects its probability of being causal. For a given PIP threshold, we calculated the percentage of fine-mapped variants from various GTEx tissues (curves) that overlap TREs of a given PRO-cap tissue type (panels) (Extended Data Fig. 8c). We observed that this percentage increases with more stringent PIP thresholds, particularly when the GTEx tissue matches the PRO-cap tissue. Interestingly, non-tissue-specific elements were pervasively enriched for eQTLs across tissues but showed little enrichment for specific GWAS phenotypes (Extended Data Fig. 8c and Supplementary Fig. 4).

Collectively, these results indicate that tissue- and cell type-resolved PRO-cap annotations enable systematic interpretation of disease- and trait-associated variants across both organismal (GWAS) and molecular (QTL) phenotypes.

### Sequence determinants of transcription initiation and context-dependent effects of regulatory variants

Given that transcription at noncoding regulatory regions confers functional relevance, we sought to elucidate the sequence features underlying transcription initiation profiled by PRO-cap. Understanding these features is critical for linking sequence, transcription, and context-dependent effects of genetic variants. To this end, we first trained ProCapNet^57^, a convolutional neural network that models base-resolution profiles of transcription initiation from local DNA sequence, across a panel of diverse tissue and cell types (n=16) (Fig. 4a). ProCapNet was trained on PRO-cap peaks, with accessible regions (DNase-seq or ATAC-seq peaks in matched tissues or cell types; Supplementary Table 3) that lack transcription initiation as background, enabling the model to learn sequence determinants of transcription initiation independent of chromatin accessibility. Overall, ProCapNet accurately predicted both total initiation activity (count task) and TSS positioning (profile task). Prediction performance on held-out chromosomes in the cross-validation test sets showed a strong correlation between observed and predicted counts at individual peaks (Supplementary Fig. 6a,b). Predicted PRO-cap profiles were also more similar to observed data (lower Jensen-Shannon distance indicates higher similarity) than baseline predictions corresponding to average profiles across peaks (Supplementary Fig. 6c,d). The gap between model performance and replicate concordance (Supplementary Fig. 6d) is expected, because ProCapNet is designed to model only the impact of local sequence context and therefore cannot capture the effects of distal regulatory elements and other non–sequence-based factors.

**Figure 4.**
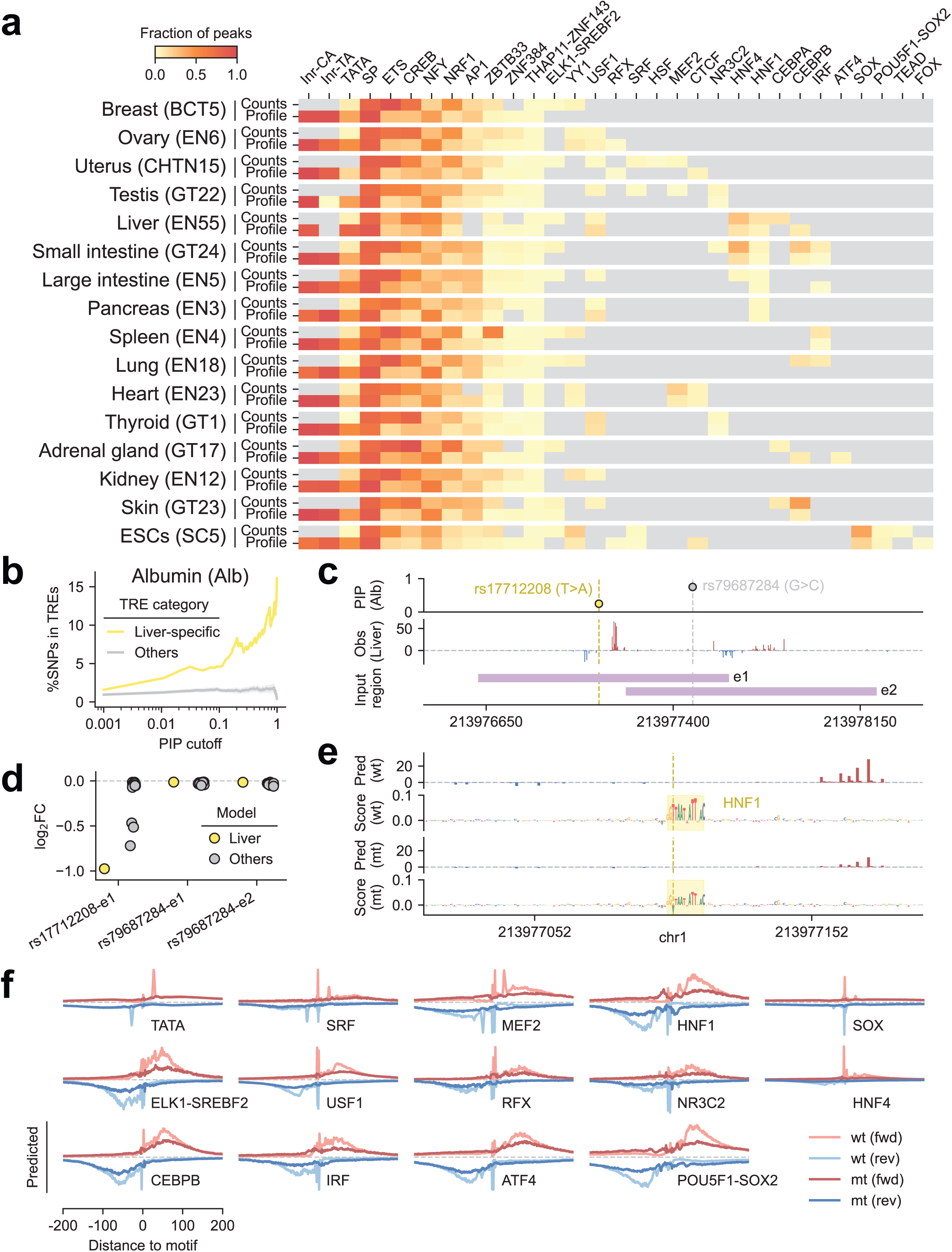
Tissue-specific modeling of transcription initiation delineates motif syntax and regulatory variant effects. **(a)** Fraction of PRO-cap peaks containing TF motifs identified by ProCapNet as contributing to transcriptional strength (count task) and/or transcription start site positioning (profile task) across multiple tissue and cell types (n=16). **(b)** Fraction of fine-mapped GWAS variants for albumin measurement (above a given PIP threshold) overlapping divergent distal elements from liver-specific (yellow) and other TRE (gray) categories. **(c)** The first track shows PIP values for two GWAS variants in a credible set associated with albumin measurement. The second track presents the PRO-cap signal observed in the liver sample (EN55). The final track displays the 1-kb input regions centered on each PRO-cap element (e1 and e2), with different variants used for prediction. **(d)** Predicted impact of variants rs17712208 and rs79687284 on transcription levels of PRO-cap elements e1 and e2, measured by log2 fold change between alternative and reference alleles in liver (yellow) and other tissue (gray) models. **(e)** Predicted transcription profiles and contribution scores (count task) for element e1 with either the reference or alternative allele of rs17712208 (yellow dashed line) using the liver-trained model. **(f)** The positional effects of representative TF motifs on transcription initiation profiles before and after *in silico* motif deletion. Median predicted profiles (profile task) are centered at motif instances and aligned by motif orientation, with reverse-strand motifs flipped. Forward and reverse strands are shown in red and blue, respectively; lighter and darker shades indicate wild-type and mutant profiles. The range of predicted values is shown in Supplementary Fig. 8.

To decipher the cis-regulatory sequence features captured by ProCapNet, we used DeepSHAP^58^, a well-established model interpretation framework, to estimate the contribution of each nucleotide within PRO-cap peaks to the count and profile prediction tasks separately. TF-MoDISco^59^ was then employed to cluster subsequences with high contribution scores across all PRO-cap peaks into a lexicon of motif patterns. The resulting motifs exhibited differential contributions across the two tasks and across tissues and cell types. Initiators (Inr-CA, Inr-TA) and the TATA box more frequently influenced TSS positioning (profile task), whereas ETS, CREB, and NRF1 predominantly affected transcription levels (count task), consistent with previous observations^57,60,61^. When comparing across models, some TF motifs (e.g., SP, ETS, CREB) were ubiquitously present, while others (e.g., SRF, MEF2, HNF1, TEAD) were more restricted to certain tissue and cell types, reflecting their potential roles in shaping enhancer logic within tissue-specific transcriptional programs. To assess whether these tissue-restricted motifs are associated with expression of their cognate transcription factors, we examined median gene expression levels (TPM) per tissue from GTEx for the corresponding TFs (Supplementary Fig. 7). We did not observe a consistent correspondence between motif restriction and TF expression levels, suggesting that tissue specificity may be partly encoded in the cis-regulatory syntax of broadly expressed TFs, rather than being driven solely by differences in TF expression levels.

Given that GWAS variants exhibit tissue-specific effects (Fig. 3a) and that our ProCapNet models capture how DNA sequence drives transcription initiation across diverse tissues and cell types (Fig. 4a), we applied these models to assess context-dependent effects of regulatory variants. As an illustrative example, we focused on GWAS data for serum albumin measurements and considered the liver as the biologically relevant tissue. The percentage of fine-mapped variants overlapping liver-specific TREs increases sharply with more stringent PIP cutoffs (more likely to be causal), whereas overlap with other TRE categories (i.e., tissue-specific TREs in non-liver tissues and non-tissue-specific TREs) remains low, consistent with tissue-specific regulatory effects (Fig. 4b).

Statistical fine-mapping typically defines 95% credible set as the minimal set of variants whose cumulative PIP exceeds 0.95 at a given locus. Because credible sets often contain multiple variants, we applied ProCapNet to predict the functional impact of each variant, enabling prioritization using both statistical evidence and predicted regulatory effects. Specifically, we examined one 95% credible set containing two variants associated with albumin measurements (Fig. 4c). Although the variant rs79687284 has a higher PIP of 0.750, *in silico* mutagenesis suggested minimal impact on transcription at either overlapping TREs (e1 or e2) across all tissue models (Fig. 4d). By contrast, rs17712208, which has a lower PIP of 0.249 and is located within a liver-specific TRE (e1), was predicted to disrupt an HNF1 motif and reduce transcription at TRE e1 by the liver-trained model (Fig. 4e). Importantly, this effect was strongest in the liver model and largely absent in models trained on other tissues (Fig. 4d), demonstrating its context-dependent regulatory impact. Therefore, our results indicate that integrating nascent RNA-based regulatory maps with statistical fine-mapping and tissue-specific deep learning models can enable prioritization and functional interpretation of noncoding variants. Although we illustrate this framework using serum albumin measurements and liver, it is broadly applicable across other tissues in our dataset and to diverse disease- or trait-associated variants.

### TF motifs exhibit distinct positional effects on transcription at TREs

Having identified a panel of TF motifs that contribute to transcriptional output and context-specific variant effects, we next investigated how these motifs also encode the positional logic of transcription initiation at TREs. While previous studies have examined positional effects of TF motifs on transcription in a limited set of cell lines using deep learning models^57,62,63^, others have experimentally dissected these rules for a different set of motifs^21,61,64^. Here, we extend these analyses to a broader panel of tissues and previously uncharacterized motifs. We first examined strand-specific PRO-cap signals centered on predictive motif instances, accounting for motif orientation, and observed high concordance between predicted and observed profiles (Supplementary Fig. 8). Among previously uncharacterized TF motifs, we identified two categories: one group (e.g., CEBPB, IRF, POU5F1–SOX2) was associated with broad effects on transcription initiation, whereas the other showed stronger positional effects (e.g., SRF, MEF2), reminiscent of the TATA box. These patterns were further supported by *in silico* ablation of motif instances (Fig. 4f).

Given the sequence similarity between the SRF motif and the TATA box, we leveraged available ChIP-seq datasets for SRF and TATA-binding protein (TBP) in human ESCs to confirm that the observed effects are attributable to SRF rather than misclassification of TATA-box. First, we found that a significantly higher proportion of SRF motifs overlapped SRF ChIP-seq peaks compared to TATA motifs (Supplementary Fig. 9a). Next, we restricted the analyses to SRF motif instances with direct ChIP-seq support and recapitulated the previously observed effect curves (Fig. 4f and Supplementary Fig. 8), characterized by a sharp spike at a fixed position from the motif, embedded within a broader peak for a given strand (Supplementary Fig. 9b). Additionally, ChIP-seq signals were symmetric and aligned with their respective motifs, with SRF and TBP signals peaking at SRF and TATA motifs, respectively (Supplementary Fig. 9c-e). Notably, at SRF motifs overlapping both SRF and TBP ChIP-seq peaks, TBP signals were offset from the motif center rather than aligned, indicating that these sites do not represent misclassified TATA motifs (Supplementary Fig. 9c,e). This overlap of TBP ChIP-seq peaks at SRF motifs is consistent with the known ability of TBP to be recruited to TATA-less regulatory elements via TBP-associate factors, rather than through direct motif binding.

Collectively, these results provide proof-of-principle for dissecting TF-specific positional effects on transcription initiation at TREs across our nascent RNA-based regulatory atlas. By integrating base-resolution TRE maps with deep learning-based motif identification and perturbation, this framework enables genome-wide, cross-tissue analyses and systematic prioritization of trait- and disease-associated variants.

### Transcription initiation and pausing patterns reveal mechanistic insights and biological significance of TRE architectures

While most TF motifs exhibited broad positional contribution profiles, a few displayed sharply localized effects on transcription initiation. We hypothesized that such motif-level differences may influence TSS positioning at TREs and potentially underlie biologically meaningful variation in regulatory function and evolutionary constraint. Therefore, we characterized initiation profiles by calculating the shape index (SI)^65,66^ for each strand of divergent TREs. Based on SI values, each strand of a divergent element was classified as Peaked (P) or Broad (B), resulting in three categories: P–P, P–B, or B–B (Fig. 5a and Supplementary Fig. 10).

**Figure 5.**
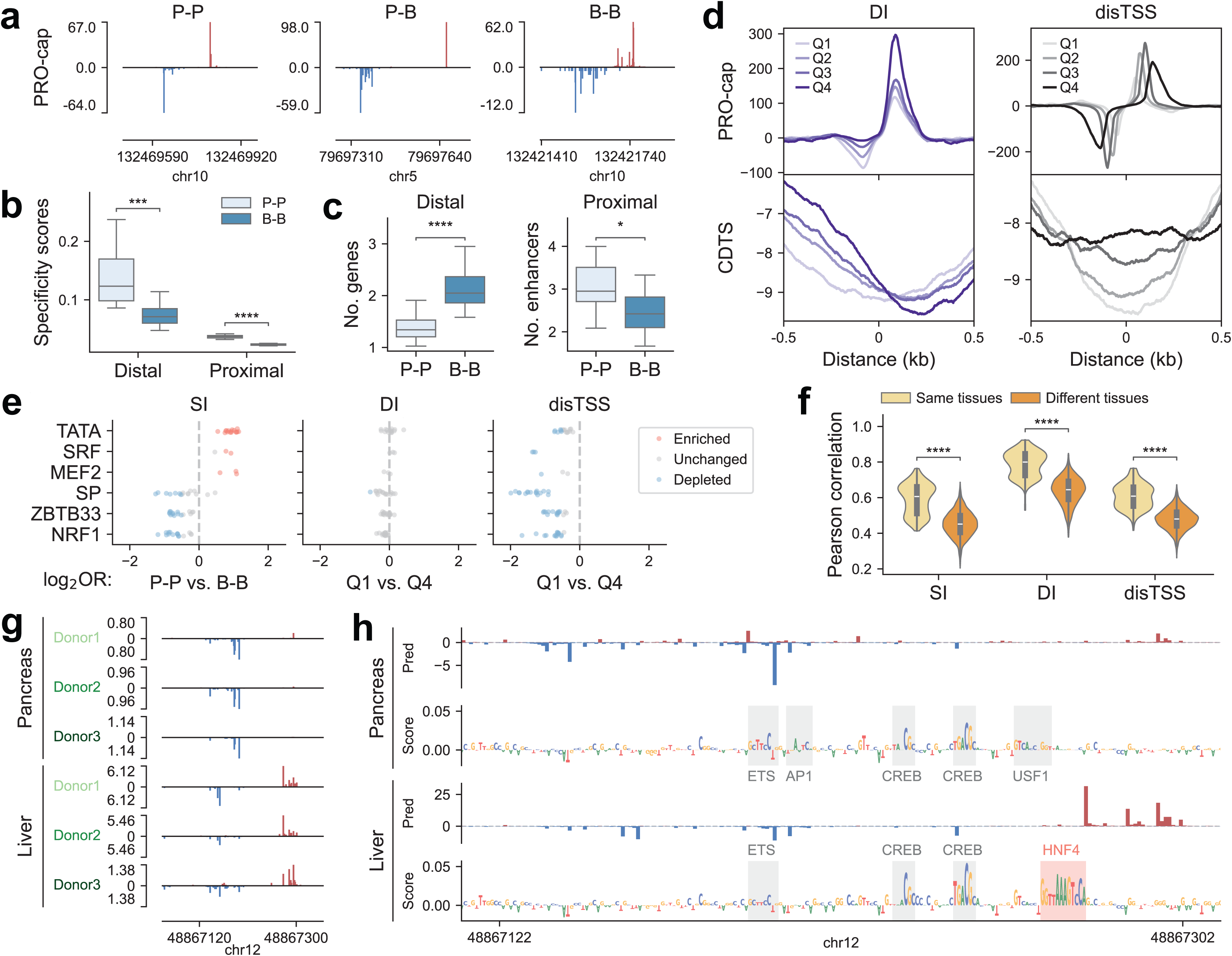
Architecture of transcriptional regulatory elements reflects evolutionary history and functional impact. **(a)** Representative 5’ PRO-cap signal tracks from one heart sample (EN23) for divergent distal TREs with P-P, P-B, and B-B peak shapes (P=peaked, B=broad). Forward and reverse strands are shown in red and blue. **(b)** Boxplot showing tissue specificity scores for divergent distal and proximal elements across peak shape categories. Each point represents the median score for a given tissue type (n=16). **(c)** *Left panel:* Violin plots showing the number of predicted target genes for distal elements with different peak shapes. *Right panel:* Violin plots showing the number of enhancers linked to genes with proximal elements of different peak shapes. Each datapoint reflects the mean value for a given tissue type (n=16). **(d)** Metaplots of PRO-cap signal and CDTS across divergent proximal TREs (n=22,035) in one heart sample (EN23). TREs are grouped by absolute DI values (Q1 = most balanced, Q4 = most skewed) or by disTSS (Q1 = nearest, Q4 = farthest). Distances are shown as ±0.5 kb from the element center. **(e)** Strip plot showing log2 odds ratio of representative TF motifs across different features (SI: P-P vs B-B; DI/disTSS: Q1 vs Q4). Each datapoint represents a tissue type (n=16). Red and blue denote motifs that are significantly enriched or depleted, respectively (padj < 0.05) in a given sample. **(f)** Violin plots showing Pearson correlation of SI, DI, and disTSS between shared TREs across pairs of samples from the same or different tissue types. Each point represents a pair of samples (236 same-tissue and 3,004 different-tissue comparisons). **(g)** Representative browser tracks of 5’ PRO-cap signals from two tissue types (pancreas and liver) obtained from three matched donors across one distal genomic locus. **(h)** Predicted transcription profiles and contribution scores (count task) for the locus indicated in (g), using ProCapNet models trained on pancreas (top) and liver (bottom) PRO-cap data. TF motifs contributing to transcription in each tissue type are labeled accordingly. *, 0.01 < padj ≤ 0.05; ***, 0.0001 < padj ≤ 0.001; ****, padj ≤ 0.0001.

This peak shape categorization revealed strong associations with expression breadth, evolutionary features, and gene regulatory function. Proximal B-B elements were broadly active across different tissues, whereas P-P elements exhibited high tissue specificity (Fig. 5b and Supplementary Fig. 11), consistent with prior work showing that broad promoters are enriched at housekeeping genes^65,66^. Strikingly, we found that this relationship also extends to distal elements, where B-B regions exhibit similarly ubiquitous activity across tissues. Consistent with their broader regulatory roles, both distal and proximal B-B elements showed stronger evolutionary constraints (Supplementary Figs. 12 and 13a), recapitulating the inverse relationship between tissue specificity and evolutionary conservation observed in Fig. 1d. To explore regulatory connectivity, we analyzed promoter-enhancer (P-E) associations using the ENCODE-rE2G^67^ predictions across various tissues (Supplementary Table 3). ENCODE-rE2G is a logistic regression model that predicts P-E links based on multiple molecular features, including chromatin state, 3D contact frequency, and promoter class. Distal B-B elements were linked to a greater number of putative target genes, whereas genes with proximal B-B elements tended to be regulated by fewer enhancers (Fig. 5c and Supplementary Fig. 14).

Peak shape captures the local distribution of TSSs within each strand. However, divergent TREs also exhibit variation in features that reflect the relationship between the two strands. We therefore examined two complementary properties: the balance of transcription between the two strands and the genomic distance between the dominant TSSs on forward and reverse strands. To quantify transcriptional directionality, we calculated a directionality index (DI) of each divergent element, defined as the difference in PRO-cap read counts between the plus and minus strands normalized by total signal. Elements were then stratified into quartiles based on absolute DI values, ranging from balanced (Q1) to strongly skewed initiation (Q4). Notably, the dominant transcriptional strand at highly skewed elements exhibited lower CDTS (Fig. 5d and Supplementary Fig. 13b), indicating stronger evolutionary constraint relative to more balanced elements. We then examined the distance between dominant TSSs on forward and reverse strands (disTSSs), categorizing elements into quartiles from closest pairs (Q1) to furthest apart (Q4). Widely spaced TSS pairs were associated with higher CDTS, reflecting reduced evolutionary constraint (Fig. 5d and Supplementary Fig. 13c). One possible explanation is that broader disTSS regions provide more sequence space for TF binding, allowing flexible regulatory configurations and thereby increasing tolerance of greater sequence variation. Interestingly, when assessing sequence age, closely spaced TSS pairs (Q1) are older at distal elements but show a weak and reversed relationship at proximal elements (Supplementary Fig. 12), arguing against a simple constraint-based model.

To explore the mechanistic basis of this variation in transcription initiation features, we examined differences in motif types utilized. Motifs with sharply localized contribution profiles (e.g. TATA, SRF, MEF2) were more likely to be enriched at P-P elements, whereas motifs with broader effects were preferentially associated with B-B elements (Fig. 5e and Supplementary Fig. 15), indicating that distinct TFs encode different initiation architectures. TREs with wider TSSs spacing were enriched for the majority of motif types, consistent with the idea that expanded initiation regions provide additional sequence space for TF binding.

Expanding on our analysis of intra-tissue variation in initiation profiles, we next quantified inter-tissue differences. All three initiation features showed higher correlations between samples from the same tissue type than between those from different tissue types, even within the same donor (Fig. 5f and Supplementary Fig. 16), supporting a role for tissue-specific regulatory programs in shaping initiation architecture. For example, at one representative distal locus, liver and pancreas samples from three matched donors showed consistent differences in directionality index, with strong forward-strand transcription uniquely observed in liver (Fig. 5g). We next asked whether TF motif usage preference underlies this inter-tissue difference. ProCapNet models of pancreas and liver accurately predicted the observed patterns, identifying both shared and distinct sets of motifs contributing to transcription (Fig. 5h). *In silico* mutagenesis further demonstrated that HNF4 motifs drive the liver-specific initiation pattern on the forward strand, consistent with higher HNF4A expression in the liver relative to pancreas based on GTEx RNA-seq (Supplementary Fig. 17). At another locus, heart and lung samples from three matched donors exhibited tissue-specific alternative TSS usage driven by distinct Inr motifs (Supplementary Fig. 18). These examples illustrate how motif preferences shape intra- and inter-tissue variation in initiation architecture, uncovering the breadth of regulatory patterns that remain to be explored across the dataset.

Having characterized initiation profiles, we asked whether pause position varies systematically across tissues. Leveraging the base-pair resolution of PRO-cap, we quantified pause distance as the distance between the TSS (5′ end) and pausing position (3′ end) at single-molecule resolution across tissues. Pause distances varied substantially, revealing three general classes: early (short), intermediate, and late (long) pausing (Supplementary Fig. 19), suggesting that pause distance may represent a stable, tissue-intrinsic regulatory feature.

To systematically assess regulators of pause site positioning, we conducted a targeted screen and integrated publicly available datasets using degron-tagged transcription-related proteins. While acute depletion of most factors did not substantially affect pause distances, depletion of NELF-C led to a global increase in pause distances (Supplementary Fig. 20), consistent with prior work^68^ showing coordinated downstream shifts in both TSS and pause positions that collectively increase transcript length. Given its ubiquitous expression, NELF-C is unlikely to account for inter-tissue variation in pause distances, which may instead reflect other factors, if any. Such variation may arise from changes in TSS usage, pause position, or both. Although not explored in depth here, our paired-end dataset, capturing both 5’ and 3’ ends of nascent RNA transcripts, enables systematic analysis of intra- and inter-tissue variation in 3’ pause position and its coordination with 5′ TSS architecture, providing a framework to disentangle contributions from initiation architecture and pausing and to interpret how tissue-specific regulatory programs shape transcriptional output.

### Tissue-specific TRE profiles inform primary site prediction in metastatic cancers

Beyond revealing principles of transcriptional regulation, a comprehensive atlas of tissue-specific TRE activity can also help address clinically relevant challenges. For instance, the clinical management of metastatic cancers can be complicated by the difficulty in identifying their tissue of origin, especially in cancers of unknown primary (CUP), which comprise 3-5% of diagnoses and frequently lack effective therapeutic strategies due to their elusive origins^69^. Despite advances in genomic profiling and imaging technologies, reliable molecular strategies for pinpointing the origin of CUP remain scarce. We therefore hypothesized that primary tumors retain regulatory features of their tissue of origin and that metastatic cancers preserve the regulatory signature of the primary tumor. Although primary tumors are not included in our dataset, the broad coverage of normal tissues may enable inference of metastatic origins.

To test this, we leveraged our nascent RNA-based regulatory dataset, which includes a diverse panel of metastatic tumors from known primary sites (n=23) and normal tissues from multiple organ systems (Fig. 6a and Supplementary Table1). We compared the PRO-cap profiles of divergent elements in metastatic cancer samples to those of normal tissues from the corresponding primary, metastatic, and other tissue types. Pairwise comparison revealed that most metastatic tumors more closely resembled their tissue of origin, supporting the hypothesis that metastatic cancers preserve the molecular signatures of their primary sites (Fig. 6a and Supplementary Fig. 21a). Additionally, correlations with metastatic sites were higher than with other tissues, suggesting an adaptation to the metastatic microenvironment.

**Figure 6.**
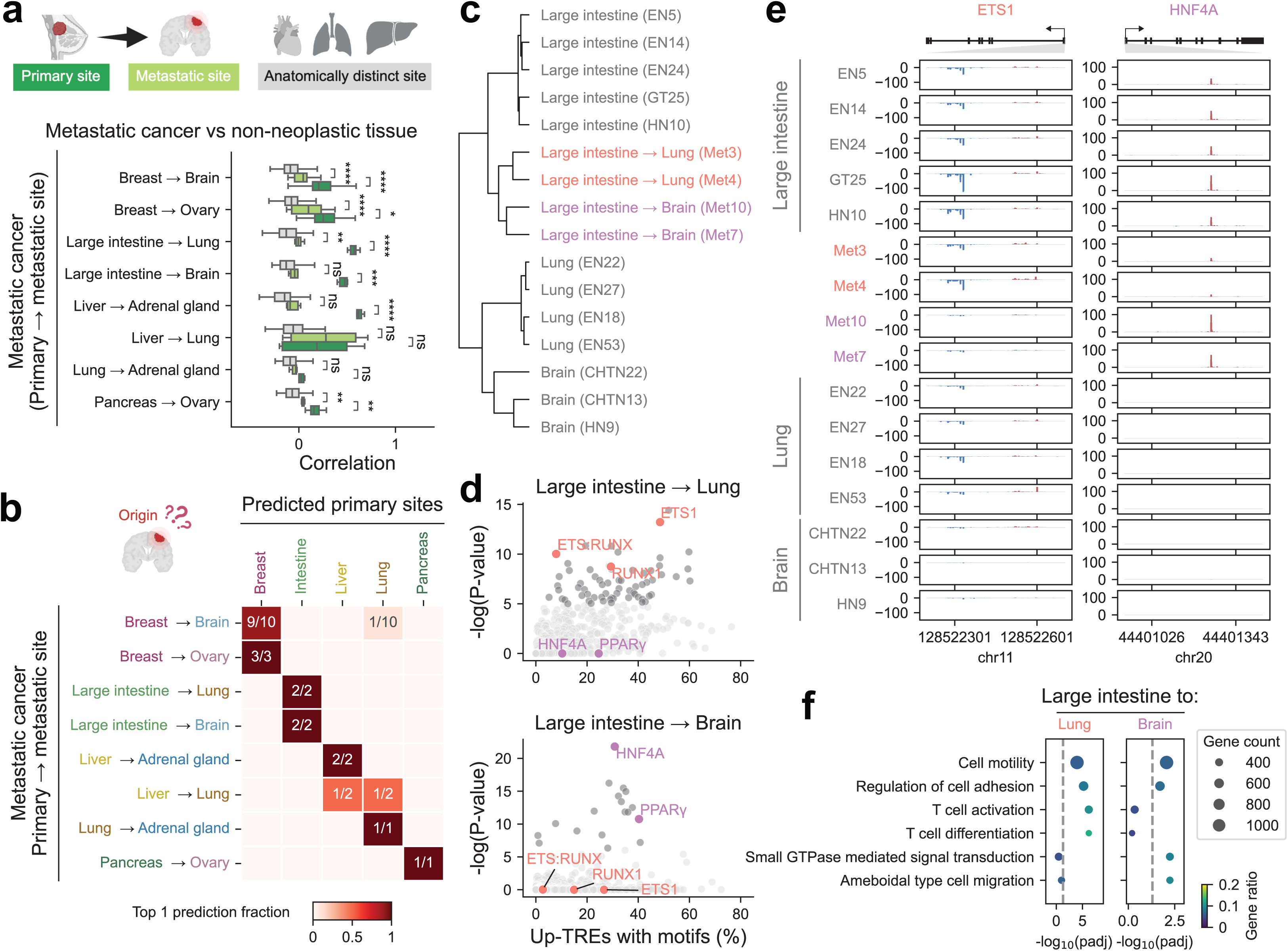
Transcriptional regulatory element signatures trace metastatic origins and facilitate organ-specific colonization. **(a)** *Top:* Schematic overview of anatomical sites profiled, including metastatic tumor biopsies (n=23) and non-neoplastic samples (n=93) from their corresponding primary and metastatic sites, as well as other unrelated tissues. *Bottom:* Pairwise comparison of divergent distal TRE expression profiles between metastatic tumors and non-neoplastic tissues from the corresponding primary (dark green), metastatic (light green), and unrelated sites (gray). ns, non-significant; *, 0.01 < padj ≤ 0.05; **, 0.001 < padj ≤ 0.01; ***, 0.0001 < padj ≤ 0.001; ****, padj ≤ 0.0001. **(b)** Top-1 prediction of the primary site origin for metastatic tumors using a linear support vector machine classifier. Prediction fraction is shown for each metastatic tumor type. **(c)** Hierarchical clustering of divergent distal TREs in metastatic biopsies from patients with colorectal cancer and related non-neoplastic tissues. **(d)** TF motif enrichment in differentially expressed TREs between lung and brain metastases from colorectal cancer. Top TF motifs associated with metastasis were highlighted in red (lung) and purple (brain). Remaining motifs were colored dark grey if the adjusted p-value < 0.05 and light grey otherwise. **(e)** Genome browser tracks of average PRO-cap signals at the promoter regions of *ETS1* and *HNF4A* in lung and brain metastases from colorectal cancer alongside related non-neoplastic tissues. The genes are depicted with arrows indicating transcription orientation and black rectangles marking exons. Grey triangles highlight regions shown in zoomed-in panels. **(f)** Pathway enrichment analysis of genes linked to differentially expressed TREs in colorectal cancer metastases to the lung and brain. Dot size reflects the number of genes in each pathway (Gene Ontology Biological Process terms). Gene ratio represents the fraction of pathway genes that overlap with the input gene set. Dashed lines indicate BH-adjusted p-value threshold of 0.05.

To evaluate the predictive utility of these patterns, we trained a linear support vector classifier using divergent distal elements from normal samples spanning 15 tissue types. Remarkably, although trained exclusively on non-neoplastic tissues, the model achieved an overall accuracy of 87.0% for the top-1 prediction and 95.7% for the top-3 predictions in identifying the primary origin of metastatic tumors (Fig. 6b and Supplementary Fig. 21b). Likewise, divergent proximal elements yielded comparable prediction accuracy. These findings demonstrate that nascent RNA profiling may offer a promising molecular strategy for resolving diagnostically ambiguous cases and guiding more precise therapeutic interventions.

### Site-specific metastatic adaptation via enhancer reprogramming

The preferential transcriptional similarity between metastatic tumors and their corresponding secondary sites, relative to unrelated tissues, suggests a high degree of plasticity (Fig. 6a and Supplementary Fig. 21a). This prompted us to examine whether specific enhancer programs underlie metastatic dissemination and organ-specific adaptation. We first performed hierarchical clustering of divergent distal TREs across metastatic biopsies from 4 patients with colorectal cancer (CRC) and related non-neoplastic tissues (i.e., large intestine, lung, and brain) (Fig. 6c). This analysis uncovered two major patterns: (i) both brain and lung metastatic lesions retained stronger regulatory similarity to their tissue of origin (i.e., large intestine) than to distant sites, which aligns with our prior observation; and (ii) within the metastatic cohort, tumors segregated by their destination site.

To interrogate the regulatory mechanisms underlying organ-specific dissemination and adaptation, we identified differentially expressed TREs (|log2FC| > 1 and adjusted p-value < 0.05) between lung and brain metastases (Supplementary Fig. 21c). Motif enrichment analysis of enhancers upregulated in lung metastases revealed a marked overrepresentation of ETS1, RUNX1, and their composite motifs ETS:RUNX (Fig. 6d top and Supplementary Table 2), consistent with their established roles in metastatic progression. Specifically, stromal ETS1 expression in primary CRC is associated with increased risk of dissemination to the lungs in patients^70^. Similarly, previous studies have demonstrated that RUNX1 promotes lung colonization of metastatic CRC in murine models, with elevated expression correlating with poor clinical outcomes^71^. Thus, our results extend these prior observations by suggesting that the coordinated activity of these TFs may shape the enhancer landscape in lung metastases of CRC. In contrast, enhancers upregulated in brain metastases showed significant enrichment for HNF4A and PPAR motifs (Fig. 6d bottom and Supplementary Table 2). Although PPARγ signaling and HNF4A expression have been linked to brain metastatic progression in melanoma^72,73^, their specific roles in CRC brain dissemination has yet to be elucidated. This selective enrichment of HNF4A and PPAR motifs in brain metastasis-specific enhancers thus uncovers a previously unrecognized regulatory axis and nominates these factors as compelling candidates for mediating colonization of the brain microenvironment.

To assess the potential driver roles of ETS1 and HNF4A, we first examined their expression patterns in metastatic cancers and non-neoplastic samples at their corresponding primary and metastatic sites. Both TFs exhibited elevated PRO-cap signals at their promoters in the corresponding metastatic contexts (i.e., ETS1 in Met 3/4 and HNF4A in Met 7/10), consistent with transcriptional activation (Fig. 6e). These factors are also expressed in non-neoplastic large intestine tissues, supporting the notion that metastases may co-opt pre-existing regulatory networks during dissemination and adaptation. To further contextualize regulatory programs associated with metastatic homing, we linked enhancers either upregulated in lung or brain metastases to their nearest expressed genes and performed pathway enrichment analysis of these putative targets. This analysis revealed shared pathways of cell adhesion and motility, highlighting core mechanisms of invasive spread in CRC metastasis (Fig. 6f). We also uncovered distinct, site-specific programs: lung metastases were enriched for pathways related to immune modulation, whereas brain metastases showed enrichment for small GTPase-mediated signaling and amoeboid cell migration, processes central to invasion across the blood-brain barrier and dissemination within the brain parenchyma^74^. Thus, these findings position ETS1 and HNF4A as key mediators of enhancer reprogramming, highlighting how metastatic tumors hijack lineage-primed regulatory circuits while reshaping their enhancer landscapes to colonize distinct organs. Together, these results underscore the importance of context-specific regulatory factors in driving the tissue tropism of metastatic cancer.

## Discussion

Transcription initiation precisely marks active gene promoters and enhancers^8,9,13,14^. Using PRO-cap across diverse developmental, physiological, and pathological contexts, we mapped transcription initiation at base-pair resolution and identified hundreds of thousands of active TREs. This comprehensive resource provides a unified view of regulatory activity across biological contexts, revealing the breadth of regulatory elements that shape cellular and tissue identity in health and disease.

Beyond its scale, this atlas provides key conceptual insights. First, our analyses demonstrate that transcriptional initiation identifies a functionally enriched set of regulatory elements within the broader landscape defined by epigenomic annotations. DNase I and ATAC hypersensitive sites can mark a wider range of open chromatin regions than those associated with active TREs, while histone modification patterns are most informative for predicting enhancers or promoters when multiple ChIP-seq datasets are integrated. By directly capturing transcriptional activity at regulatory elements, PRO-cap provides an effective, unified transcription-based framework (one-stop-shop) for prioritizing active enhancers and promoters. As shown here, transcription-defined TREs exhibit stronger enhancer activity, greater evolutionary constraint, and increased enrichment for trait-associated variants, establishing transcription initiation as a more precise and functionally informative indicator of regulatory activity with clear biological and clinical relevance. Notably, while CAGE-seq–based FANTOM annotations have provided foundational maps of transcription initiation, our PRO-cap-based profiling extends these resources by identifying a substantially larger and more sensitive repertoire of active regulatory elements, reflecting improved detection of transcription initiation at TREs.

Second, we uncover a tight coupling between tissue specificity of regulatory activity and evolutionary constraint. Broadly active TREs are more conserved and exhibit signatures of stronger purifying selection, whereas tissue-restricted elements display increased evolutionary turnover. This relationship extends prior observations from epigenomic studies by directly linking nascent RNA output at TREs to selective pressures, suggesting that patterns of transcription initiation reflect both regulatory function and evolutionary history.

Third, the base-pair resolution of PRO-cap enables systematic dissection of transcription initiation architecture across tissues. By quantifying features such as peak shape, transcriptional directionality, and spacing between divergent TSSs, we show that initiation profiles encode distinct regulatory behaviors. These architectural features are associated with tissue specificity, TF usage, and regulatory connectivity, indicating that initiation is not merely a marker of activity but also a structured and information-rich layer of gene regulation. In parallel, we find that transcriptional pausing also varies systematically across tissues and may be modulated by specific regulatory factors. These observations link transcription initiation and elongation dynamics, suggesting that variation in pause positioning represents an additional layer of regulatory control contributing to tissue-specific transcriptional programs. Thus, our PRO-cap-based approach reveals regulatory features that are not accessible through conventional profiling methods and enables mechanistic dissection of transcriptional features at TREs underlying intra- and inter-tissue variation.

Fourth, integrating TRE maps with deep learning models and statistical fine-mapping provides a framework for prioritizing noncoding variants from GWAS, eQTL, and other studies in a tissue-specific context. By linking sequence features to transcription initiation and quantifying predicted regulatory effects, we prioritize candidate causal variants with increased confidence and enable mechanistic interpretation of their functional impact across cellular contexts.

Finally, the tissue-resolved nature of this atlas enables the identification of lineage-specific regulatory programs and their alterations in disease. In cancer, TRE landscapes capture regulatory signatures that reflect both the tissue of origin and adaptations to metastatic environments. These patterns can be leveraged to trace the origins of metastatic tumors and to identify regulatory circuits associated with tissue-specific dissemination, illustrating the translational potential of transcription-based regulatory profiling.

Overall, these findings position transcription initiation as a defining and mechanistically informative layer of gene regulation that complements and refines existing epigenomic annotations. By combining scale with base-pair resolution, this atlas enables systematic interrogation of regulatory element activity, architecture, and variation across biological contexts and provides a framework for integrating transcriptional initiation with complementary genomic and epigenomic measurements.

More broadly, placing these results in the context of other large-scale efforts highlights convergent insights into regulatory genome organization and supports a more unified model of transcriptional regulation across biological systems. Integration of TRE maps with gene expression and chromatin state data, including RNA-seq, PRO-seq, and epigenomic profiling, will further refine models of gene regulatory output by linking initiation dynamics to downstream transcriptional consequences.

Collectively, this detailed atlas of active TREs, representing all major human organs, pluripotent and their differentiated lineages, and several disease states, including T1D, leukemia, lymphoma, and a variety of metastatic cancers, provides an essential resource for dissecting gene regulation and advancing applications from basic molecular biology to clinical genomics. Beyond the analyses presented here, the scale and base-pair resolution of this dataset enable systematic interrogation of regulatory features not exhaustively examined in this study, including the landscape of unidirectional TREs, shifts in transcriptional directionality between divergent and unidirectional initiation across biological contexts, and alternative promoter usage at genic loci. The base-pair resolution of PRO-cap enables precise delineation of both transcription start sites and pause positions, enabling high-resolution analyses of the architecture of active TREs, pause positioning, and regulatory logic across diverse biological contexts.

## Methods

### Sample collection and PRO-cap library preparation

Snap frozen human tissue samples from cadaveric or surgical biopsies were obtained from a range of sources, including ENCODE, and GTEx, as part of the ENCODE consortium’s coordination Phase 4 efforts, and the National Cancer Institute’s Cooperative Human Tissue Network (CHTN). Solid tissues were pulverized on dry ice using a mortar and pestle. Peripheral blood mononuclear cells (PBMCs) were isolated using Lymphoprep, and specific immune cell populations were purified from LeukoPaks via fluorescence-activated cell sorting (FACS) using positive or negative selection based on surface marker expression. Cells were either cultured in-house or obtained as snap-frozen dry pellets. PRO-cap libraries were prepared as previously described, with modifications to streamline the protocol, reducing the experimental time from ∼4 days to ∼10 hours, while enabling the use of limited tissue input and low cell numbers from diverse sample types, without compromising the assay’s sensitivity or specificity. Briefly, permeabilized cells or pulverized tissues underwent nuclear run-on reactions to capture nascent RNA. Total RNA was then isolated and subjected to two rounds of custom adaptor ligation and reverse transcription. Between adaptor ligations, 5′ cap selection was performed through a series of enzymatic reactions to enrich for capped nascent transcripts. RNA was washed, followed by phenol:chloroform extraction and ethanol precipitation at each step, all under RNase-free conditions. Following PCR amplification and library clean-up, sequencing was performed on an Illumina NovaSeq platform. Likewise, we generated PRO-cap libraries from two replicates, each of 10 million HCT116 cells genetically modified using CRISPR targeting H. sapiens BRD4, CDK7, CTCF, MED14, POLR2A, RAD21, SMARCA5, SUPT16H and O. sativa LOC4335696 (OsTIR1 auxin receptor for the auxin-inducible degron system), before and after 6 hours of treatment with 1 μM 5-Phenyl-1H-indole-3-acetic acid. Details of all biosample and library information can be found in Supplementary Table 1.

### Data preprocessing

This dataset was managed and analyzed using the Resource Management System (https://github.com/aldenleung/rmsp/). Raw reads were preprocessed with fastp75 (v0.23.4) for adapter trimming and unique molecular index (UMI) processing, retaining only reads ≥18 bp for downstream analyses. Processed reads were aligned to the human reference genome hg38 (GCA_000001405.15) and ribosomal DNA (U13369.1) using STAR76 (v2.7.11a). Uniquely mapped reads were filtered with samtools77 (v1.18) and deduplicated using umi_tools (v1.1.5). FASTQ files of PRO-cap data from the NELF-C degron lines (NELFC_U, NELFC_T) were obtained from GSE144786 and reprocessed with the same pipeline, except that PCR deduplication was omitted due to the lack of UMI information.

The resulting PCR-free, uniquely mapped reads were converted to BigWig format using biodatatools (v0.0.7). For replicates with good inter-replicate correlation, BigWig files were merged prior to peak calling. Peak calling and classification were performed with our in-house tool PINTS^17^ (v1.1.10). Briefly, PINTS applies zero-inflated Poisson models to evaluate local read densities and employs an interquartile range-based refinement step to reduce false negatives via conditional masking of candidate peaks in the local background. Statistically significant peaks are then classified as divergent elements if they form peak pairs on opposite strands within 300 bp of each other, and as unidirectional elements otherwise.

After peak calling in individual samples, we observe that some elements were classified as unidirectional in certain samples but divergent in others, which may reflect biological variation across cell and tissue types and/or technical factors such as sequencing depth. To obtain consistent labels across the dataset, we merged elements from all samples to define a union set, which was used for the majority of analyses in the paper. Within this union set, an element was classified as unidirectional if it was called unidirectional in all samples in which it was detected, and as divergent if it was called divergent in at least one sample. Elements in the union set were further categorized as proximal (within ±500 bp) or distal (outside ±500 bp) relative to the TSSs of all transcripts annotated in GENCODE^78^ v37. Specifically, we extracted TSSs for all transcripts, defined ±500 bp windows around each TSS, and merged overlapping windows to generate a unified set of proximal regions. Elements overlapping any merged windows were classified as proximal, whereas those that did not overlap were classified as distal.

Median RNA length was reported for each sample, reflecting the distribution of transcript lengths. RNA length was defined as the distance from the 5′ end (TSS) to the 3′ end (pausing site) and is also referred to as pause distance in downstream analyses. The gene body ratio was calculated as a quality control metric to assess 5′ enrichment in capped RNA sequencing experiments. Analyses were restricted to highly expressed genes, defined as the top 10% ranked by PRO-cap signal in a promoter-proximal window. For each gene, PRO-cap counts were summed from −100 to +1000 bp relative to the TSS of each transcript, and the maximum value across transcripts was used for ranking. For the selected genes, all annotated transcripts were then used to define genomic regions. Specifically, TSS regions were defined as 500 bp windows downstream of each transcript TSS and merged across transcripts to generate a union set for each sample. Similarly, transcription termination site (TTS) regions were defined as windows extending 500 bp upstream of each transcript TTS and merged across transcripts to generate a union set. Gene body regions were defined as all remaining regions not overlapping TSS or TTS regions. The gene body ratio was computed as the length-normalized PRO-cap signal in gene body regions divided by the sum of length-normalized PRO-cap signals in gene body and TSS regions, with each region normalized by its own length. This measure reflects the extent to which capped reads are concentrated at TSS regions in high-quality data. One sample with a gene body ratio ≥ 0.025 was excluded from downstream analyses.

Details of sequencing and quality control metrics are provided in Supplementary Table 1.

### Quantification of transcription initiation

For each sample, we counted reads whose 5’ ends aligned within the boundaries of individual elements in the union set. The resulting count matrix was normalized using the median of ratios method in DESeq2^53^ to account for differences in library size and RNA compositional bias. We then applied variance stabilizing transformation to mitigate the mean-variance dependency, as recommended by DESeq2, for downstream analyses including PCA and clustering.

### Calculate specificity scores

The specificity scores were calculated following an approach similar to that previously described^9,79^. All samples from Fig. 1b were included, and *q_e,t_* was defined as the average normalized expression of an element *e* across samples of the same tissue type *t*. We first converted *q_e,t_* to probabilities 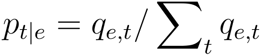. The entropy of a given element was then computed as 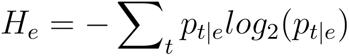. To constrain the specificity score within the range of 0 to 1 (where 0 indicates ubiquitous expression and 1 represents exclusively specific expression), we defined 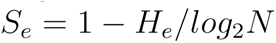, with *N* denoting the number of tissue types.

### Get tissue-specific and non-tissue-specific elements

We computed a *t*-statistic for the specific expression of each divergent distal or proximal element in tissue types with at least three samples, as highlighted in Fig. 1b, following the procedure described in Finucane *et al*^80^. Briefly, a design matrix *X* was constructed with an indicator column denoting sample membership in the given tissue type (1 for yes, -1 for no) and an intercept term. The outcome *Y* was the normalized expression level of a given element across samples. We fit the model via ordinary least squares and calculated the *t*-statistic for the membership variable. The top 5% of elements from the union set, ranked by their *t*-statistic, were defined as the specifically expressed set for each tissue type. Non-tissue-specific elements were defined by the following criteria: (1) distal elements with a specificity score < 0.1 and proximal elements with a specificity score < 0.02 were included, resulting in a number of elements similar to that of the tissue-specific set for each tissue type; (2) elements belonging to any tissue-specific set were excluded.

### Sequence age and evolutionary metrics

To estimate the sequence age of TREs, we employed a syntenic block-based approach following Fong *et al*^22^. Specifically, we obtained the hg38 100-way vertebrate Multiz alignment data from the UCSC database^81^ and assigned an age to each syntenic block based on the most recent common ancestor (MRCA) of the species in the alignment block. We grouped MRCA nodes into 10 age categories and reported the age as the oldest ancestral branch, as implemented in our previous work^21^. Evolutionary constraint across species was quantified using 447-way phyloP scores retrieved from the UCSC database. To capture more recent selective pressures within the human lineage, we additionally examined CDTS derived from 7,794 unrelated individuals, as reported previously^24^. In Fig. 2d, we compared these metrics between cCREs with and without overlapping PRO-cap elements using two-sided Mann-Whitney U tests. For each element, we summarized a 1-kb window centered on the peak by taking the maximum sequence age (higher values indicate older origin) and phyloP score (higher values indicate greater conservation), and the minimum CDTS value (lower values indicate reduced tolerance to variation).

### Comparing PRO-cap TREs with VISTA enhancer annotations

We obtained experimental data (as of 2024/12/02) from the VISTA Enhancer Browser (https://enhancer.lbl.gov/vista/), restricting the analyses to human elements with the reference allele, regardless of plasmid backbone, transgenic method, or embryonic stage. We then focused on elements validated as enhancers in at least one of the following mouse embryonic tissues: pancreas, blood vessels, neural crest-derived tissues (dorsal root ganglion, trigeminal ganglion, and cranial nerve), and neural tube-related regions (neural tube, forebrain, midbrain, and hindbrain), as these are relevant to cell lines assayed in our ESC/iPSC differentiation panel (Fig. 1e). Elements not validated as enhancers in any VISTA tissue were also included as negative controls.

To assess whether VISTA-validated enhancers exhibit elevated PRO-cap signal in relevant cellular contexts, we first calculated Z-scores of PRO-cap signal for each divergent element across samples in the ESC/iPSC panel. For Fig. 1e, we grouped VISTA elements into categories defined by enhancer activity in a given tissue (e.g., pancreas) or by lack of activity in all tested tissues (negative controls). Each tissue category includes all elements validated in that tissue, regardless of whether they are also active in other tissues. For each VISTA category (rows), we intersected the union set of PRO-cap–detected divergent elements from our ESC/iPSC panel with the corresponding VISTA elements, requiring the overlap to span at least 50% of the PRO-cap element length. This yields a fixed set of overlapping PRO-cap elements per category that is identical across all PRO-cap samples (columns). We then visualized the average Z-scores of these elements for each sample in the heatmap.

### Time series clustering of dynamically transcribed TREs during pancreatic lineage differentiation

We took the union set of PRO-cap elements across four timepoints (ESC, ESC-derived endodermal cells, pancreatic progenitor cells, and insulin-producing beta-like cells) to generate a count matrix. DESeq2 was applied to identify elements with differential PRO-cap signals between any pair of timepoints, using a Benjamini-Hochberg (BH)-adjusted p-values, with a significance threshold of 0.01/6 to account for six pairwise comparisons. Dirichlet process-Gaussian process (DP-GP) time-series clustering^82^ was applied to categorize the dynamically transcribed elements into distinct trajectories over time. We followed a similar two-stage strategy described in Kim *et al*^83^, except that replicate reproducibility was not required due to the absence of biological replicates in our dataset. First, we subsampled the expression data (n=5,000 for computational efficiency, as the algorithm was originally designed for thousands of genes) and applied the DP-GP algorithm with default parameters to generate the initial set of time series clusters. The cluster set was further filtered by excluding (1) clusters with fewer than 2% of dynamically transcribed elements and (2) non-dynamic trajectories whose multivariate Gaussian process did not reject the null hypothesis of no change over time. In the second stage, each dynamically transcribed element was assigned to a cluster if it fell within the 95% multivariate confidence interval of the trajectory. If multiple clusters matched, the element was assigned to the one with the smallest Euclidean distance from the mean trajectory. Elements without a matching cluster were discarded.

### Pathway enrichment analysis

To validate the biological relevance of TRE categories, we assessed whether their putative target genes were enriched for expected pathways (Extended Data Fig. 4b: TRE clusters from pancreatic lineage differentiation; Extended Data Fig. 7b: TREs up- or downregulated in T cells from T1D patients compared to controls; Fig. 6f: TREs up- or downregulated in colorectal cancer metastases to lung versus brain). Elements were linked to the nearest gene expressed in at least one sample within the panel of interest. For the pancreatic lineage differentiation panel, a gene was considered expressed in a sample if its TPM > 1 in both ENCODE RNA-seq replicates (ESC: ENCFF274WSK, ENCFF216CFE; endodermal cells: ENCFF599HQU, ENCFF643LBM; pancreatic progenitor cells: ENCFF111ORO, ENCFF411UJX; insulin-producing beta-like cells: ENCFF299BMI, ENCFF206GDB). The expressed gene set was then defined as the union of genes expressed in at least one sample across the differentiation time course. For the T1D and metastatic panels, in the absence of RNA-seq data, expressed genes were defined as those whose gene TSS overlapped a proximal element detected by PRO-cap in at least one sample within a given panel. Gene TSS coordinates were obtained from a curated RefSeq-based gene set selecting the TSS with the highest number of coding isoforms per gene (https://github.com/broadinstitute/ABC-Enhancer-Gene-Prediction/blob/main/reference/hg38/CollapsedGeneBounds.hg38.TSS500bp.bed). Pathway enrichment analysis was performed using GSEApy^84^, with all expressed genes within each panel used as the background. Gene sets were obtained from the Molecular Signatures Database (MSigDB)^85^, including Reactome Canonical Pathways (c2.cp.reactome.v2024.1) for the pancreatic differentiation and T1D panels, and Gene Ontology Biological Process terms (c5.go.bp.v2024.1) for the metastatic samples.

### Motif enrichment analysis

For the leukemia and lymphoma panel (Extended Data Fig. 5a), we performed *de novo* motif discovery with HOMER^86^ for each TRE module, using the union set of divergent distal elements from this panel (excluding TREs belonging to the module being tested) as the background. The top and biologically relevant motifs identified in each module were compiled into a custom motif set, which was then used to assess motif enrichment across all modules. For the T1D (Extended Data Fig. 7a) and metastatic (Fig. 6d) panels, we performed known motif enrichment using HOMER with the default motif database. TREs from the unchanged group were used as the background for enrichment analysis separately for up- and down-regulated elements. The full list of motif enrichment results is provided in Supplementary Table 3.

### CRISPRi screening at the *MYC* locus

B-ALL cell lines REH, SEM, and NALM-6 were co-transduced with virus generated from lentivectors TRE-KRAB(ZNF10)-dCas9-IRESGFP (RRID: Addgene_85556) and EF1a_TetOn3G (Clontech), and were serially flow sorted to derive populations that were GFP-negative in the absence of doxycycline and GFP-positive after induction with 500 ng/ml doxycycline. For SEM, a polyclonal population of inducible KRAB-dCas9+ cells was used, while for REH and NALM-6, single cell clones were derived by limiting dilution and clones with uniform transgene inducibility were validated and used for screening. Gene knockdown efficiency was validated by qRT-PCR after transduction with sgRNAs targeting the ENO1 promoter or non-genome-targeting control in the presence and absence of doxycycline. Inducible KRAB-dCas9+ cell populations were validated for cell line identity by STR profiling prior to the CRISPRi screen. CRISPRi screens to determine fitness effects of *MYC* locus TRE modules were performed in B-ALL cell lines via the same sgRNA library and protocol previously described for K562^87^ and KARPAS-422^35^. CRISPRi screening results from SU-DHL-5 was previously published^35^, and used a different sgRNA library targeting nucleosome-free regions of *MYC* locus elements with significant acetylation in mature B-cell lymphoma cell lines.

### Percent overlap of PRO-cap elements with other annotations

We focused on non-diseased tissue samples and listed the mapping of matched tissue types across datasets in Supplementary Table 3. We obtained peak files of ATAC-seq, DNase-seq, H3K27ac ChIP-seq, and cCRE annotations (dELS, pELS, and PLS only) for each biosample from the ENCODE portal. For FANTOM CAGE-seq data (https://fantom.gsc.riken.jp/5/datafiles/reprocessed/hg38_latest/extra/), peaks are defined as a single consensus set across the dataset (CAGE_peaks/hg38_fair+new_CAGE_peaks_phase1and2.bed.gz), rather than as sample-specific peak calls. To obtain peaks active in each sample, we retained those with TPM > 1 based on the corresponding quantified file (CAGE_peaks_expression/hg38_fair+new_CAGE_peaks_phase1and2_tp m.osc.txt.gz).

For pairwise comparisons between datasets, elements were first merged across samples of the same tissue type within each dataset. For example, PRO-cap peaks (CHTN11, GT1) and ATAC-seq peaks (ENCSR914DTI, ENCSR201FIW, ENCSR474XFV) from the thyroid gland were merged separately prior to comparison. We then anchored on PRO-cap elements (divergent only) and quantified their percent overlap with other annotations (Fig. 2a, top), followed by a reciprocal analysis in which we anchored on other annotations and assessed their percent overlap with PRO-cap elements (either divergent or unidirectional) (Fig. 2a, bottom). A minimal 1-bp overlap criterion was used to maximize sensitivity for detecting shared regions. Even under this permissive definition, substantial discrepancies remain for certain annotation pairs, suggesting that these differences are unlikely to be driven solely by stringent overlap thresholds.

### ENCODE STARR-seq activity of cCREs with and without transcription marks

To evaluate enhancer activity of elements with and without transcription marks, we first adopted a cell- and tissue-agnostic approach. We leveraged the latest cCRE set (dELS, pELS, and PLS only; n = 1,766,201) from Moore *et al*.^2^ as a reference (https://users.moore-lab.org/ENCODE-cCREs/Supplementary-Data/Supplementary-Data-1.GRCh38-cCREs-V4.bed.gz) and categorized elements into four groups based on their 1-bp overlap with all FANTOM CAGE-seq peaks (hg38_fair+new_CAGE_peaks_phase1and2.bed.gz) and all PRO-cap elements from our dataset. Specifically, cCREs were classified as: (1) “CAGE-seq(-), PRO-cap(-)”: overlapping neither CAGE-seq nor PRO-cap, (2) “CAGE-seq(+), PRO-cap(-)”: overlapping CAGE-seq only, (3) “CAGE-seq(-), PRO-cap(+)”: overlapping PRO-cap only, or (4) “CAGE-seq(+), PRO-cap(+)”: overlapping both. In Moore *et al*., cCRE activity was evaluated using seven ENCODE STARR-seq datasets (ENCSR895FDL, ENCSR064KUD, ENCSR858MPS, ENCSR547SBZ, ENCSR135NXN, ENCSR983SZZ, and ENCSR661FOW) from six cell lines (A549, HCT116, K562, MCF7, HepG2, and SH-SY5Y). cCRE were considered active if they met either of the following criteria: (1) at least 50% of the cCRE overlapped a STARR-seq peak; (2) the cCRE had a significant STARR score (score > 0; p < 0.05) based on CRE-centric Analysis and Prediction of Reporter Assays (CAPRA), using either solo or double quantification schemes. CAPRA is a method that uses cCREs as predefined anchors to directly quantify STARR-seq activity without requiring peak calling. We calculated the active rate of cCREs across the categories described above to assess how cCREs with and without transcriptional marks, and with different types of transcriptional evidence (PRO-cap or CAGE-seq), differ in enhancer activity. Next, we evaluated cell-type specific enhancer activity by focusing on cCREs derived from K562 (ENCFF414OGC_ENCFF806YEZ_ENCFF849TDM_ENCFF736UDR.bed). These were categorized based on their 1-bp overlap with PRO-cap elements as: (1) detected in K562, (2) not detected in K562 but present in other biosamples in our dataset, or (3) not found in any biosamples in our dataset.

We then calculated the active rate of each category using K562 STARR-seq datasets (ENCSR858MPS and ENCSR661FOW). To test differences in active rate between a given pair of categories, we used two-sided Fisher’s exact test to calculate p-values and displayed BH-adjusted p-values in Fig. 2b, c.

### Heritability partition using S-LDSC

A total of 176 GWAS summary statistics were obtained from the curated collection of the Alkes Price lab (https://console.cloud.google.com/storage/browser/broad-alkesgroup-public-requester-pays/LDSCORE/all_sumstats/). S-LDSC analyses were performed across multiple element sets and GWAS phenotypes to address distinct biological questions, using two complementary metrics: (1) heritability enrichment, defined as the proportion of SNP heritability explained by an annotation divided by the proportion of SNPs it contains, and (2) the regression coefficient *τc*, which estimates the contribution of an annotation C to per-SNP heritability conditional on the other annotations in the joint model. When comparing tissue and cell types, it is important to account for overlap between tissue- and cell-type specific annotations and other functional categories^42^.

In Fig. 2e, we analyzed cCREs (dELS, pELS, and PLS only; n = 141,970) derived from adult heart samples, stratified by 1-bp overlap with PRO-cap elements in matched heart tissues, and reported heritability enrichment for a representative cardiac trait (systolic blood pressure). To estimate enrichment for these cCRE annotations with and without PRO-cap, we included them alongside the baselineLD v2.2 model (LDSCORE/GRCh38/baselineLD_v2.2.tgz), where “Coding” in Fig. 2e corresponds to the “Coding_UCSC” annotation in the baseline model. BH-adjusted p-values of heritability enrichment are shown in the figure.

In Fig. 3a and Supplementary Fig. 4, we used PRO-cap elements defined in the “Get tissue-specific and non-tissue-specific elements” section to capture both tissue-specific and non-tissue-specific effects across all available GWAS summary statistics. In Fig. 3d and Extended Data Fig. 6, we included all divergent distal PRO-cap elements identified in each blood sample to assess heritability enrichment of immune-related phenotypes (e.g., blood cell counts, IBD, and diabetes). For these tissue- and cell type-specific analyses, we followed the recommended practice of using the baseline v1.2 model (LDSCORE/GRCh38/baseline_v1.2.tgz) and ranked tissues and cell types by the BH-adjusted p-values of the regression coefficient *τc*.

### Luciferase assay for evaluating GWAS variant effects on enhancer activity

In the section “Tissue-specific effects of disease- and trait-associated variants”, we assessed the effect of the UC-associated variant rs6426833 on enhancer activity using a dual-luciferase reporter assay. Elements (chr1:19844867-19845867) containing either the reference or alternative alleles were cloned following previously described protocols^13^. EGFP and CMV584/RSV sequences were used as negative and positive controls, respectively. Primers were designed using our in-house web tool^88^, incorporating attB1′ (forward) and attB2′ (reverse) 5′ overhangs (Supplementary Table 4). K562 genomic DNA (E493; Promega Corp.) served as the template for PCR amplification using Phusion High-Fidelity (M0530; New England Biolabs) and PrimeSTAR GXL (R050A; Takara Bio Inc.) DNA polymerases. Amplicons were inserted into pDONR223 via Gateway BP cloning, sequence verified, propagated in spectinomycin-supplemented lysogeny broth (LB), and purified with E.Z.N.A. Plasmid DNA Mini Kit II (D6904; Omega Bio-tek, Inc.). Verified elements were transferred to pDEST-hSTARR-luc-pMYC via Gateway LR cloning, propagated in ampicillin-supplemented LB, and extracted using E.Z.N.A. Endo-Free Plasmid DNA Midi Kit (D6915; Omega Bio-tek, Inc.).

HCT116 (CCL-247; ATCC) cells were cultured in McCoy’s 5A Medium (30-2007; ATCC) supplemented with 10% FBS (30-2020; ATCC) at 37°C with 5% CO2. Caco-2 (HTB-37; ATCC) cells were cultured in EMEM (30-2003; ATCC) supplemented with 20% FBS (30-2020; ATCC) under the same conditions. Vectors were transfected into HCT116 and Caco-2 cells using Lipofectamine 3000 (L3000001; Invitrogen), with 0.5 x 10^6^ cells receiving 1 μg of pDEST-hSTARR-luc-pMYC and 10 ng of pGL4.75 (E6931; Promega Corp.). After 24h of incubation, cells were dissociated with 0.25% Trypsin-EDTA (25200056; Gibco) for the dual-luciferase reporter assay. Luminescence was measured using the Dual-Glo Luciferase Assay System (E2920; Promega Corp.) on an Infinite M1000 microplate reader (30034301; Tecan Group Ltd.) following the manufacturer’s instructions. Cells transfected with only pDEST-hSTARR-luc-pMYC or only pGL4.75 were used as background controls for firefly and Renilla luciferase activities, respectively.

In Fig. 3c, we used a two-sided Welch’s t-test to compare the relative luciferase activity between two alleles of rs6426833 (G vs. A) in Caco-2 and HCT116 cells (three replicates per condition), and reported BH-adjusted p-values in the figure.

### eQTL enrichment using TORUS

eQTL summary statistics were obtained from 28 GTEx v8 tissue types matching those in our PRO-cap dataset (https://console.cloud.google.com/storage/browser/gtex-resources/GTEx_Analysis_v8_QTLs/GTEx_Analysis_v8_EUR_eQTL_all_associations). We applied TORUS to estimate functional enrichment for each annotation described in the “Get tissue-specific and non-tissue-specific elements” section. The tool outputs 95% confidence intervals for the log enrichment parameters, from which p-values are derived under the assumption of asymptotic normality^89^.

### Fine-mapped GWAS and eQTL variants

Both fine-mapped eQTL variants for GTEx tissues and GWAS variants for UK Biobank traits were obtained from https://www.finucanelab.org/data/. In this dataset, fine-mapping was performed using FINEMAP^90^ and SUSIE^91^. Variants included in the primary release were selected as follows: for GWAS, those within the 95% CS or with PIP > 0.001; for eQTL, those within the 95% CS or with PIP > 0.0001. We analyzed data from both fine-mapping approaches and observed consistent trends. For clarity and conciseness, only the FINEMAP results are presented in the manuscript.

### ProCapNet models and related analyses

We trained the ProCapNet models (https://github.com/kundajelab/ProCapNet) for 16 samples using default parameters, except that *in_window* and *out_window* were set to 1000 and 500, respectively. The PRO-cap sample IDs and matched chromatin data (DNase-seq or ATAC-seq) are listed in Supplementary Table 3.

We computed base-level contribution scores using DeepSHAP^58^ for both count and profile tasks. TF-MoDISco^59^ was then applied to cluster subsequences with high contribution scores across all PRO-cap peaks into a lexicon of motif patterns. In Fig. 4a, we highlighted motifs selected using the following criteria: (1) robust correspondence to known motifs (Tomtom^92^ q-value < 0.1 followed by manual inspection; (2) exclusion of simple GC-rich repeats; and (3) support from at least 200 seqlets identified by TF-MoDISco. Motif instances of these learned patterns were then identified using Fi-NeMo (https://github.com/kundajelab/Fi-NeMo), a GPU-accelerated hit-calling method for locating occurrences of TF-MoDISco motifs in contribution score tracks. To assess whether motifs identified in specific tissues and cell types are associated with expression of their cognate transcription factors, we examined median gene-level log2TPM values per tissue from GTEx RNA-seq data (https://storage.googleapis.com/adult-gtex/bulk-gex/v11/rna-seq/GTEx_Analysis_2025-08-22_v11_RNASeQCv2.4.3_gene_median_tpm.gct.gz).

To assess variant effects across diverse contexts (Fig. 4b,c), we applied these ProCapNet models to 1-kb input sequences centered on PRO-cap elements containing either the reference or alternate allele. Variant effects were quantified as the log_2_ fold change in total counts between the alternate and reference alleles. To evaluate the effect of motif ablation on transcription initiation (Fig. 5f and Supplementary Fig. 17a), we replaced the one-hot encoding of each base in a given motif instance with [0,0,0,0] to simulate *in silico* deletion.

### Profile features

To obtain a more accurate characterization of profile features, we used the original boundaries of divergent elements identified by PINTS in individual samples, rather than the union set. We examined three profile features defined as below:

i. To calculate shape index (SI) for a given strand, we applied the formula described in Hoskins *et al*.^66^, 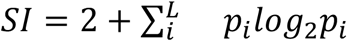, where *p_i_* is the probability of observing a read at base position *i* within the element, and *L* is the set of base positions that have at least one read. To ensure reliable peak estimation, we required at least 30 reads on each strand. Peaks on a given strand with SI > -1.5 were classified as “peaked” (P); all others were classified as “broad” (B). Elements were further classified as P-P, P-B, or B-B based on peak shapes on both strands.
ii. The directionality index (DI) is defined as the difference in read counts between the two strands normalized by the total read counts, 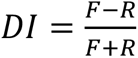 where *F* and *R* denote reads on the forward and reverse strands, respectively.
iii. disTSS refers to the distance between two prominent TSSs on opposite strands.

### ENCODE-rE2G promoter-enhancer (P-E) connections

We obtained ENCODE-rE2G predictions of P-E interactions for each tissue type of interest from the ENCODE portal. Distal PRO-cap elements were mapped to tested elements in the “element gene links” files (1-bp overlap; if multiple elements overlap, the one with the largest overlap is selected; elements with zero overlap are discarded). Proximal PRO-cap elements were mapped to tested elements in the “element gene links” files with the “isSelfPromoter” column set to True and assigned the corresponding gene symbol. The number of target genes for a given distal element and the number of enhancers regulating a given gene are calculated based on the “thresholded element gene links” files.

### Pairwise comparison of metastatic cancer samples and normal tissue samples

We included 93 normal tissues samples from 15 tissue types (highlighted in Fig. 1b), and 23 metastatic cancer samples. PCA was performed using normalized PRO-cap signals at divergent distal and proximal elements identified in these samples. Principal components explaining 90% of the variance were extracted to calculate Pearson correlations between metastatic cancer samples and normal tissue samples from the corresponding primary site, metastatic site, and other sites.

### Predicting the primary site of metastatic tumor samples

The training dataset consists of the 93 normal samples from 15 tissue types (Fig. 1b). Tissue-specific elements, obtained as described in the “Get tissue-specific and non-tissue-specific elements” section, were combined into a union set of features, excluding those found in multiple tissues. A linear support vector classifier was trained on this feature set with default hyperparameters and evaluated on metastatic samples with known primary sites.

### Statistical tests and multiple testing corrections

For statistical testing not described above, we used the two-sided Mann-Whitney U test to assess the differences between groups and reported BH-adjusted p-values (padj). Unless otherwise noted in the figure legend, significance was annotated using the following thresholds: ns, non-significant; *, 0.01 < padj ≤ 0.05; **, 0.001 < padj ≤ 0.01; ***, 0.0001 < padj ≤ 0.001; ****, padj ≤ 0.0001.

### Schematics

All schematics in Fig. 1, Fig. 6, and Extended Data Fig. 1 were created using BioRender, with the appropriate publication licenses.

## Data availability

The PRO-cap datasets generated in this study have been deposited in the ENCODE portal with accession codes listed in Supplementary Table 1. The visualization of BigWig and BigBed files is provided at the UCSC trackhub (https://genome.ucsc.edu/s/cybluetree/TRE%20hub).

## Code availability

All code used in this study has been deposited at https://github.com/haiyuan-yu-lab/TRE_landscape.

## Acknowledgments

We would like to thank all members of the ENCODE consortium for generating the other datasets used in this paper. We are grateful to Brad Bernstein at Dana-Farber and Nina Farrell at the Broad Institute for providing the ESC/IPSCs and their differentiated cell lines. We also thank Kyle Orwig from the University of Pittsburgh for providing human testis tissue samples. We thank all the members of our labs for their constructive feedback. This work was supported by the National Institutes of Health grants UM1HG009393, 5RM1GM139738 and 1R01HG012970 (to J.T.L. and H.Y.) and UM1HG009442 (to M.P.S). S.R.S was supported in part by a Cornell University Center for Vertebrate Genomics Scholar Award and Breast Cancer Coalition of Rochester Research Fellowship.

---

**Extended Data Figure 1.**
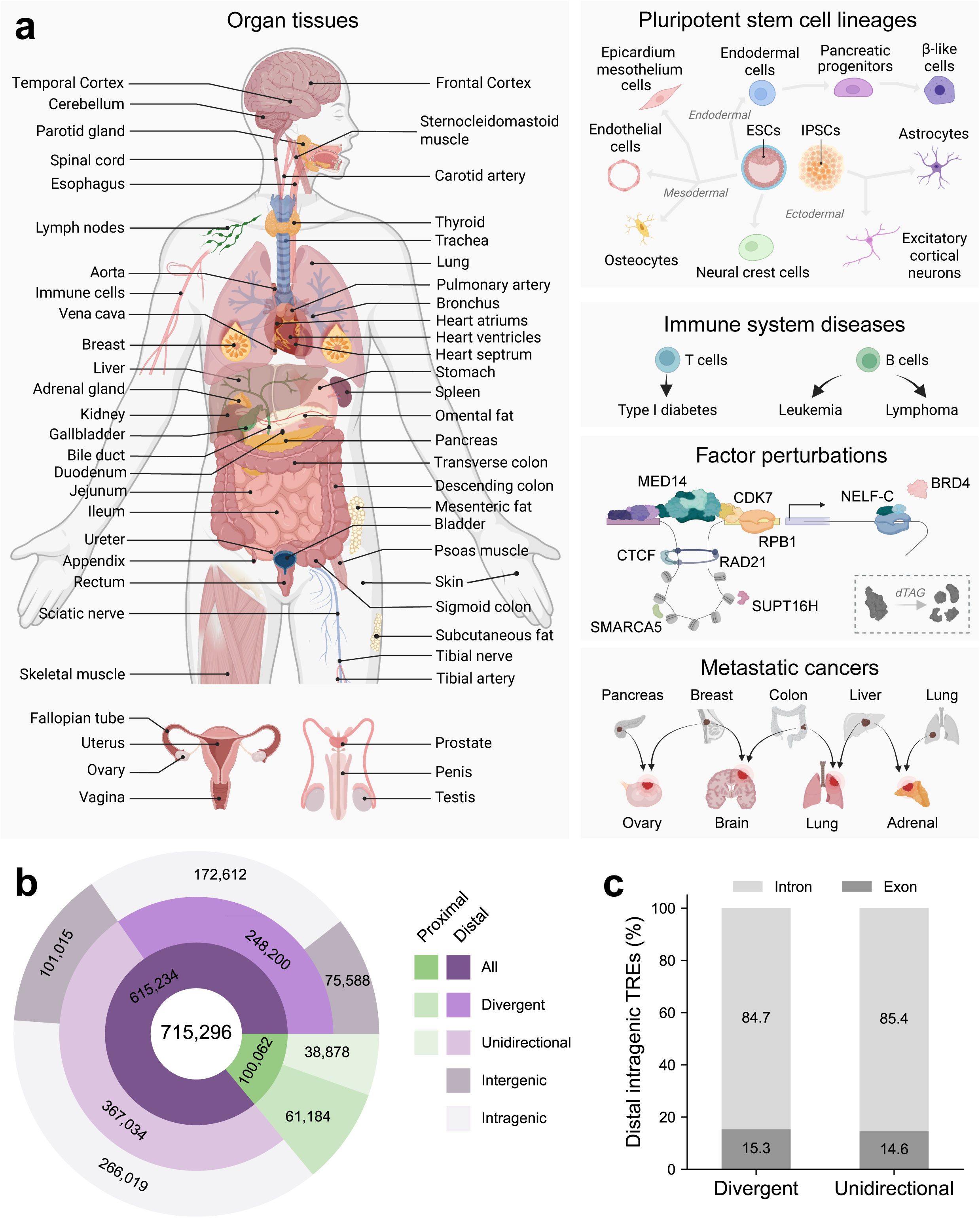
Comprehensive mapping of active transcriptional regulatory elements across human physiology and pathology. **(a)** Schematic depicting the diversity of samples profiled (n = 215), spanning tissues and cell types across all major human organ systems, pluripotent stem cells and their differentiated lineages, and a variety of disease states, as well as cells subjected to degron-mediated perturbation of targeted factors indicated in the schematic. **(b)** Donut plot showing the total number of TREs identified, classified by transcriptional directionality (divergent or unidirectional), distance to annotated gene TSSs (proximal or distal), and genomic context (intergenic or intragenic). **(c)** Barplot showing the proportion of divergent and unidirectional distal intragenic TREs, stratified by genomic context (intronic vs. exonic).

**Extended Data Figure 2.**
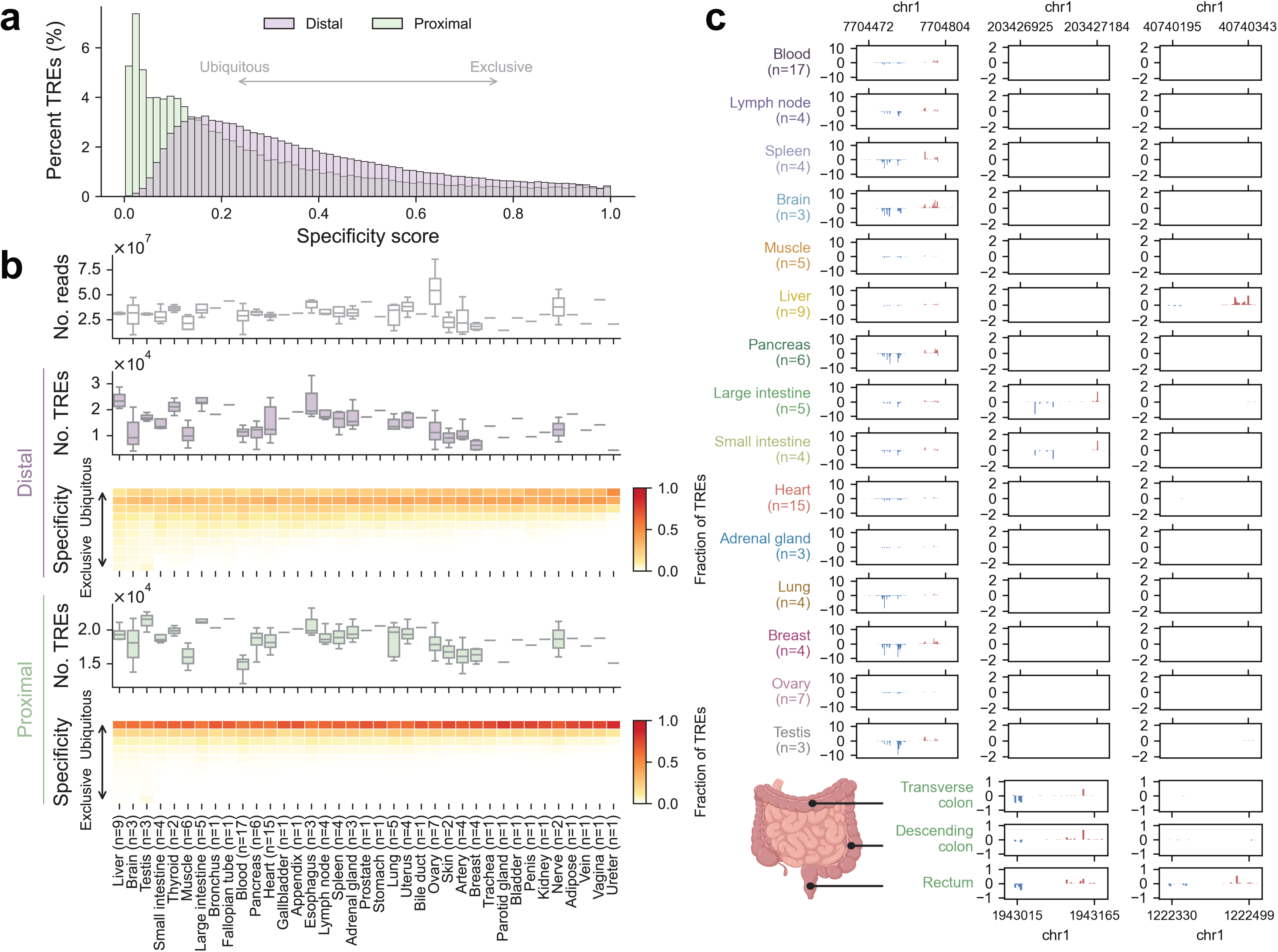
Tissue-specific TRE expression patterns across the human body. **(a)** Distribution of specificity scores for divergent distal (purple) and proximal (green) TREs. Scores represent a spectrum of tissue specificity, with higher values indicating more tissue-restricted activity and lower values reflecting more ubiquitous activity across tissues. **(b)** Summary of TRE specificity in each tissue type. *Top panel:* Number of deduplicated uniquely mapped reads per sample. *Second panel:* Number of detected divergent distal TREs per sample. *Third panel:* Expression specificity of divergent distal TREs, with colors indicating the fraction of elements per tissue (columns) falling into 10 specificity bins (rows; e.g., 0-0.1, 0.1-0.2, etc.). Fourth and *fifth panels:* Same as second and third panels, respectively, but for divergent proximal TREs. The number of profiled samples is indicated alongside each tissue type label. **(c)** *Upper panel:* Representative browsershots of PRO-cap signals from 15 tissue types across three genomic loci. Each track represents average RPM-normalized 5’ PRO-cap signals mapped to the plus (red) and minus (blue) strands across samples of the same tissue type. *Lower panel:* Representative browsershots of 5’ PRO-cap signals from three segments of the large intestine (transverse colon, descending colon, and rectum) across two genomic loci.

**Extended Data Figure 3.**
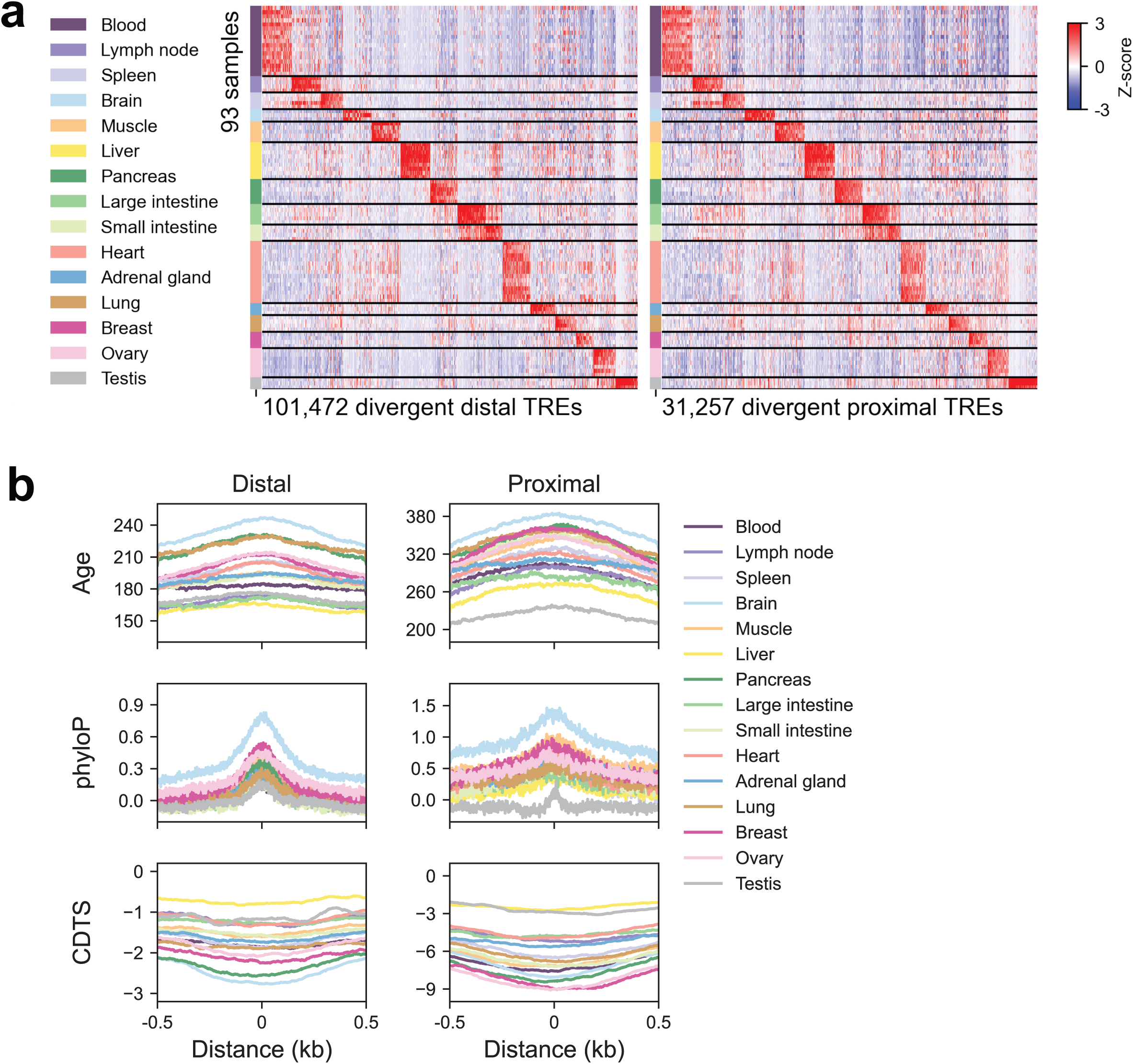
Evolutionary features of tissue-specific TREs. **(a)** Heatmap of PRO-cap signals for tissue-specific divergent distal and proximal TREs (columns) across the samples (rows). Tissue types are color-coded along the sample axis, consistent with Fig. 1B, and the heatmap colors represent Z-scores of each TRE across samples. **(b)** Metaplots of sequence age (million years ago), phyloP, and CDTS across tissue-specific divergent and proximal TREs. Tissue types are color-coded consistently with (a), and distances are shown as ±0.5 kb from the element center.

**Extended Data Figure 4.**
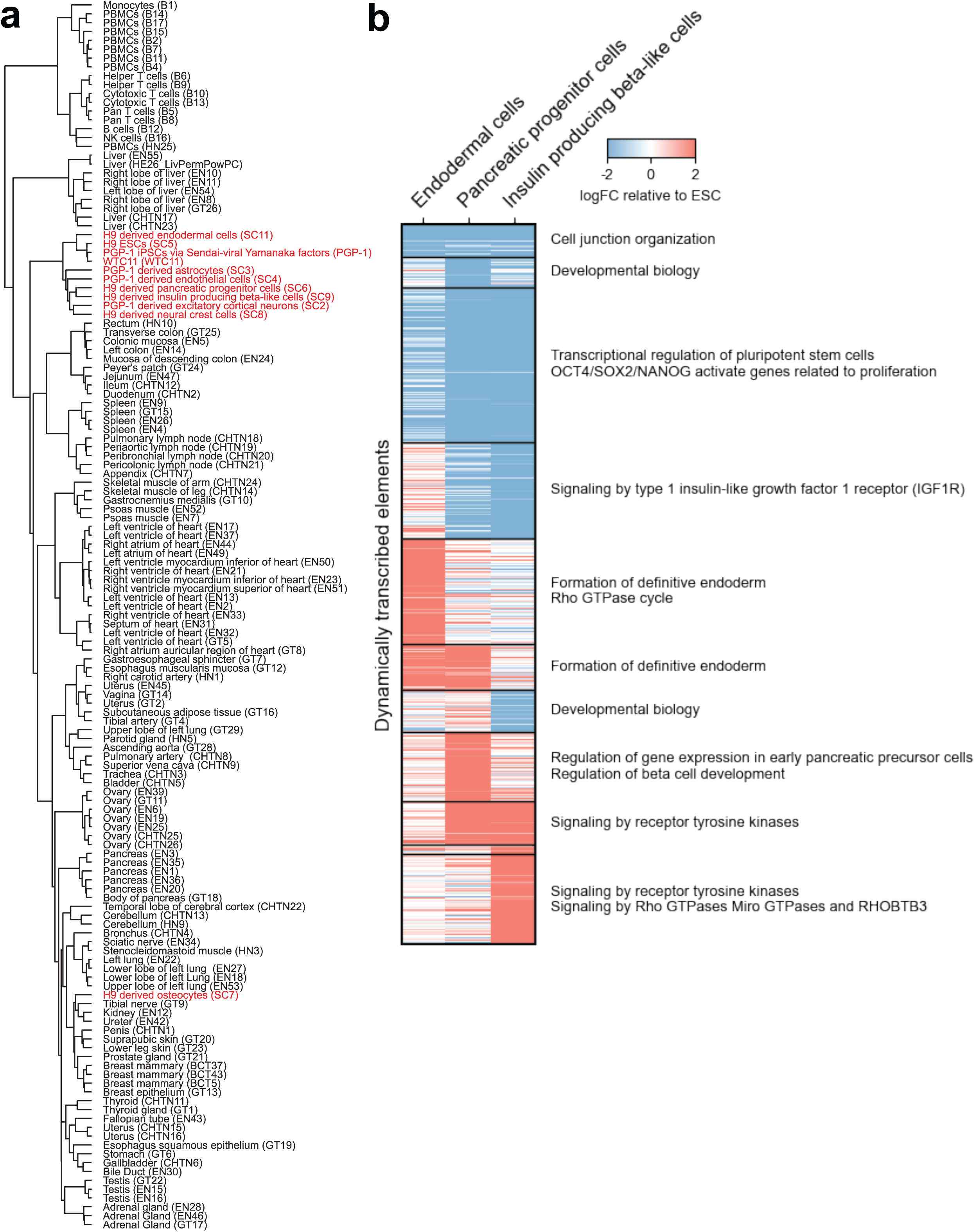
Lineage-specific transcriptional regulatory element usage during pluripotent cell differentiation. **(a)** Dendrogram showing hierarchical clustering of iPSCs/ESCs and their differentiated lineages along with normal tissues based on normalized PRO-cap expression levels of divergent distal TREs. **(b)** Heatmap of PRO-cap signals (relative to ESC) for divergent distal elements differentially expressed during pancreatic lineage differentiation, grouped by distinct transcriptional trajectories. Enriched Reactome canonical pathways of putative target genes are shown alongside each trajectory.

**Extended Data Figure 5.**
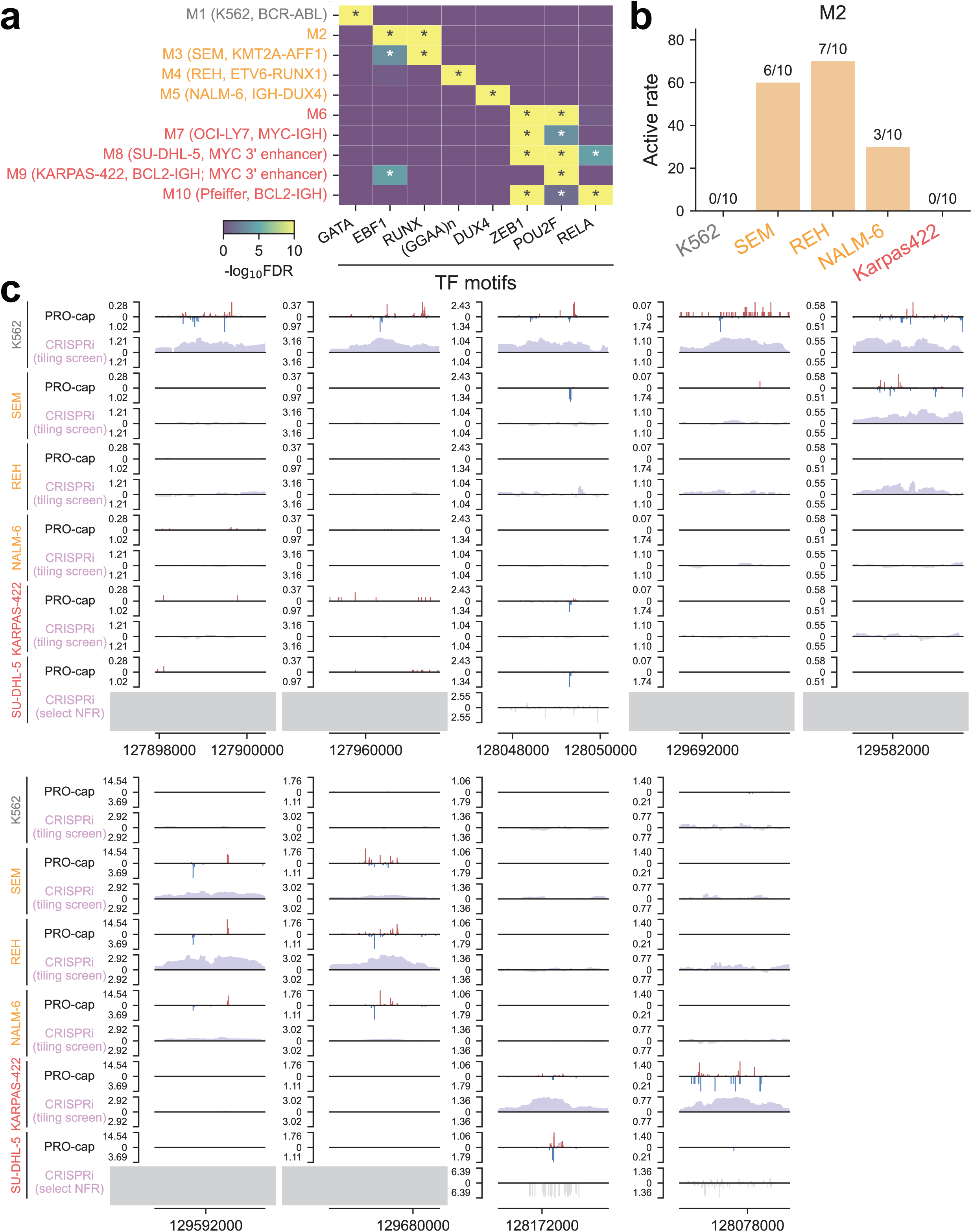
Subtype-specific transcriptional landscape of leukemia and lymphoma. **(a)** Heatmap of top *de novo* TF motif enrichment across subtype-specific TRE modules from Fig. 1f. Colors indicate –log10(BH-adjusted p-value) of motif enrichment. Cell line names and key pathogenic mechanisms (e.g., fusion proteins and overactivation of the MYC 3′ enhancer) are labeled alongside the corresponding TRE module. **(b)** Active rate (%) of divergent distal elements in TRE module M2 based on CRISPRi tiling screen at the *MYC* locus tested in five different cell lines representing CML, B-ALL, and GCB-DLBCL subtypes. **(c)** Genome browser tracks of 5’ PRO-cap signal and CRISPRi scores for distal elements in the *MYC* locus across different cell lines, with each example covering a 2.5-kb region on chromosome 8. In SU-DHL-5, CRISPRi guides were restricted to selected nucleosome-free regions (NFRs), and gray boxes indicate regions that were not targeted in the screen.

**Extended Data Figure 6.**
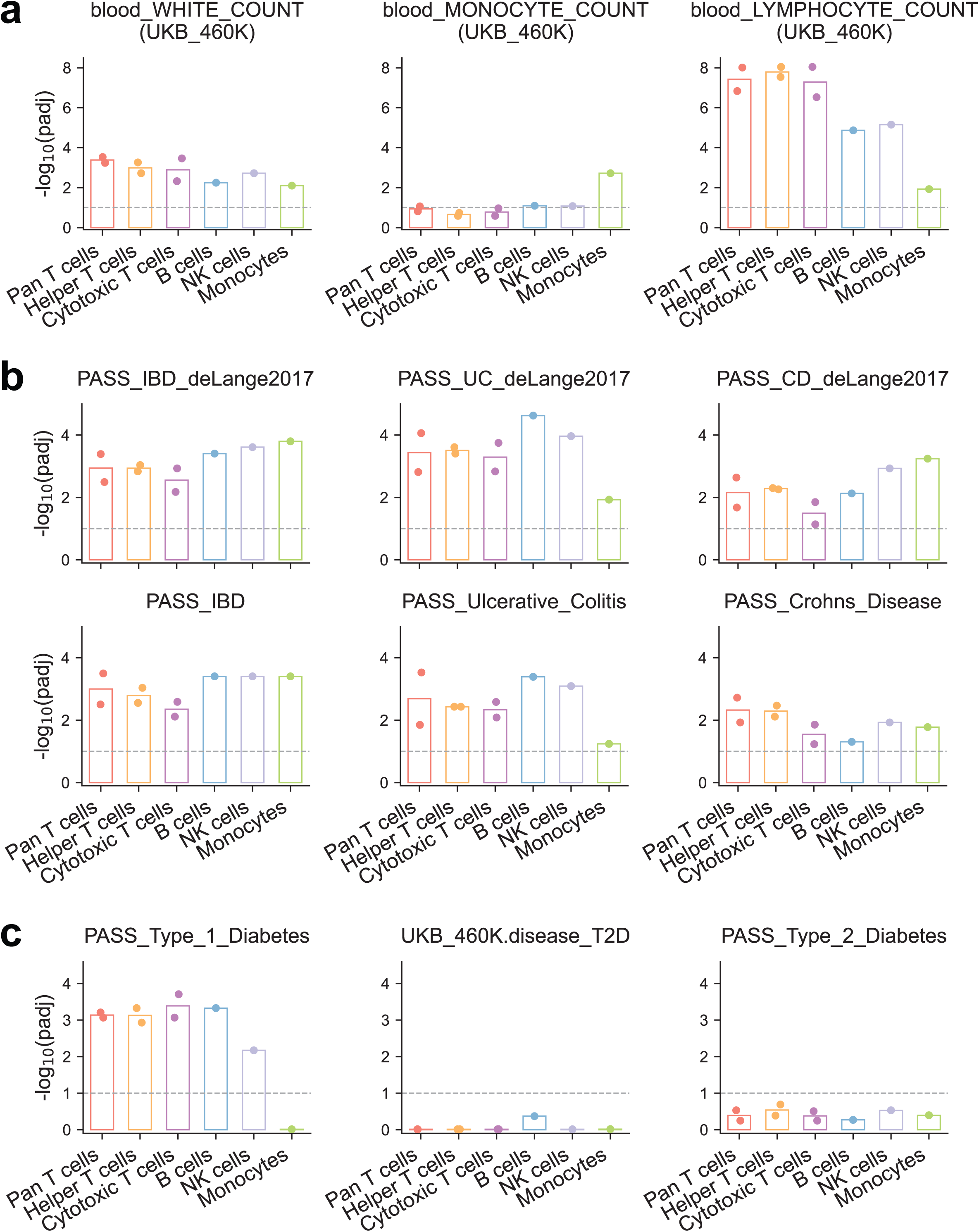
Enrichment of immune-related phenotypes in diverse blood cell types. **(a)** The significance of enrichment for blood cell count traits across major immune cell types from healthy individuals. **(b)** Same as (a) but for IBD, UC, and CD. **(c)** Same as (a) but for T1D and T2D. Each point represents an individual sample. The dashed line denotes the BH-adjusted p-value threshold of 0.1. IBD: Inflammatory bowel disease; CD: Crohn’s disease; UC: Ulcerative colitis; T1D: Type 1 diabetes; T2D: Type 2 diabetes. GWAS summary statistics were obtained from the curated collection of the Alkes Price lab (see Methods).

**Extended Data Figure 7.**
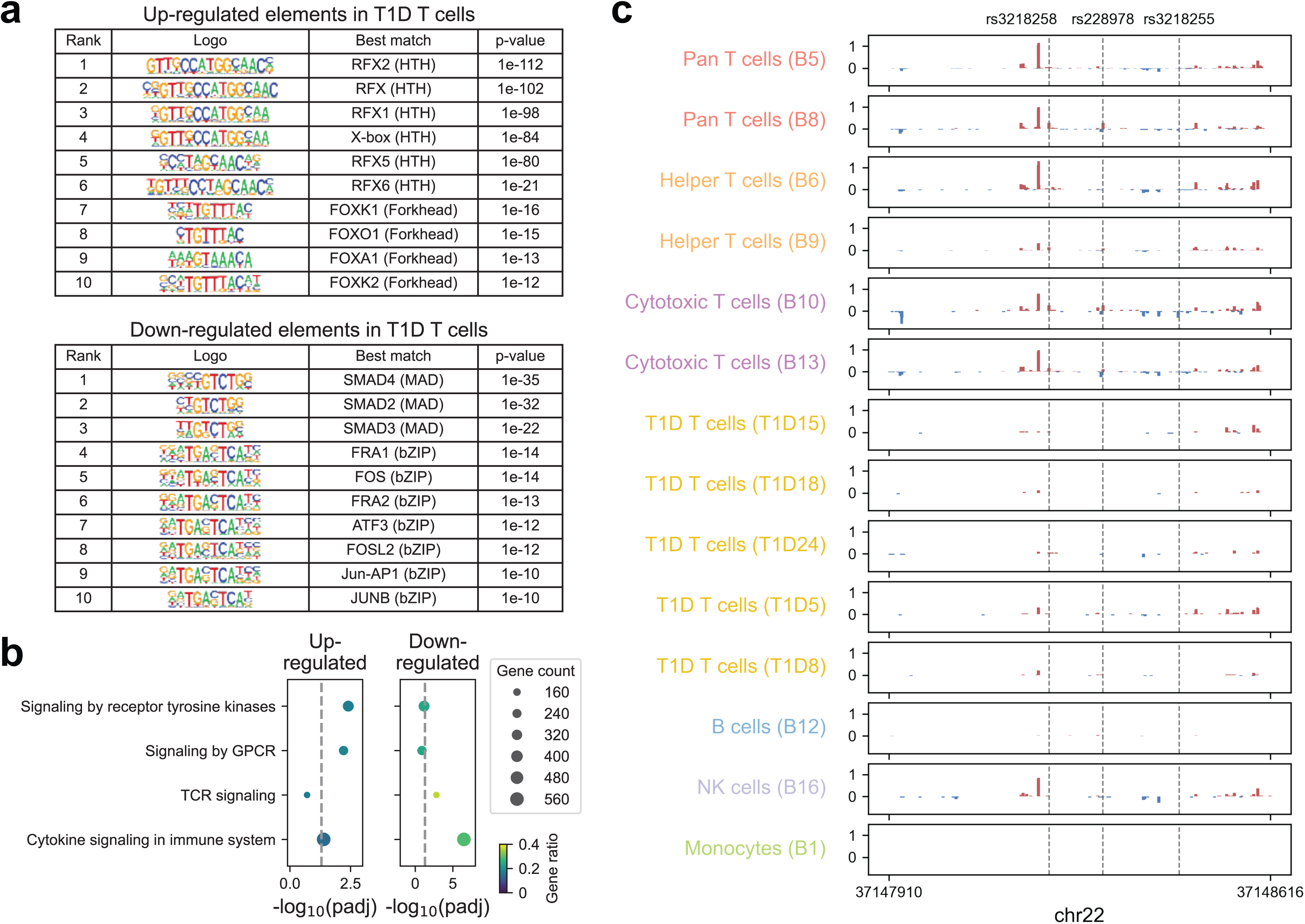
Functional characterization of differentially expressed transcriptional regulatory elements in T1D patient-derived T cells. **(a)** Enrichment of TF binding motifs in differentially expressed TREs between T1D patient- and non-diseased donor-derived T cells. Top 10 significantly enriched motifs in upregulated and downregulated TREs are shown (BH-adjusted p-value < 0.05). Sequence logos with their corresponding TF identity match (TF name and DNA-binding domain) and p-value are shown. **(b)** Pathway enrichment analysis of genes linked to differentially expressed TREs in T cells from T1D patients versus controls. Dot size indicates the number of genes in each pathway (Reactome Canonical Pathways). Gene ratio represents the fraction of pathway genes that overlap with the input gene set. Dashed lines indicate BH-adjusted p-value threshold of 0.05. **(c)** Browser shot of PRO-cap signal tracks (RPM-normalized) at the *IL2RB* intronic enhancer locus in T cells from individual T1D patients and in different immune cell types from each non-diseased donor. Same locus as shown in Fig. 3f. Dashed lines indicate fine-mapped T1D-associated variants located within this region.

**Extended Data Figure 8.**
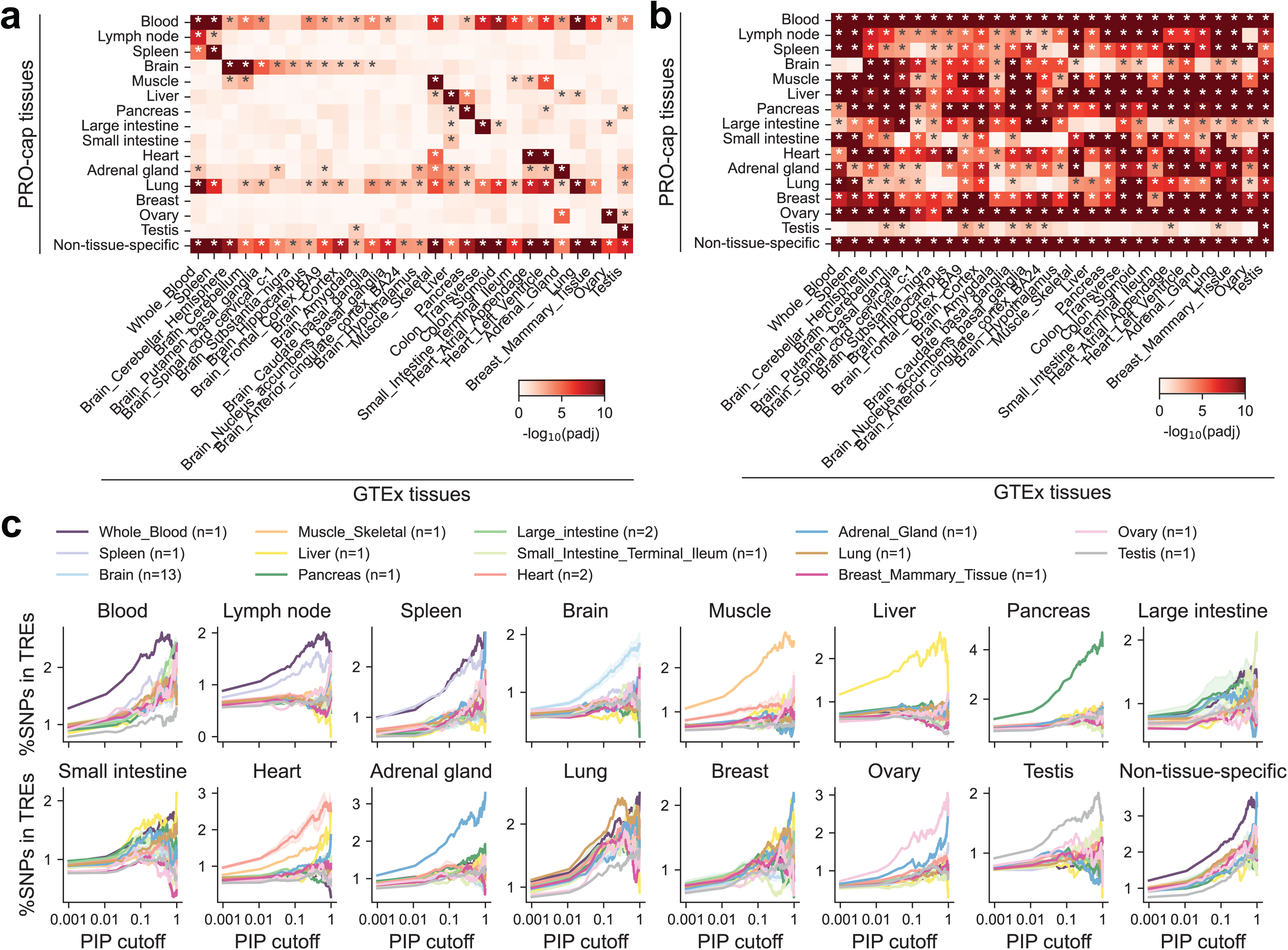
eQTL enrichment across transcriptional regulatory element annotations. **(a)** Heatmap showing the significance of eQTL enrichment across divergent distal tissue-specific and non-tissue-specific TRE annotations. Asterisks (*) denote BH-adjusted p-value < 0.05. **(b)** Same as (a), but for divergent proximal elements. **(c)** Percent of fine-mapped eQTL variants (above a given PIP threshold) overlapping tissue-specific and non-tissue-specific divergent distal elements. Lines show the average across related GTEx tissues; subplot titles indicate TRE categories. Traces are shown for thresholds with ≥5 variants. Shaded areas denote standard error.

**Supplementary Figure 1.**
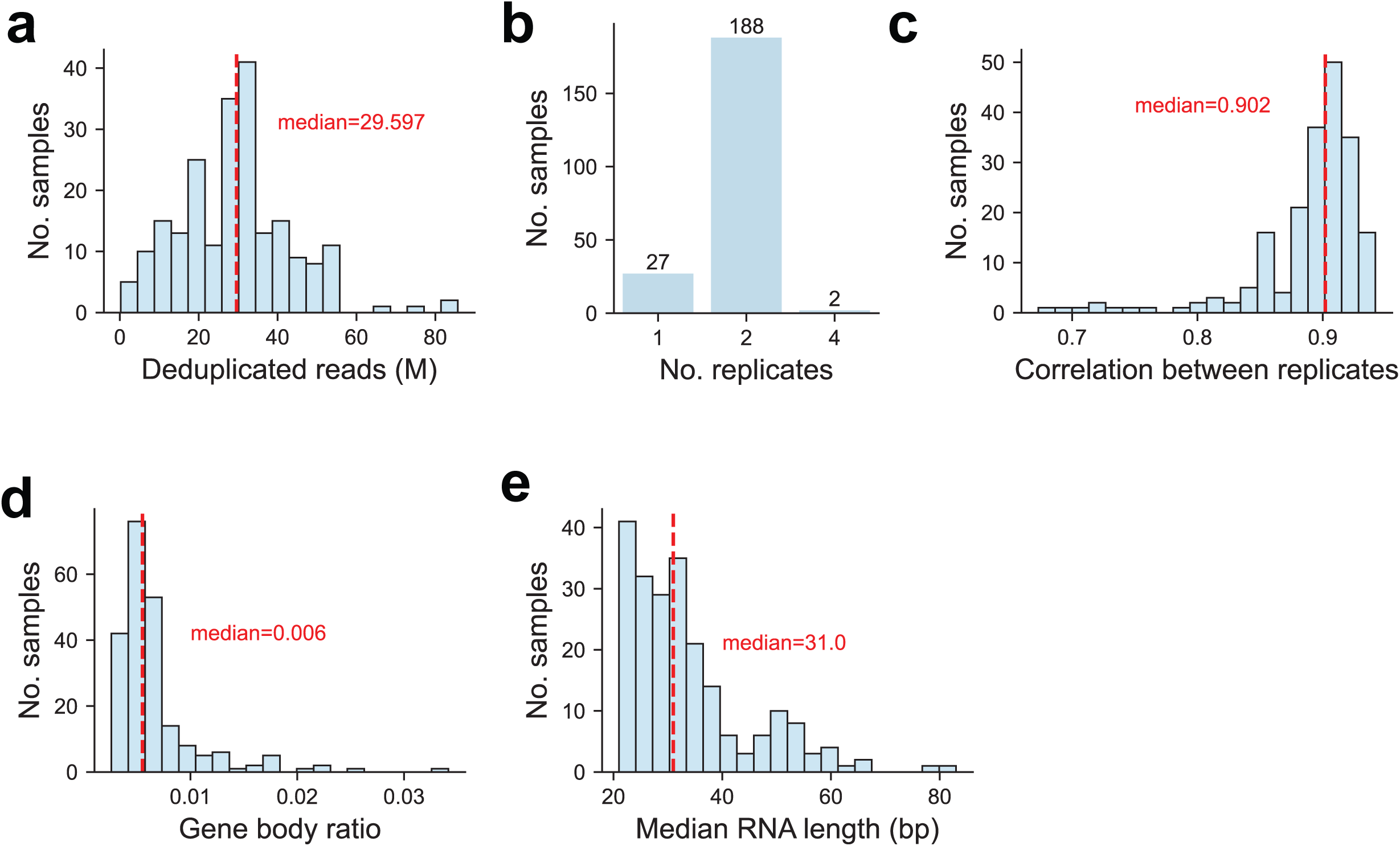
Overview of PRO-cap data quality. **(a)** Histogram of deduplicated uniquely mapped reads (in millions) across samples. These represent high-quality reads after adapter trimming, quality filtering, alignment to the genome, and removal of PCR duplicates. **(b)** Barplot showing the number of replicates of PRO-cap libraries generated per sample. **(c)** Histogram of Pearson correlations between replicates (samples with ≥2 replicates), computed across genome-wide 5’ PRO-cap signals. **(d)** Histogram of gene body ratios across samples. Ratios represent length-normalized PRO-cap signal within the gene body relative to the sum of length-normalized TSS and gene body signals, for highly expressed genes. **(e)** Histogram of median nascent RNA lengths (bp) captured by PRO-cap across samples. Lengths are derived from paired-end PRO-cap reads, reflecting the distance from the 5′ capped end to the 3′ pause site.

**Supplementary Figure 2.**
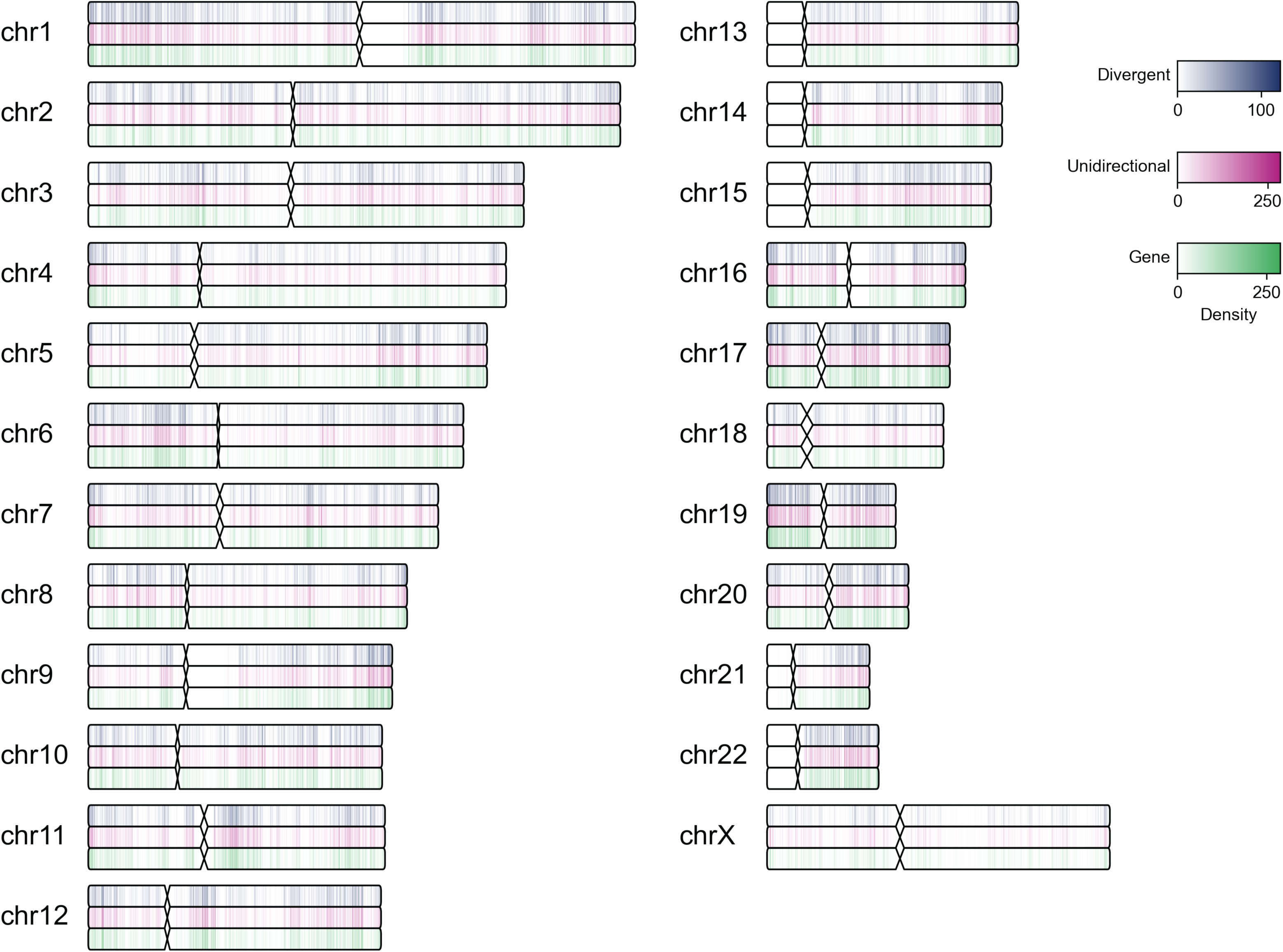
Genomic distribution of PRO-cap-detected TREs. Genomic distribution of TREs and genes across the genome. For each chromosome, the top track shows divergent TREs (purple), the middle track shows unidirectional TREs (pink), and the bottom track shows genes (green). Densities represent log10-transformed counts in 100 Mb bins and are displayed across the chromosomes.

**Supplementary Figure 3.**
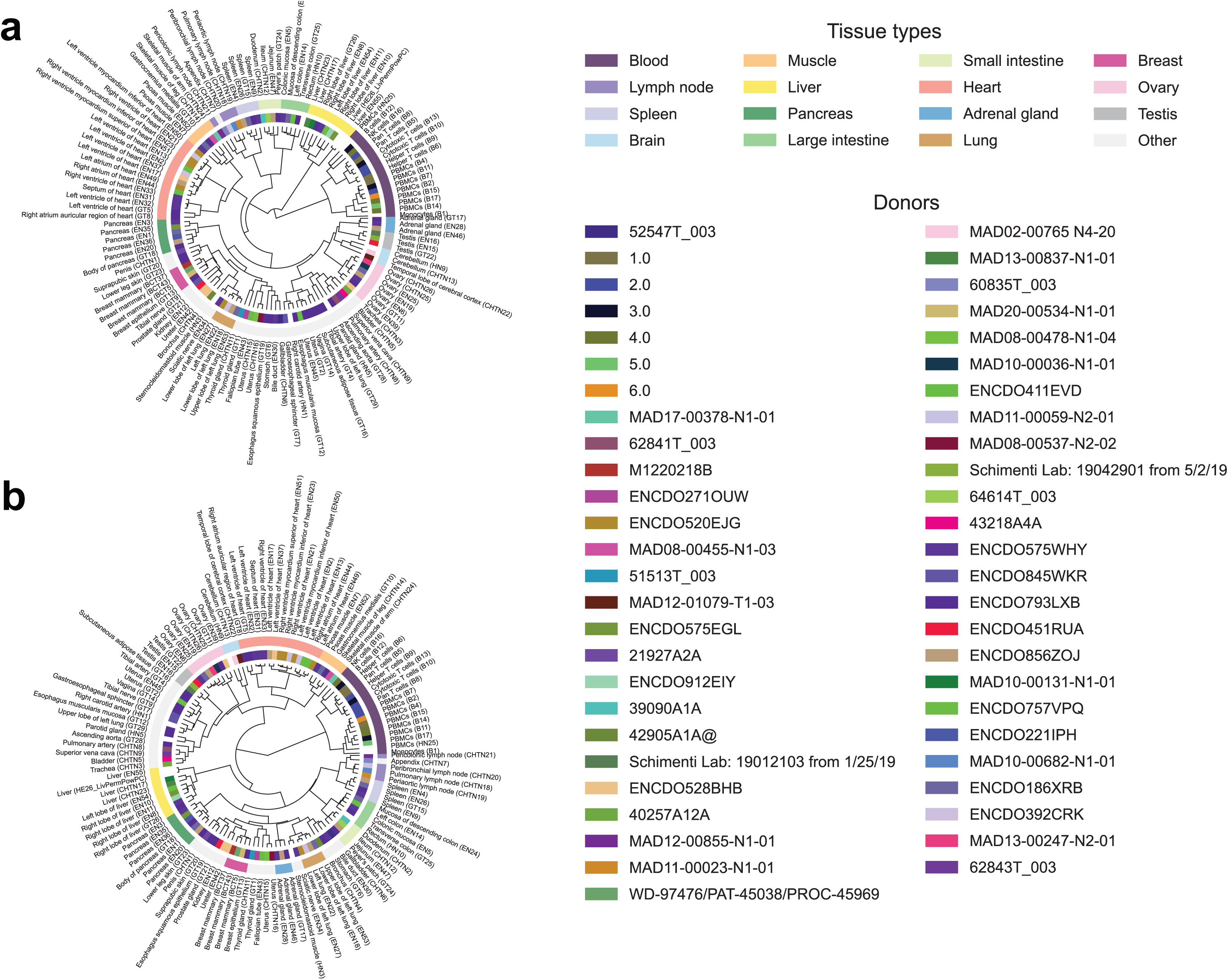
Hierarchical clustering of human tissue samples by divergent TREs. **(a)** Dendrogram showing hierarchical clustering of tissue samples based on normalized PRO-cap expression at divergent distal TREs. Sample names are labeled; subtrees primarily representing a single tissue type (≥3 samples) are highlighted. The outer circle indicates tissue type, and the inner circle indicates donor ID. **(b)** Same as (a), but for divergent proximal TREs.

**Supplementary Figure 4.**
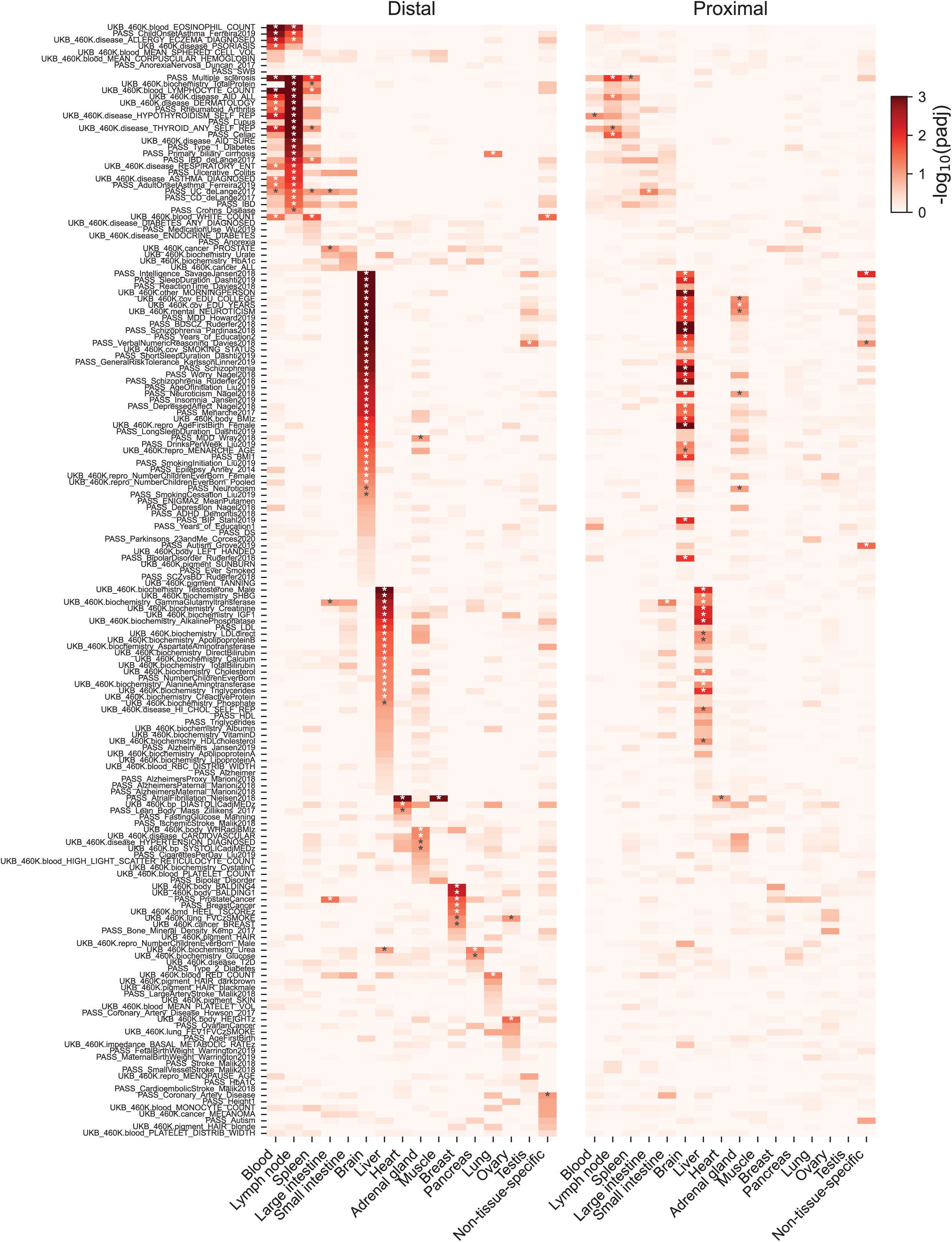
Tissue-specific effects of disease- and trait-associated variants. Heatmap showing the significance of enrichment in human diseases and complex traits from 176 studies across divergent distal (*left panel*; 7,943 tissue-specific elements per tissue type and 7,659 non-tissue-specific elements) and proximal (*right panel*; 2,540 tissue-specific elements per tissue type and 2,707 non-tissue-specific elements) TRE annotations. Asterisks (*) denote BH-adjusted p-values < 0.1. Only tissue types with at least 3 biological replicates were included.

**Supplementary Figure 5.**
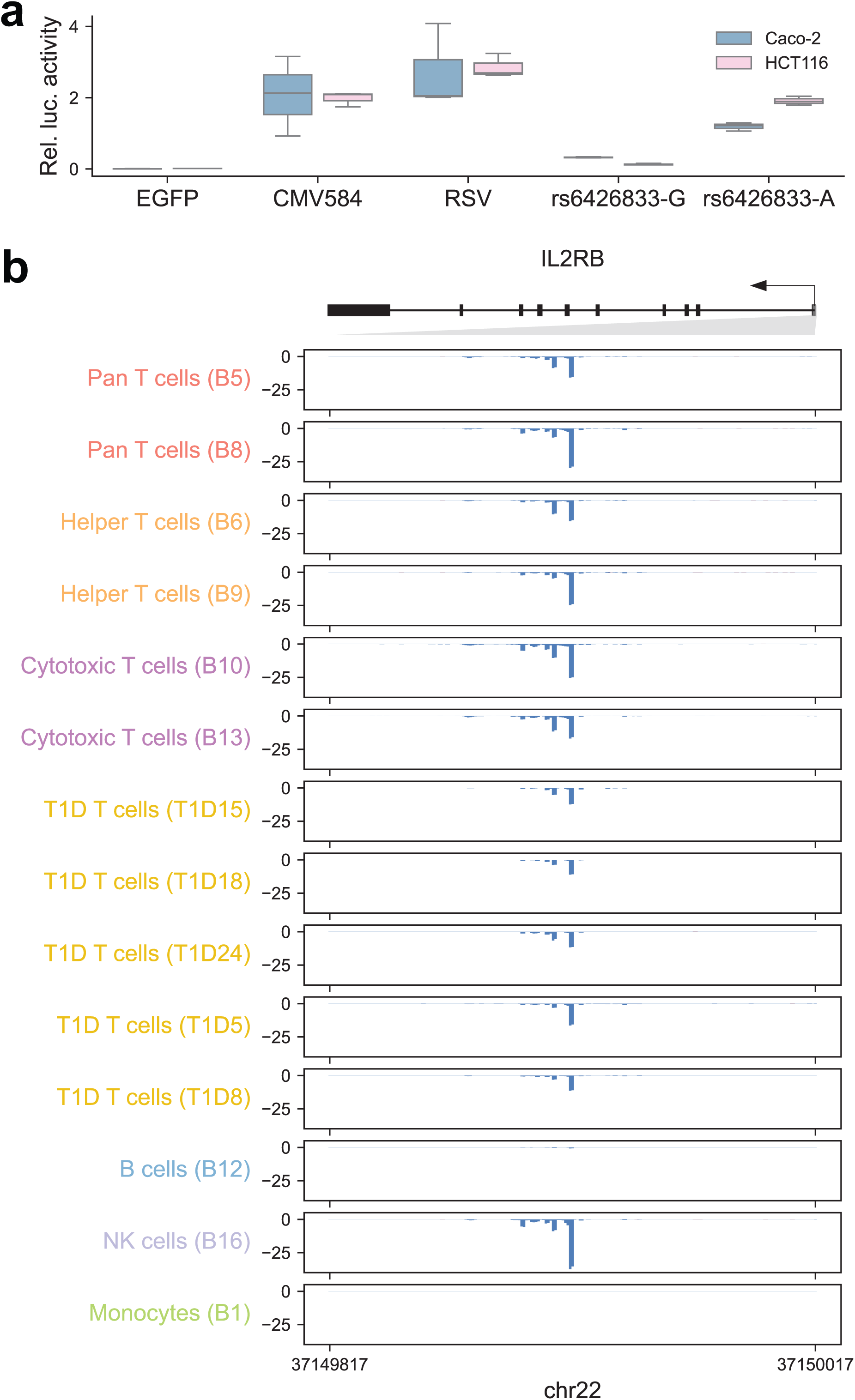
Disease-associated variants in TREs modulate regulatory activity. **(a)** Luciferase reporter assay showing enhancer activity of a negative control (EGFP), positive controls (CMV584, RSV), and two alleles (G vs. A) of rs6426833 in Caco-2 and HCT116 cells. Three biological replicates were included for each group. **(b)** Browser shot of PRO-cap signal tracks (RPM-normalized) at the *IL2RB* promoter locus in T cells from individual T1D patients and in different immune cell types from each non-diseased donor. The *IL2RB* gene is depicted with an arrow indicating transcription orientation and black rectangles marking exons. The grey triangle highlights the region shown in the zoomed-in panels.

**Supplementary Figure 6.**
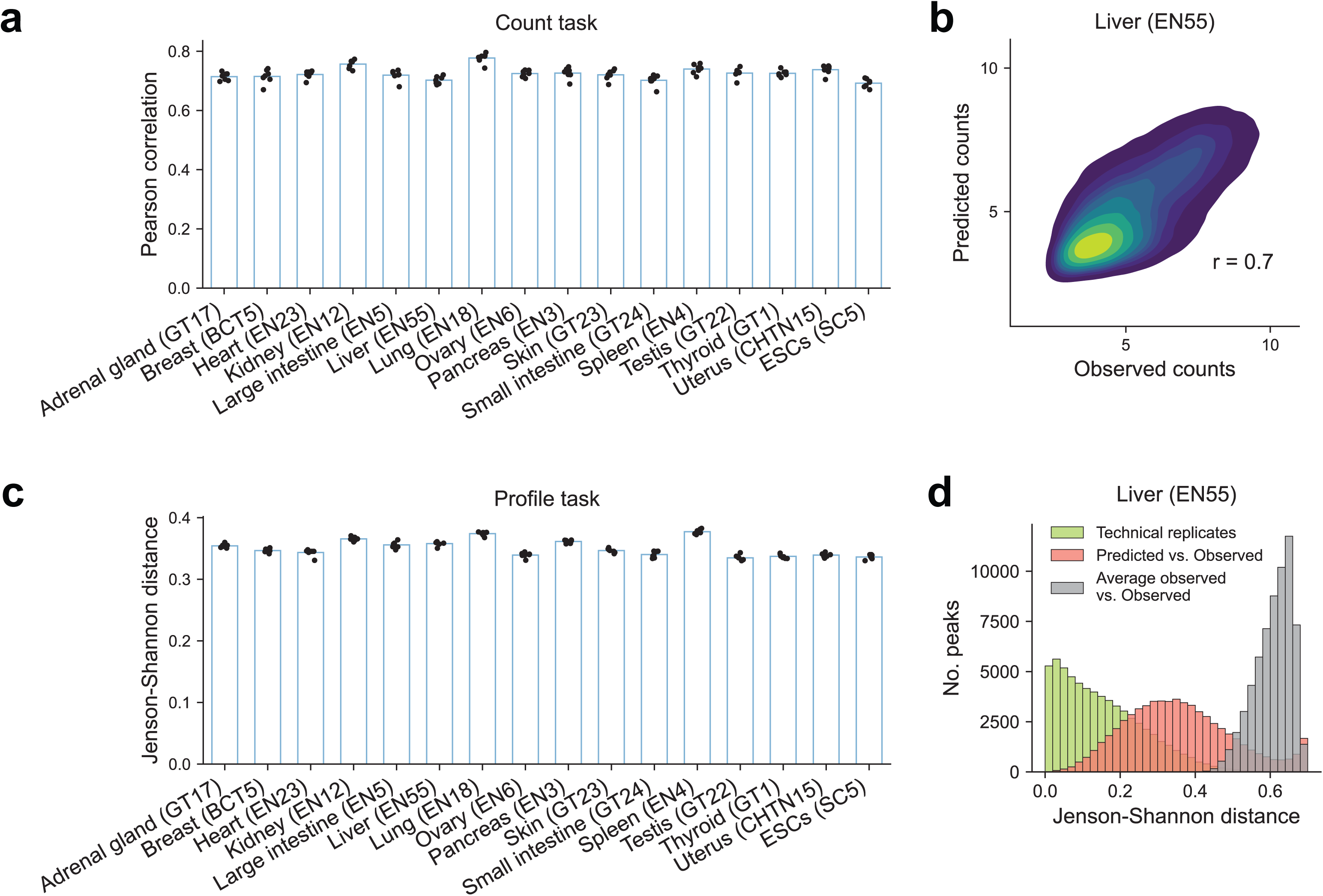
ProCapNet model performance across tissues. **(a)** Barplots showing the Pearson correlations between log-transformed observed and predicted total counts on held-out test chromosomes across 7-fold cross-validation for each model. Each datapoint represents one fold. **(b)** Density plot of log-transformed observed and predicted counts at PRO-cap peaks from held-out test chromosomes across 7-fold cross-validation, based on the model trained on the liver sample (EN55). **(c)** Same as (a), but showing the mean Jensen-Shannon distance between observed and predicted profiles for each model. **(d)** Distribution of Jensen-Shannon distances between observed and predicted base-resolution profiles at PRO-cap peaks from held-out test chromosomes across 7-fold cross-validation (red), based on the model trained on the liver sample (EN55). For comparison, distances between observed profiles from two technical replicates of EN55 are shown in green (upper bound), and distances between observed profiles and profiles averaged over all peaks are shown in grey (baseline).

**Supplementary Figure 7.**
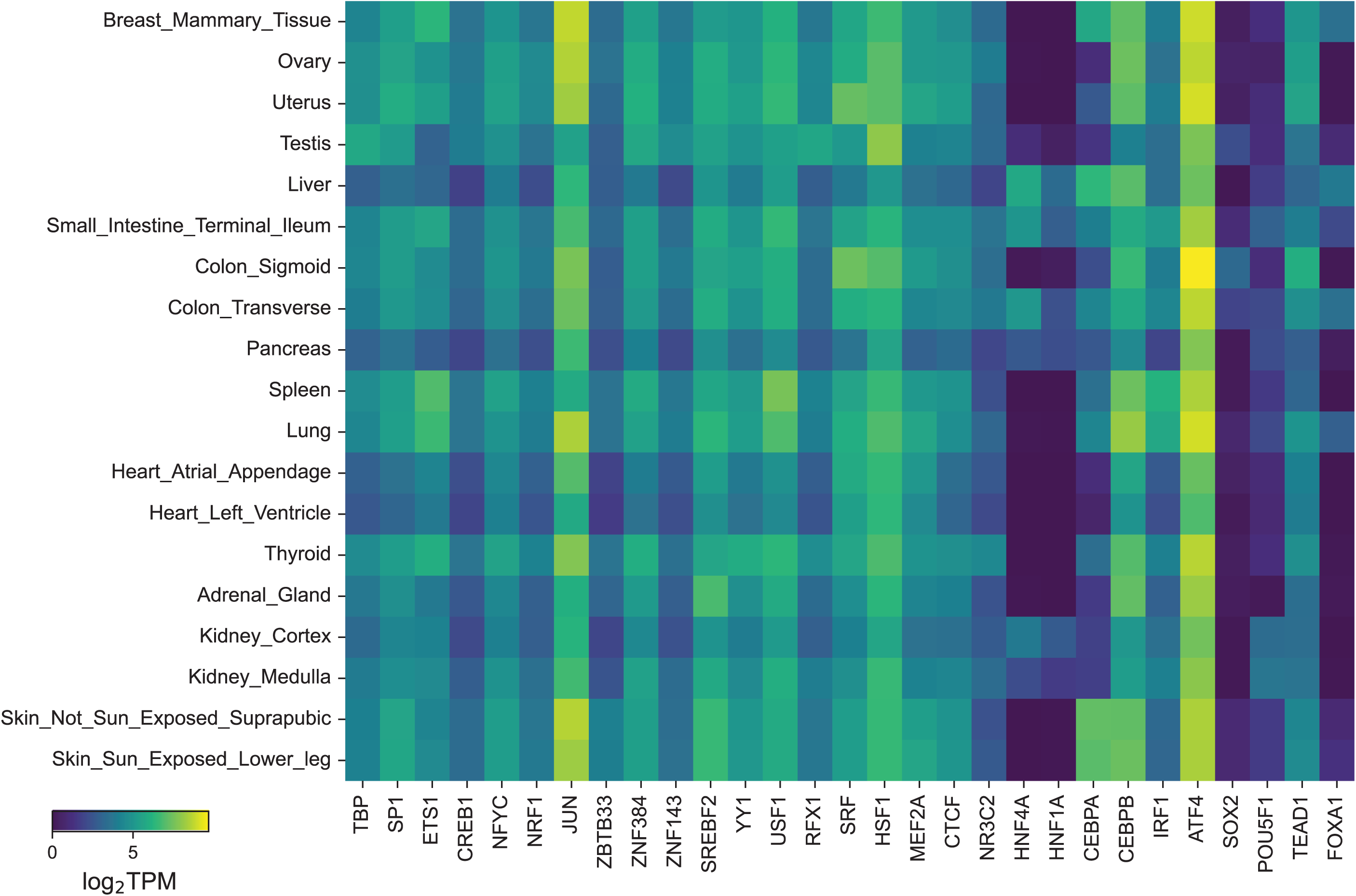
Expression patterns of TFs corresponding to motifs contributing to transcription initiation at TREs across tissue types. Expression levels (median gene-level log_2_ TPM per tissue from GTEx RNA-seq data) of representative TFs corresponding to motifs identified by ProCapNet as contributing to transcriptional strength (count task) and/or TSS positioning (profile task) across matched GTEx tissues.

**Supplementary Figure 8.**
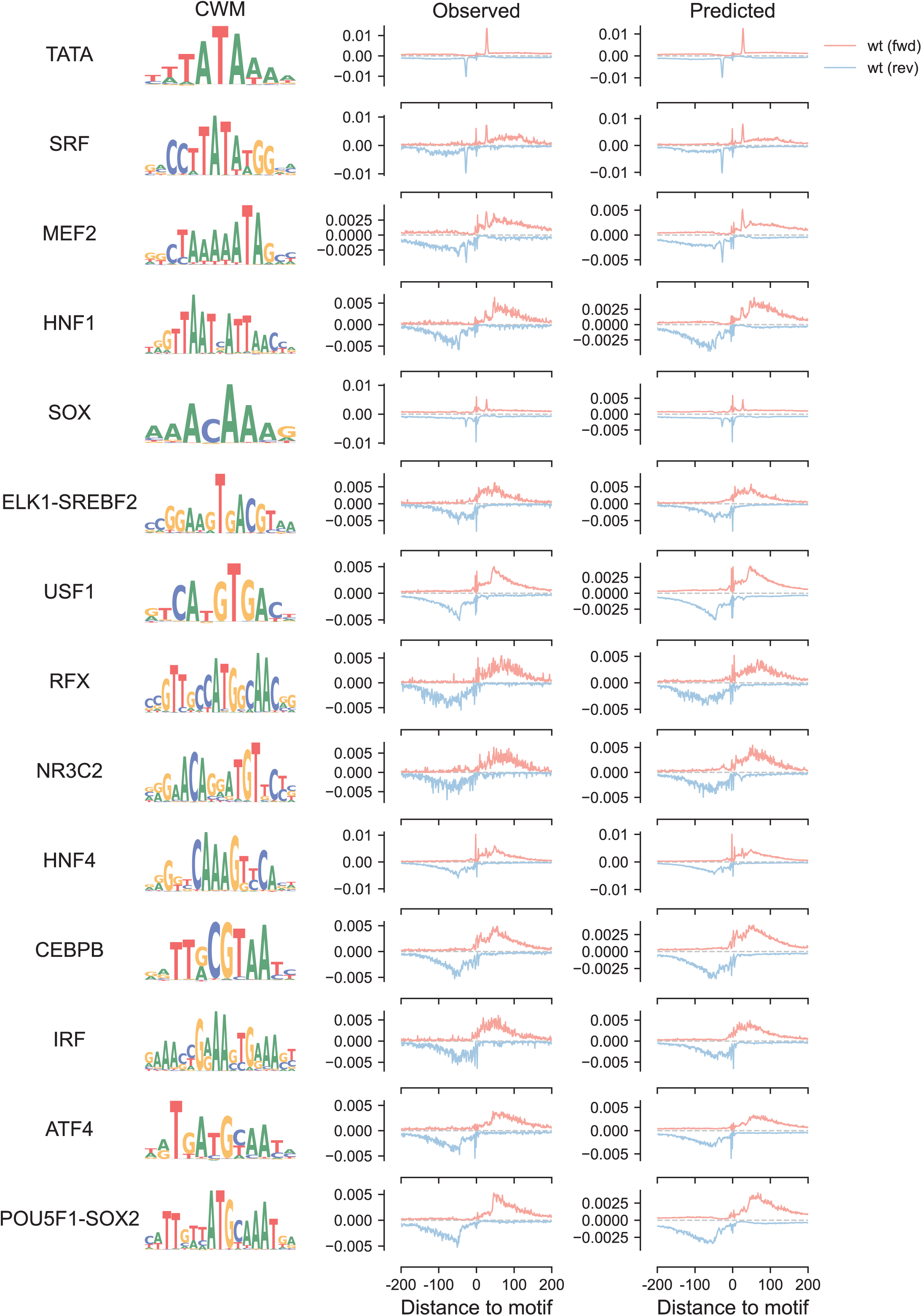
Distinct transcription initiation patterns associated with TF motifs at TREs. Representative TF motifs showing distinct effect curves on transcription profiles. *Left panel*: TF motif name and corresponding contribution weight matrix from the profile task of TF-MoDISco output. *Middle panel*: average observed PRO-cap profiles are centered at motif instances and aligned by motif orientation, with reverse-strand motifs flipped. *Right panel*: same as middle panel, but for predicted profiles. One representative ProCapNet model is shown for a given motif.

**Supplementary Figure 9.**
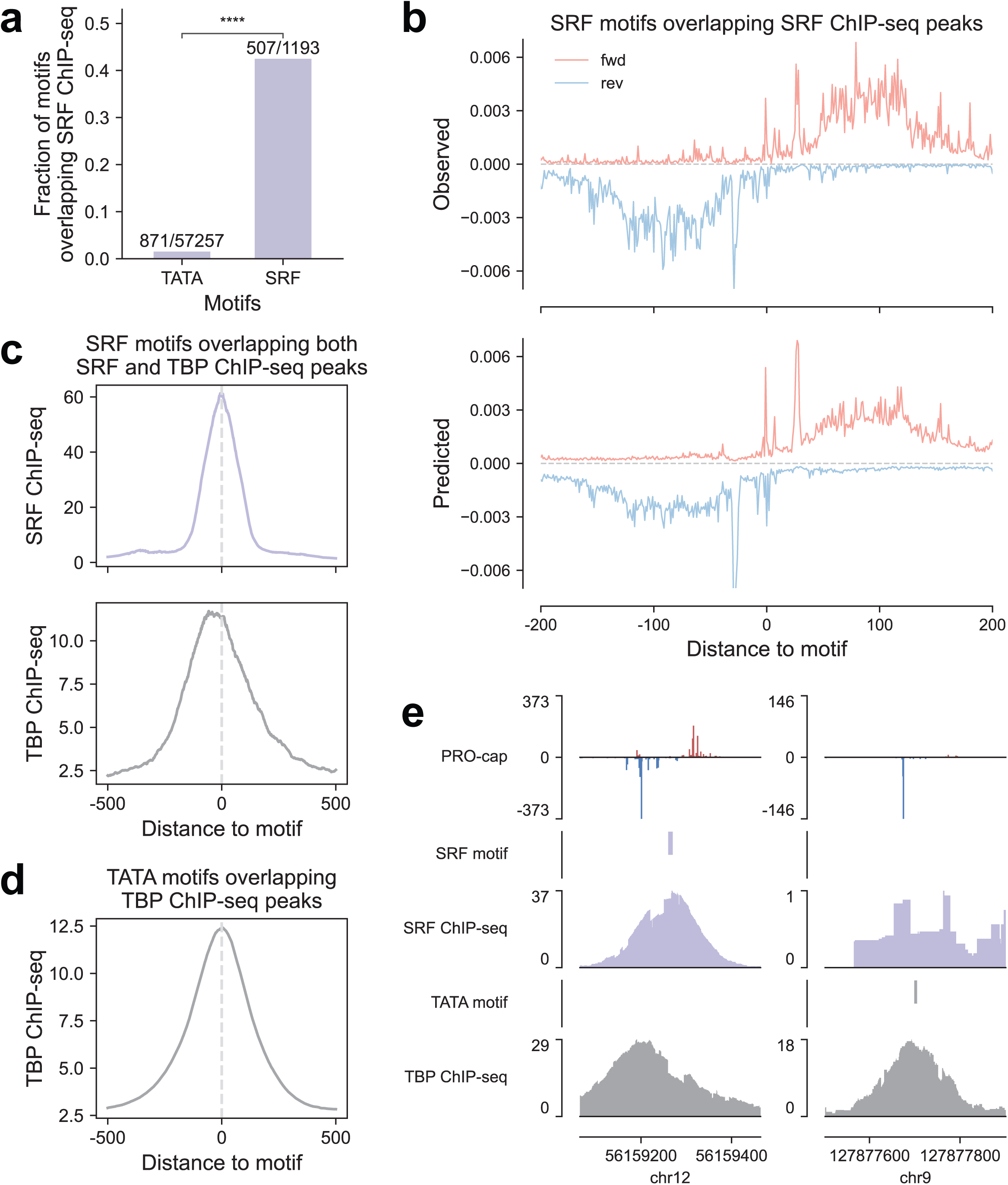
SRF motif enrichment and PRO-cap initiation profiles at ChIP-supported binding sites. **(a)** Fraction of TATA and SRF motifs overlapping SRF ChIP-seq peaks (ENCFF648QJE) in ESCs. **(b)** *Top*: observed PRO-cap profiles centered on SRF motif instances overlapping SRF ChIP-seq peaks, accounting for motif orientation. *Bottom*: average predicted profiles centered on the same SRF motif instances with motif orientation considered. **(c)** Metaplots of SRF (ENCFF920RYL) and TBP (ENCFF206PWF) ChIP-seq signal overlapping SRF motifs. Distances are shown as ±0.5 kb from the motif center. **(d)** Metaplot of TBP ChIP-seq signal overlapping TATA motifs. Distances are shown as ±0.5 kb from the motif center. **(e)** PRO-cap signal and ChIP-seq tracks for SRF and TBP at a representative locus. The SRF and TATA motif instances are indicated.

**Supplementary Figure 10.**
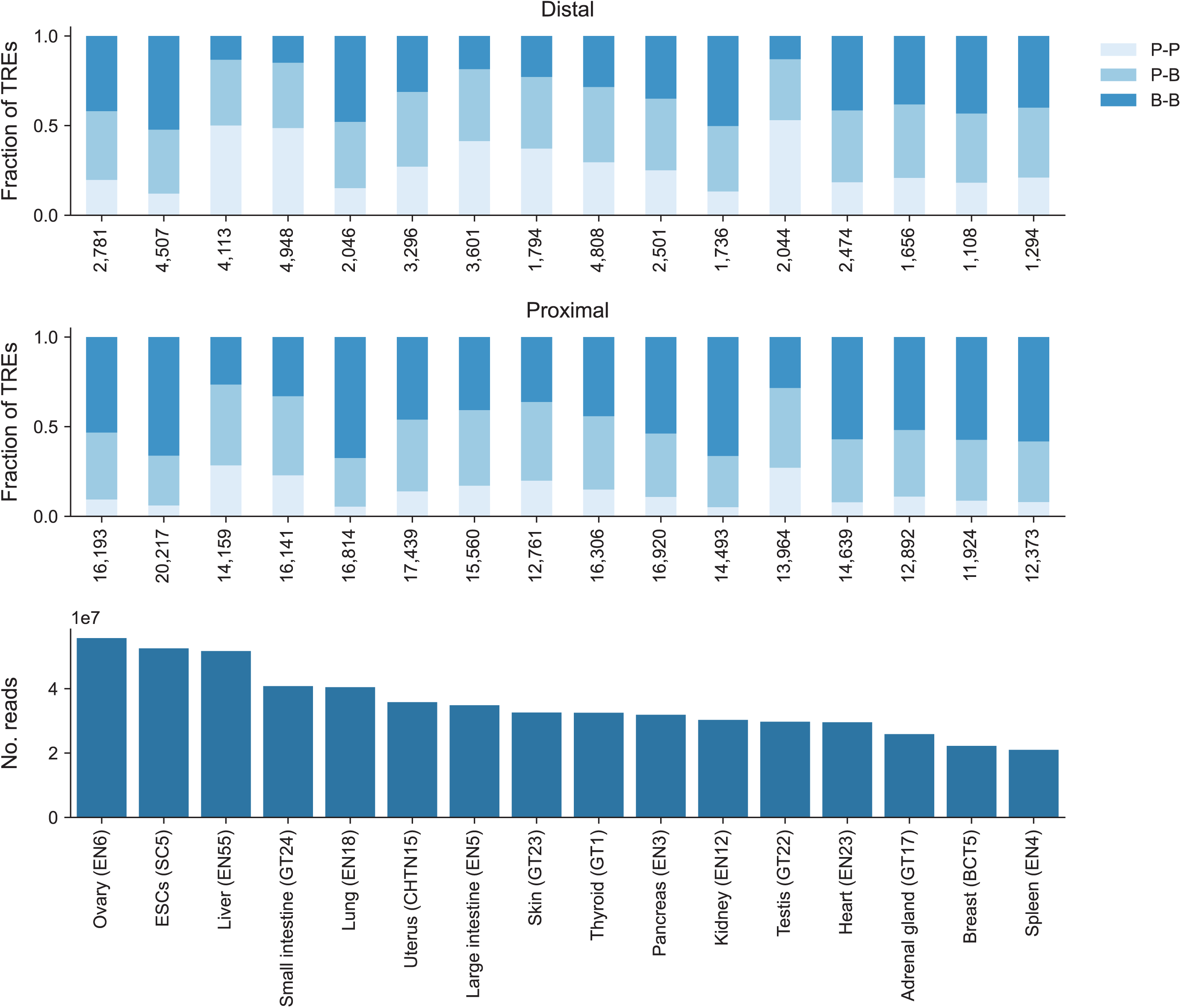
Distribution of TRE peak shape classes across tissue and cell types. Fractions of TREs classified as P-P, P-B, and B-B peak shapes across tissues and cell types (t*op:* distal elements; m*iddle:* proximal elements). *Bottom:* sequencing depth (number of uniquely mapped, PCR deduplicated reads) for each corresponding sample shown above.

**Supplementary Figure 11.**
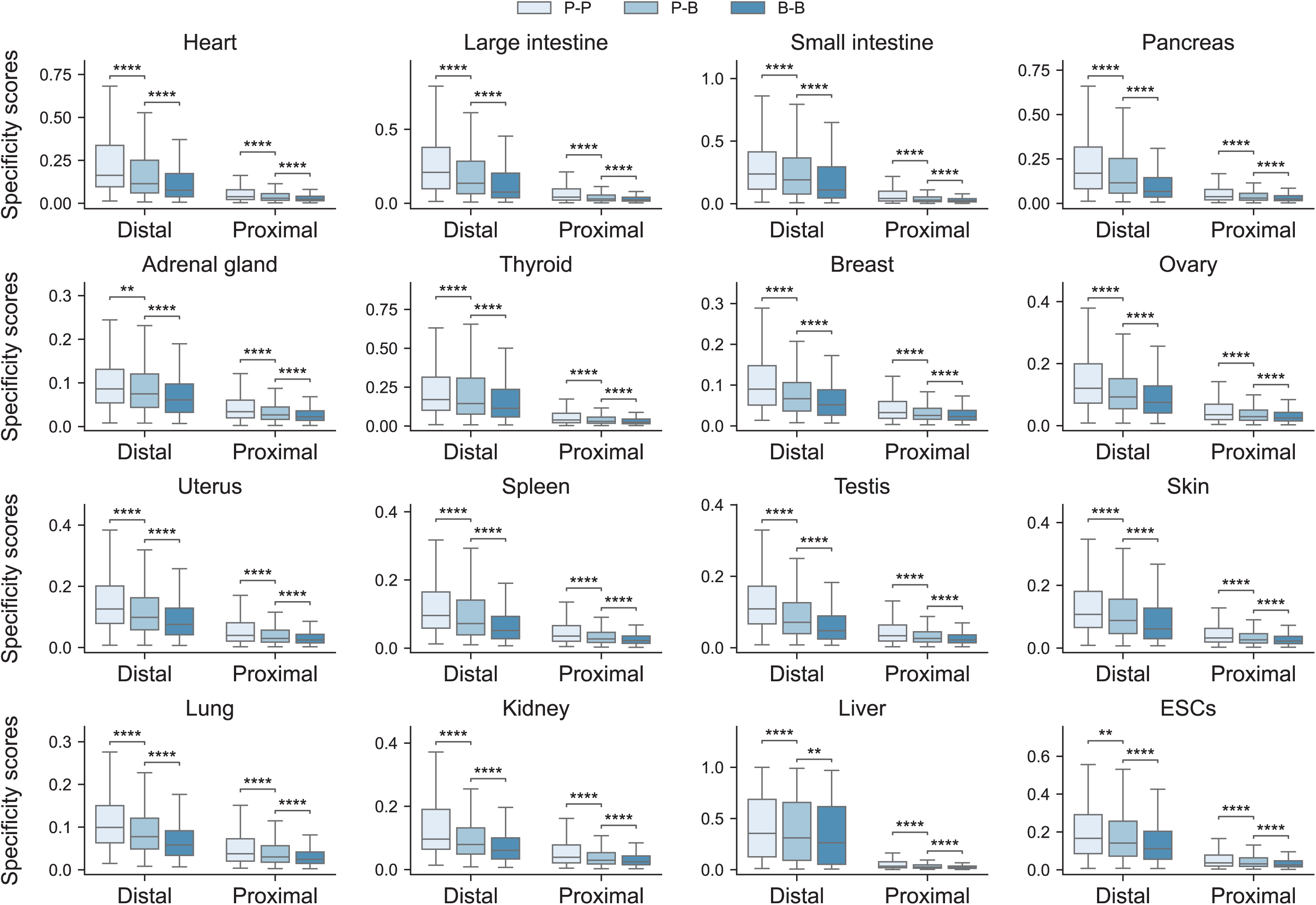
Specificity scores of TREs with different peak shapes across tissue and cell types. Boxplots showing tissue specificity scores for TREs with P-P, P-B, and B-B peak shapes, stratified by divergent distal and proximal elements across tissues and cells. Each point represents the specificity score of an individual TRE in the indicated biosample. **, 0.001 < padj ≤ 0.01; ****, padj ≤ 0.0001.

**Supplementary Figure 12.**
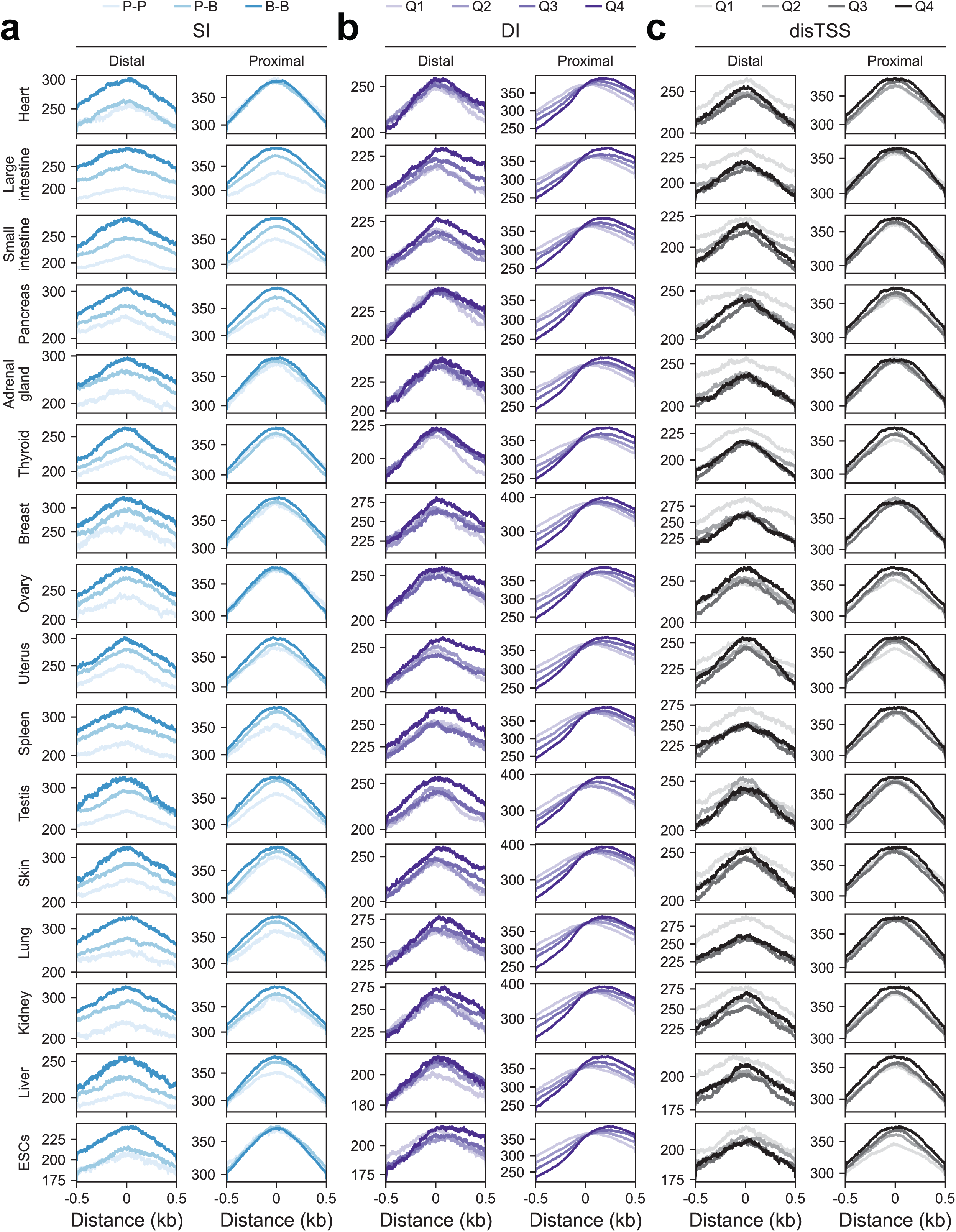
Sequence age of TREs with distinct initiation features across SI, DI, and disTSS categories. Metaplots of sequence age (million years ago) across divergent distal and proximal elements, grouped by SI (**a**), DI (**b**), and disTSS (**c**) categories. Distances are shown as ±0.5 kb from the peak center.

**Supplementary Figure 13.**
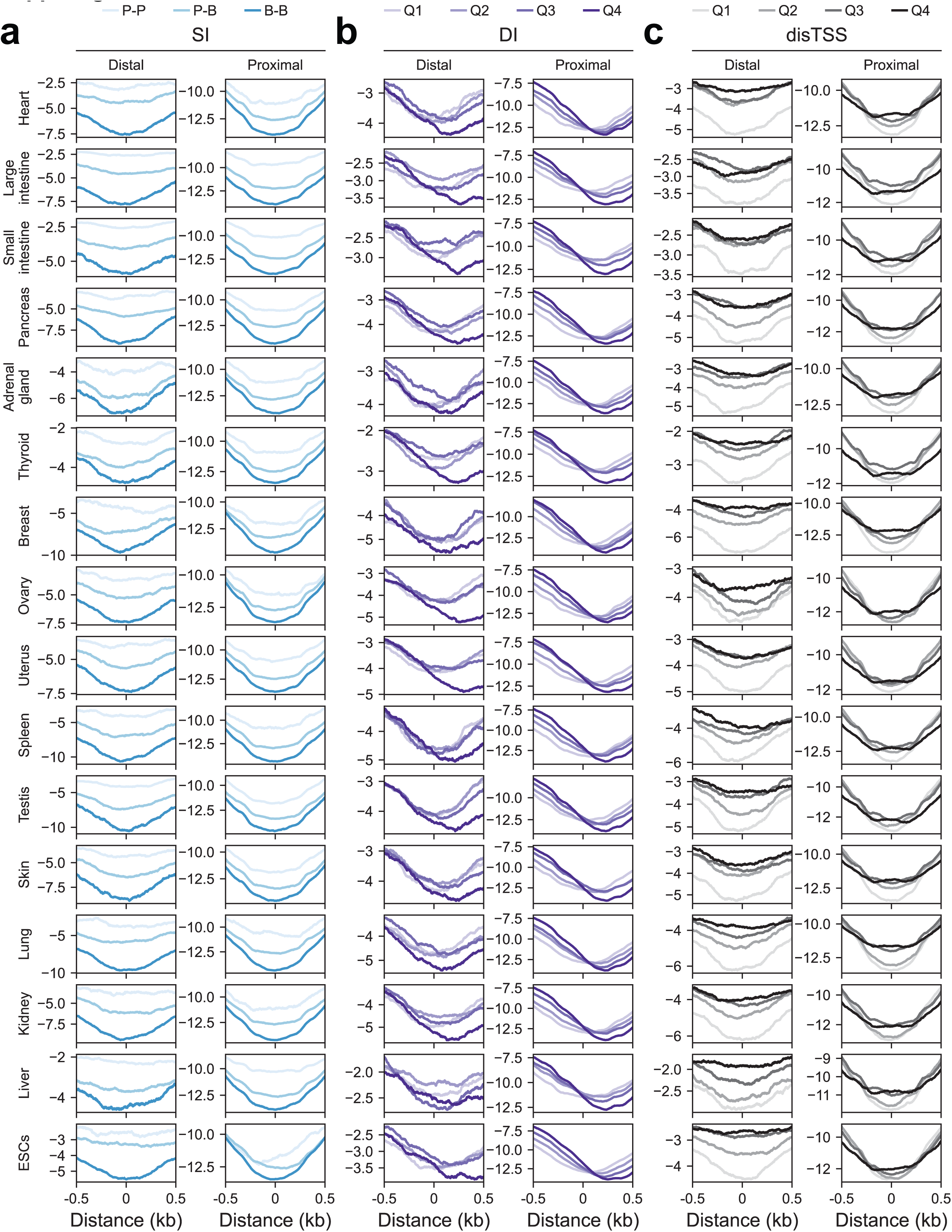
CDTS-based evolutionary constraint of TREs with distinct initiation features across tissue and cell types. Metaplots of CDTS across divergent distal and proximal elements, grouped by SI (**a**), DI (**b**), and disTSS (**c**) categories. Distances are shown as ±0.5 kb from the peak center.

**Supplementary Figure 14.**
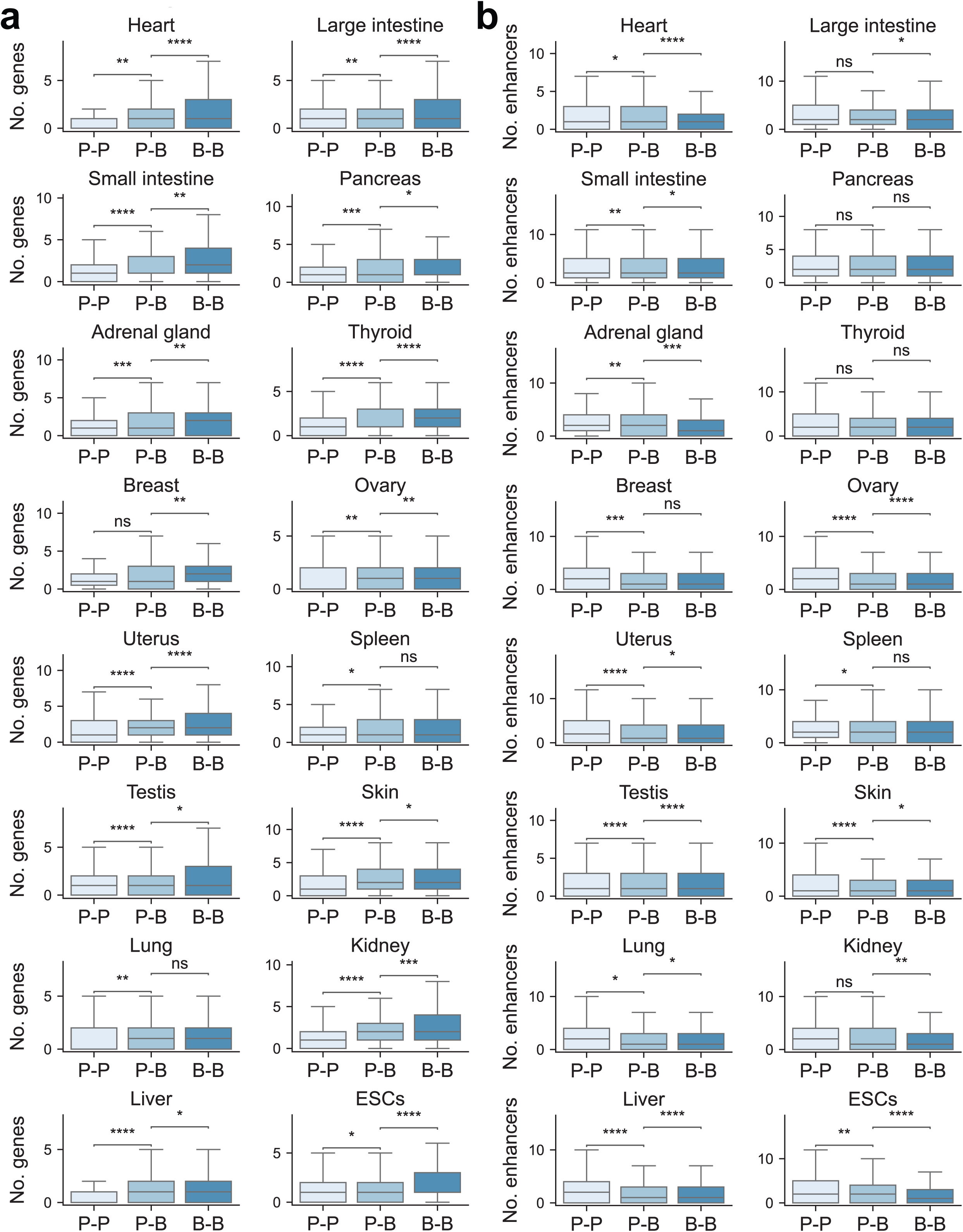
Relationship between TRE peak shape and regulatory connectivity across tissue and cell types. **(a)** Box plots showing the number of predicted target genes for distal TREs with different peak shapes across tissues and cells. Each point represents the number of target genes linked to a given TRE in the indicated biosample. **(b)** Box plots showing the number of enhancers linked to genes with proximal TREs of different peak shapes across tissues and cells. Each point represents the number of linked enhancers for a given gene in the indicated biosample. ns, non-significant; *, 0.01 < padj ≤ 0.05; **, 0.001 < padj ≤ 0.01; ***, 0.0001 < padj ≤ 0.001; ****, padj ≤ 0.0001.

**Supplementary Figure 15.**
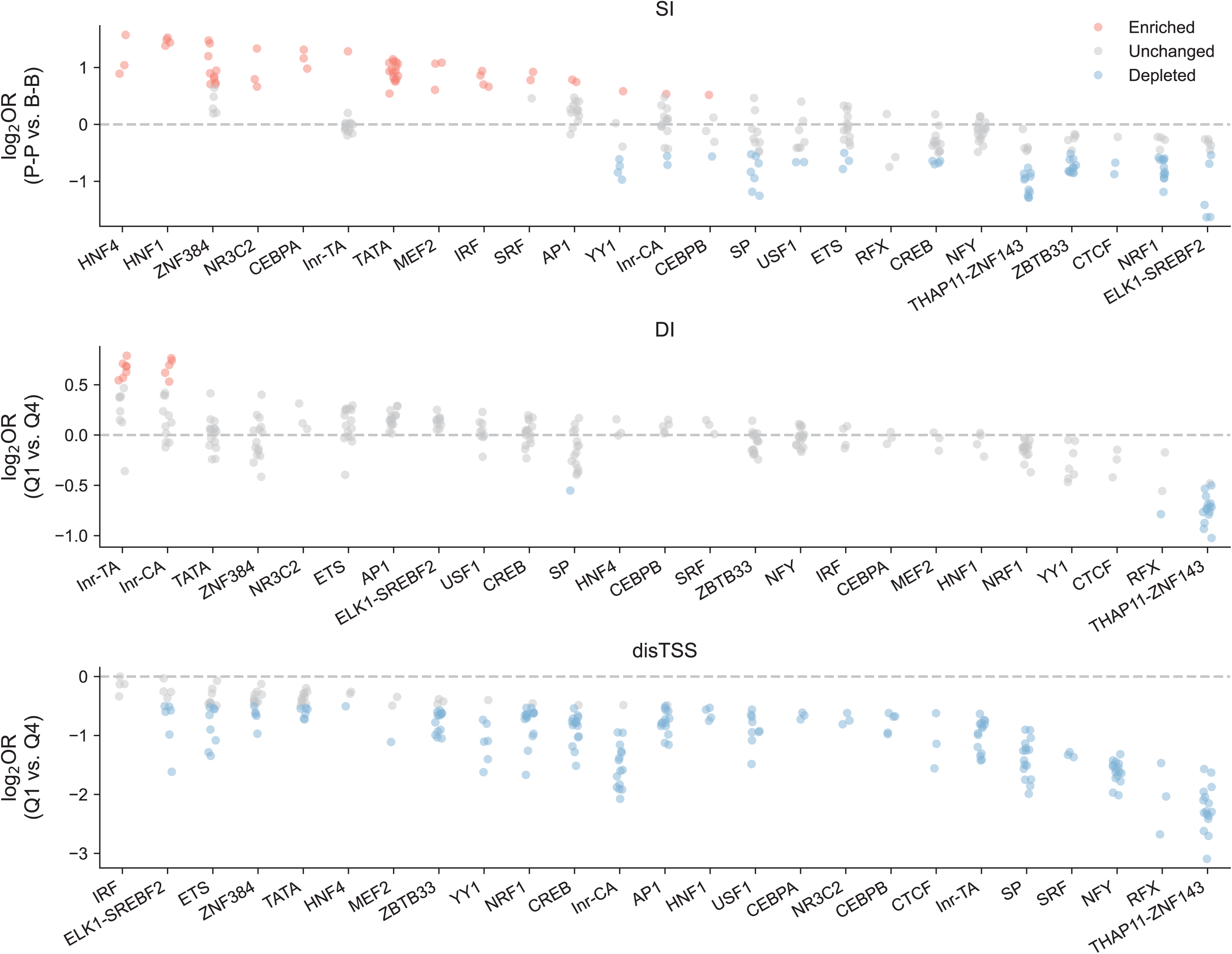
Motif enrichment across TREs with distinct initiation features. Complete panel corresponding to Figure 5E. Strip plot showing log_2_ odds ratios of motifs across TREs stratified by transcription initiation features (P-P vs B-B for SI; Q1 vs Q4 for DI and disTSS). Each point represents one tissue type. Colors indicate enrichment (red), depletion (blue), or no change (grey). Only motifs found in at least three tissue types are shown.

**Supplementary Figure 16.**
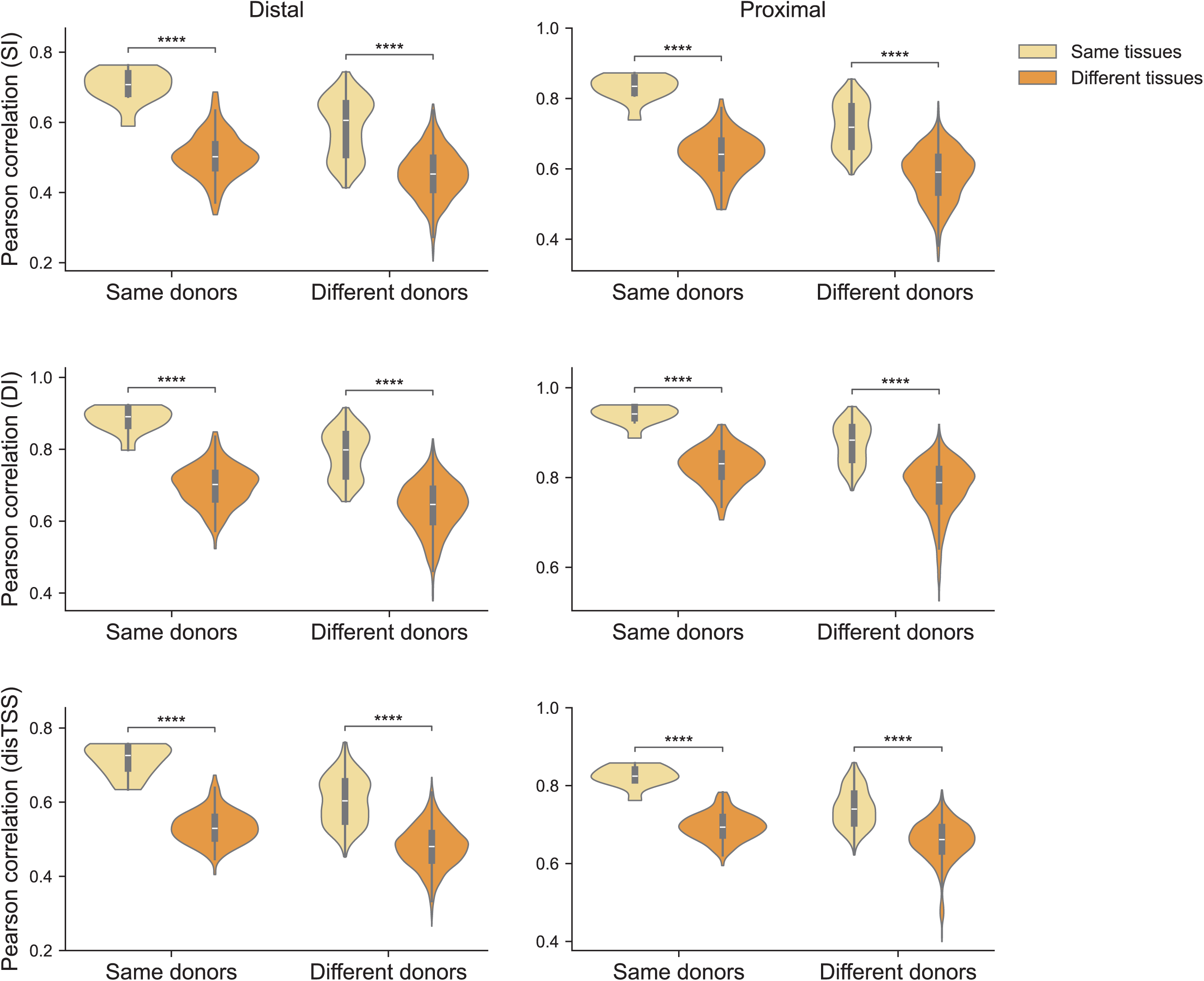
Intra- and inter-tissue variation in transcription initiation profiles. Violin plots showing Pearson correlations of SI, DI, and disTSS between shared distal and proximal TREs across samples of the same or different tissue types and donors. Each point represents the correlation for a given pair of samples. ****, padj ≤ 0.0001.

**Supplementary Figure 17.**
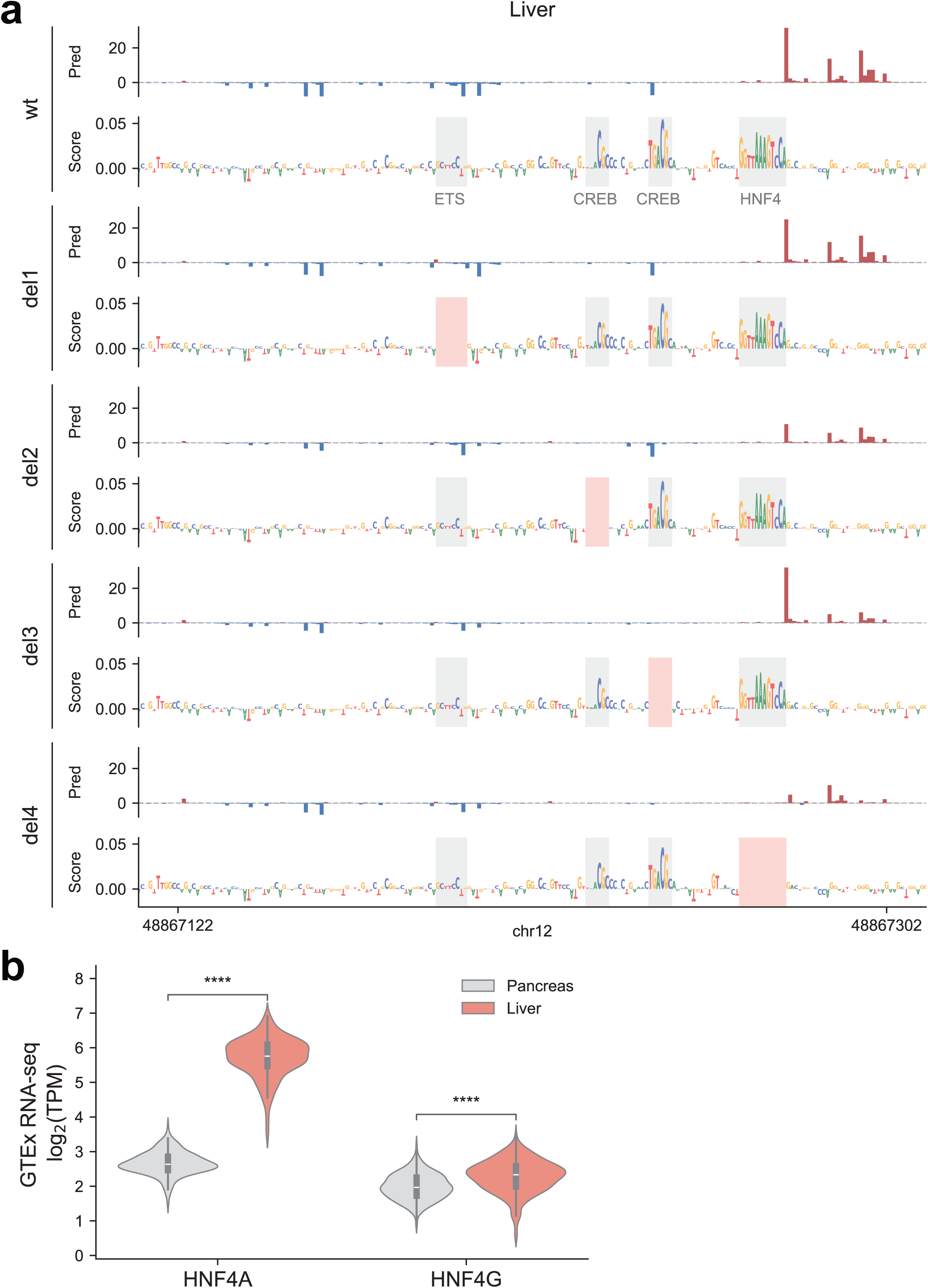
Inter-tissue effects of HNF4 motifs on transcription initiation profiles. **(a)** Predicted transcription profiles and contribution scores (count task) for the locus shown in Figure 5G, generated using a liver-trained ProCapNet model. The top track shows the wild type, with subsequent tracks showing the corresponding outputs following in silico deletion of the motif instance labeled in red. **(b)** Expression levels (log_2_TPM) of HNF4A and HNF4G in pancreas and liver tissues from GTEx RNA-seq data.

**Supplementary Figure 18.**
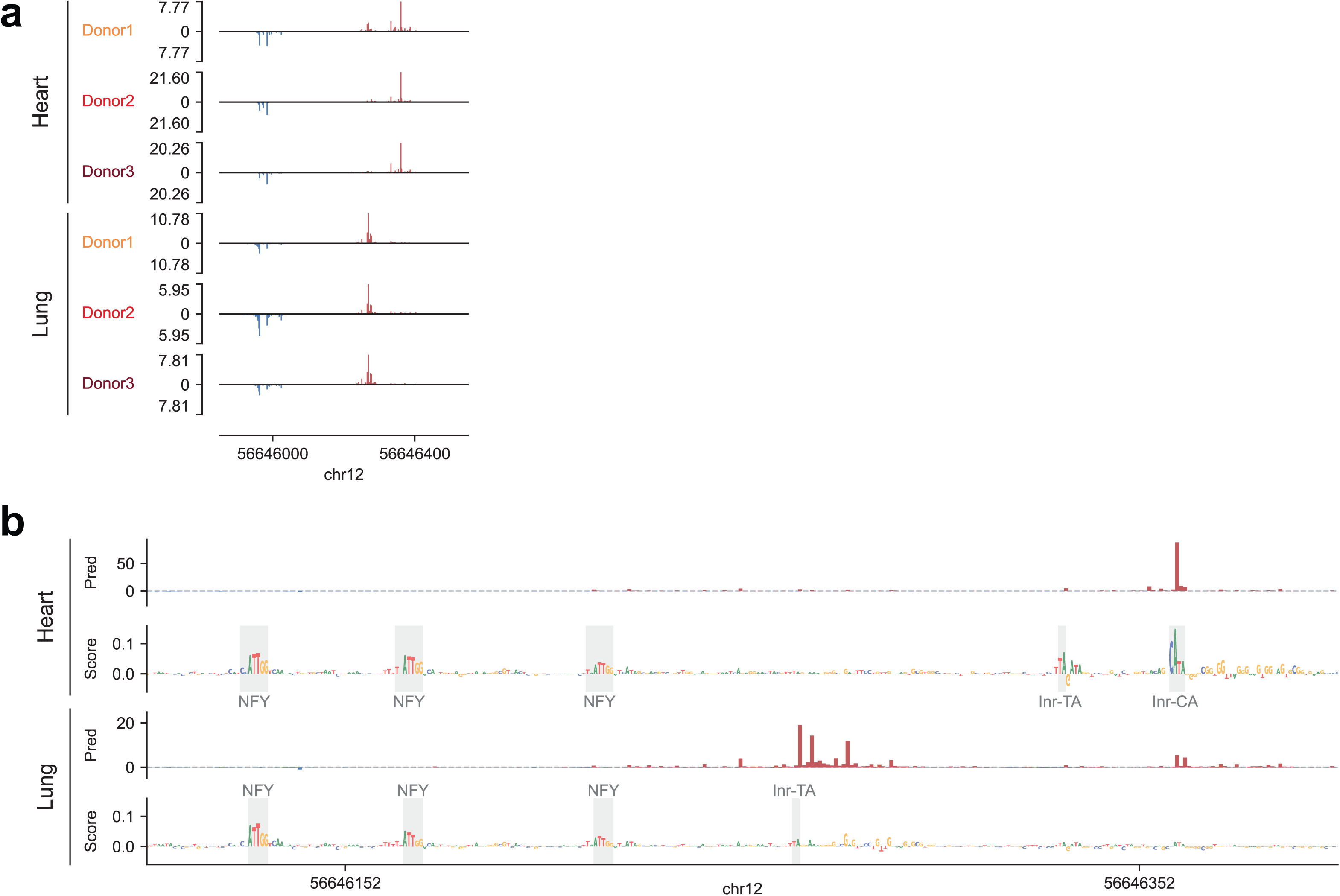
Tissue-specific preference for Initiator sequences shapes transcription initiation profiles. **(a)** Representative browser tracks of 5’ PRO-cap signals from two tissue types (heart and lung) of three matched donors across one distal genomic locus. **(b)** Predicted transcription profiles and contribution scores (profile task) for the locus shown in (a), generated using models trained on heart (top) and lung (bottom) PRO-cap data. Motifs contributing to transcription in each tissue type are labeled accordingly.

**Supplementary Figure 19.**
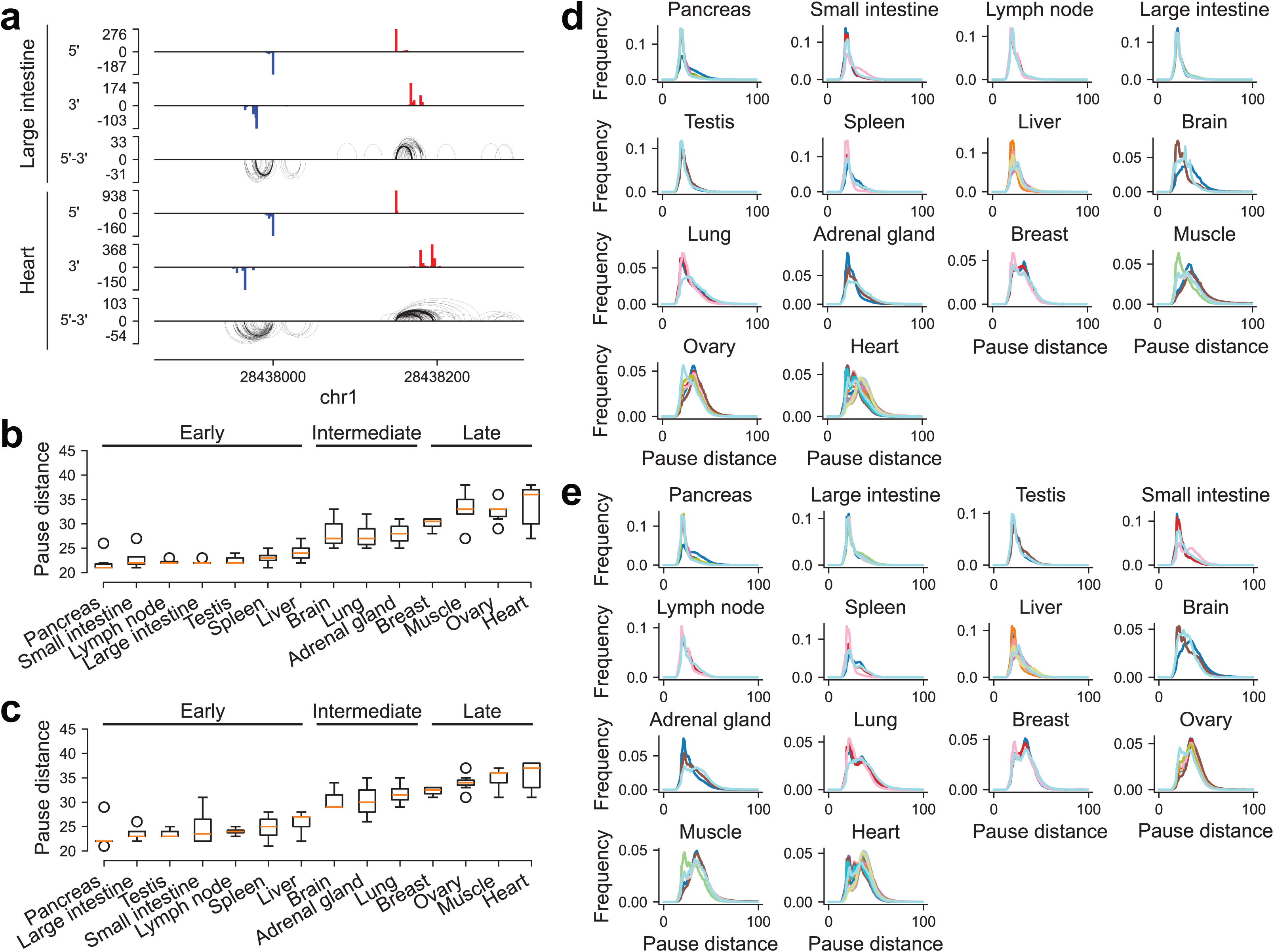
Variation of RNAPII pause distances of transcriptional regulatory elements across tissues. **(a)** Representative 5’ and 3’ PRO-cap signal tracks and pause distances for a divergent element in the large intestine (top) and heart (bottom). Loops denote 5′-3′ pause distances of individual nascent RNA molecules. **(b)** Boxplot summarizing median pause distances (bp) across tissues, grouped into early, intermediate, and late pausing categories, using reads mapped to distal elements. **(c)** Same as (b), but for proximal elements. **(d)** Distribution of pause distances (bp) across tissues, with each line representing one sample of a given tissue type, using reads mapped to distal elements. **(e)** Same as (d), but for proximal elements.

**Supplementary Figure 20.**
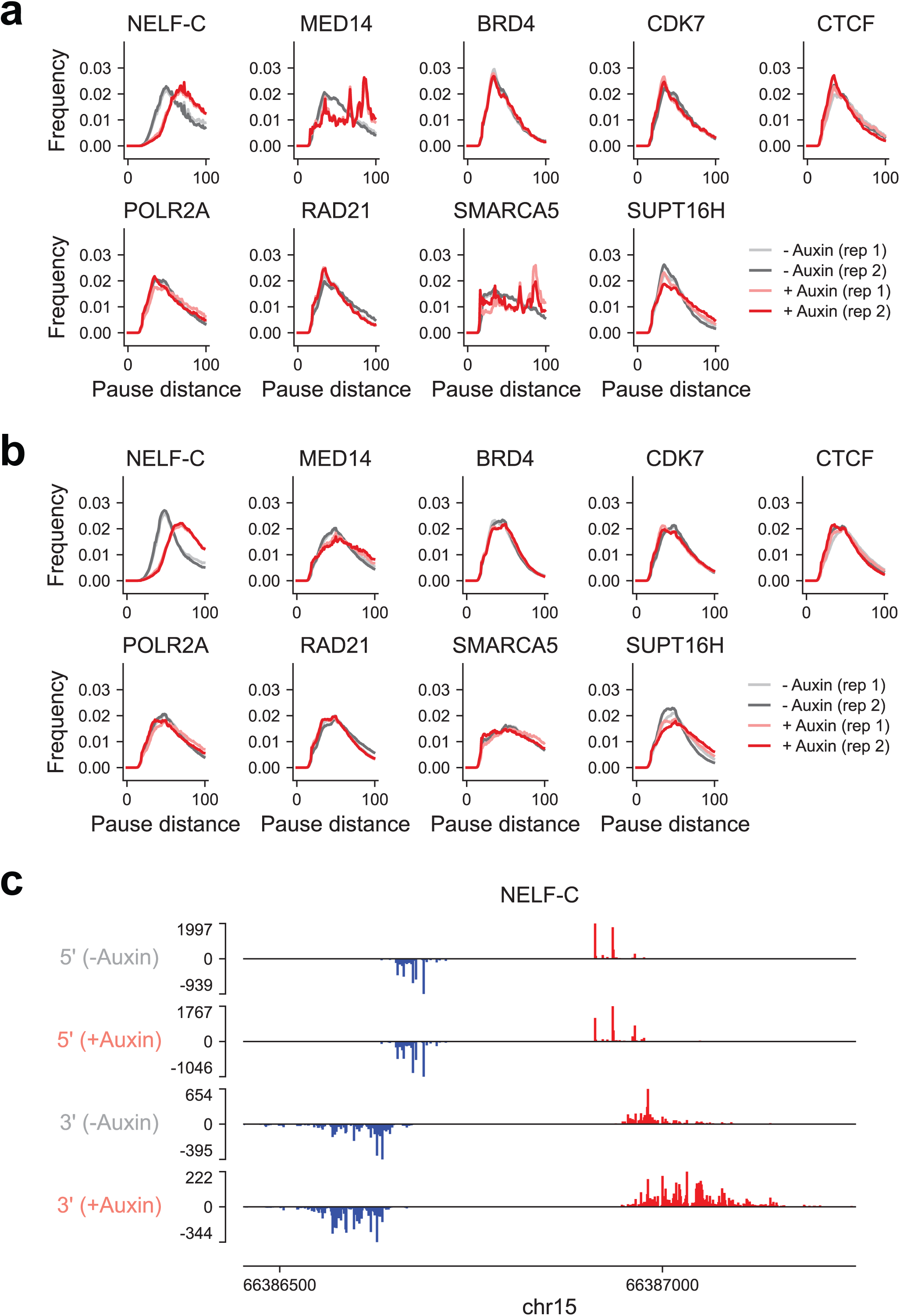
Genome-wide distribution of pause distances in transcriptional regulatory elements following depletion of different factors. **(a)** Genome-wide distribution of pause distances for distal elements before and after acute depletion of the indicated factors using auxin-inducible degron systems. All degron-tagged experiments were performed in HCT116 cells, except for NELF-C, which was generated in DLD-1 cells. The NELF-C degron-tagged dataset was obtained from a previously published study (GSE144786) and reprocessed using our pipeline. Two biological replicates were included per condition (−auxin and +auxin). **(b)** Same as (a), but for proximal elements. **(c)** Representative 5’ and 3’ PRO-cap signal tracks (merged across replicates) of an element before and after NELF-C degradation in DLD-1 cells.

**Supplementary Figure 21.**
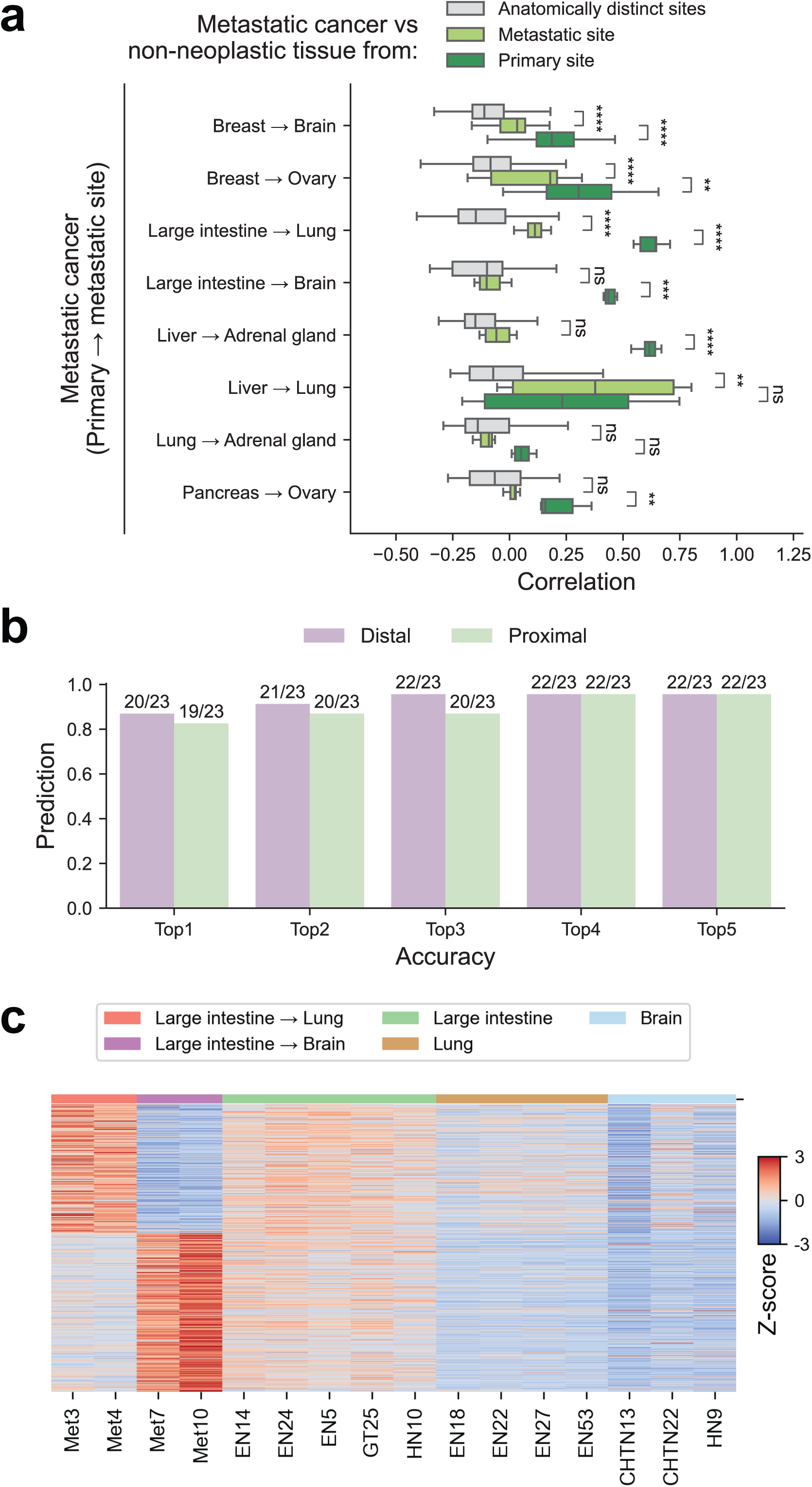
Transcriptional regulatory element signatures predict the tissue of origin in patient-derived tumors. **(a)** Pairwise comparisons of divergent proximal TRE expression profiles between metastatic tumors (n=23) and non-neoplastic tissues from the primary (dark green), metastatic (light green), and unrelated sites (gray). ns, non-significant; **, 0.001 < padj ≤ 0.01; ***, 0.0001 < padj ≤ 0.001; ****, padj ≤ 0.0001. **(b)** Accuracy of tissue-of-origin prediction for metastatic tumors using models trained on divergent distal and proximal PRO-cap profiles. Results are shown for top-1 through top-5 ranked predictions. **(c)** Heatmap of differentially expressed divergent distal TREs (n=1,128) between lung and brain metastases. Z-scores of PRO-cap signals were calculated across metastatic samples and related non-neoplastic tissues.

## Supplementary Table legends

**Supplementary Table 1 | Metadata of PRO-cap libraries**

Summary of biosample, experimental, and sequencing details for all libraries generated using PRO-cap. PBMC: peripheral blood mononuclear cell; T1D: type 1 diabetes; iPSC: induced pluripotent stem cell; ESC: embryonic stem cell; CML: chronic myelogenous leukemia; B-ALL: B-cell acute lymphocytic leukemia; GCB-DLBCL: germinal center B-cell diffuse large B-cell lymphoma; MCL: mantle cell lymphoma.

**Supplementary Table 2 | Motif enrichment across different TRE sets** Summary of motif enrichment and associated statistics for three TRE set comparisons: (1) TRE modules 1-10 from a leukemia and lymphoma cell line panel (LL-M1 to LL-M10); (2) T cells from T1D patients versus healthy donors (T1D-Up and T1D-Down); and (3) CRC metastases to lung versus brain (CRC Met-Brain and CRC Met-Lung). De novo motif enrichment is reported for the leukemia and lymphoma panel, while known motif enrichment is reported for the T1D and CRC metastasis comparisons.

**Supplementary Table 3 | Datasets and analyses across different annotations**

Details of datasets and analyses across annotations. The table is organized into four sheets: (1) data sources for tissue-matched comparisons across assays in Fig. 2a; (2) the total number of peaks for each annotation, along with the number and fraction of overlapping peaks with PRO-cap elements, reported for both PRO-cap–anchored and other annotation-anchored comparisons (irrespective of genomic location), for each matched tissue type, based on the datasets described in Sheet 1; (3) ProCapNet model training datasets; and (4) E2G prediction of promoter–enhancer (P-E) link datasets.

**Supplementary Table 4 | Primer sequences**

Details of cloning primers and TRE sequences used in the section “Tissue-specific effects of disease- and trait-associated variants.”

